# YAP-Driven Oral Epithelial Stem Cell Malignant Reprogramming at Single Cell Resolution

**DOI:** 10.1101/2023.07.24.550427

**Authors:** Farhoud Faraji, Sydney I. Ramirez, Lauren M. Clubb, Kuniaki Sato, Valeria Burghi, Thomas S. Hoang, Adam Officer, Paola Y. Anguiano Quiroz, William M.G. Galloway, Zbigniew Mikulski, Kate Medetgul-Ernar, Pauline Marangoni, Kyle B. Jones, Alfredo A. Molinolo, Kenneth Kim, Kanako Sakaguchi, Joseph A. Califano, Quinton Smith, Alon Goren, Ophir D. Klein, Pablo Tamayo, J. Silvio Gutkind

**Affiliations:** Department of Otolaryngology-Head and Neck Surgery, University of California San Diego Health; La Jolla, California, 92037; United States; Gleiberman Head and Neck Cancer Center, Moores Cancer Center, University of California San Diego Health; La Jolla, California, 92037; United States; Division of Infectious Diseases and Global Public Health, Department of Medicine, University of California San Diego Health; La Jolla, California, 92093; United States; La Jolla Institute for Immunology; La Jolla, California, 92037; United States; University of California San Diego, Biomedical Sciences Graduate Program; La Jolla, California, 92093; USA; Department of Pharmacology, University of California San Diego, School of Medicine, La Jolla, California, 92093; United States; University of California San Diego, Bioinformatics and Systems Biology Graduate Program; La Jolla, California, 92093; USA; Department of Chemical and Biomolecular Engineering, University of California Irvine; Irvine, California, 92697; United States; Department of Orofacial Sciences and Program in Craniofacial Biology, University of California San Francisco; San Francisco, California, 94143; United States; IDEXX Laboratories KK, Tokyo, 168-0063; Japan; Sue and Bill Gross Stem Cell Research Center, University of California Irvine, Irvine, California, 92697; United States; Department of Pediatrics, Cedars-Sinai Guerin Children’s, Los Angeles, California, 90048; United States; Division of Medical Genetics, Department of Medicine, University of California San Diego, California, 92093; United States; Center for Novel Therapeutics, University of California San Diego, La Jolla, California, 92037; United States

**Keywords:** Tumor initiation, Tumor initiating cell, Cancer stem cell, Squamous cell carcinoma, Oral cancer, HNSCC, Hippo pathway, YAP, HPV

## Abstract

Tumor initiation represents the first step in tumorigenesis during which normal progenitor cells undergo cell fate transition to cancer. Capturing this process as it occurs *in vivo*, however, remains elusive. Here we employ spatiotemporally controlled oncogene activation and tumor suppressor inhibition together with multiomics to unveil the processes underlying oral epithelial progenitor cell reprogramming into tumor initiating cells (TIC) at single cell resolution. TIC displayed a distinct stem-like state, defined by aberrant proliferative, hypoxic, squamous differentiation, and partial epithelial to mesenchymal (pEMT) invasive gene programs. YAP-mediated TIC programs included the activation of oncogenic transcriptional networks and mTOR signaling, and the recruitment of myeloid cells to the invasive front contributing to tumor infiltration. TIC transcriptional programs are conserved in human head and neck cancer and associated with poor patient survival. These findings illuminate processes underlying cancer initiation at single cell resolution, and identify candidate targets for early cancer detection and prevention.

## INTRODUCTION

Current models of carcinogenesis posit that tumor initiation requires oncogene activation^1^ concomitant with inactivation of intrinsic tumor suppressive mechanisms, including terminal differentiation,^2^ oncogene-induced senescence,^3^ and apoptosis.^4^ These insights are supported by recent genome-wide sequencing efforts that have cataloged candidate genomic alterations underlying most human malignancies.^5^ However, these studies in established, often advanced tumors are confounded by cellular and mutational heterogeneity and thus cannot directly identify tumor initiating cells (TIC) or discriminate between alterations driving tumor initiation from those promoting tumor progression. As such, the underlying molecular mechanisms mediating malignant reprogramming of normal progenitor cells into tumor initiating cells remains poorly understood.

Head and neck squamous cell carcinoma (HNSC) represents the most common malignancy arising from the upper aerodigestive epithelia.^6^ Extensive molecular characterization of HNSC has revealed that alterations in numerous genes in a given tumor converge to impact a finite set of oncogenic molecular pathways.^7^ HNSC is characterized by near universal loss-of-function of *TP53* and *CDKN2A* tumor suppressors by genomic alteration or human papillomavirus (HPV) E6 and E7 oncoprotein-mediated inhibition.^7,8^ Notably, in prior studies we reported that alterations in *FAT1*, observed in nearly one third of HNSC,^7^ disrupt Hippo pathway signaling and result in unrestrained activation of the transcriptional co-activator YAP.^9^ Furthermore, beyond *FAT1* mutation, several other genomic alterations observed in HNSC have been associated with Hippo pathway disruption and YAP activation.^10^ Yet the direct effects of unrestrained YAP activation on tumor initiation are unknown.

The cell of origin of HNSC, oral tumor initiating cells, and mechanisms of HNSC initiation remain poorly understood.^11^ Self-renewing oral epithelial progenitor cells (OEPCs) reside in the basal layer of the stratified squamous epithelium.^12^ These cells contribute to long-term epithelial maintenance and give rise to different cell types that form tongue and soft palate epithelia.^13,14^ As such, OEPCs may represent the cell of origin for HNSC, and render the oral epithelium an ideal system to elucidate early molecular events underlying malignant reprogramming.^15^ In the present study, we combine knowledge of the landscape of oncogenic pathway alterations in HNSC with genetically engineered animal models, lineage tracing, and multiomics to unveil the underpinnings of cancer initiation *in vivo*.

## RESULTS

### YAP activation and E6-E7 expression in OEPCs is sufficient to induce rapid tumor initiation

Tumor initiation represents the first step in tumorigenesis during which normal progenitor cells undergo cell fate transition to cancer. To investigate this process, we developed genetically engineered murine systems focusing on prevalent and co-occuring genomic alterations in HNSC. While genomic alterations involving *FAT1* are observed in ~30% of HNSC, this may represent one of multiple mechanisms promoting YAP activation.^16^ YAP activation may also occur through amplification of *YAP1* or the YAP paralog TAZ (*WWTR1*), indicating that YAP activation is observed with even higher frequency in HNSC.^10,16–18^ We performed immunohistochemical (IHC) staining of YAP in human tissue microarrays, using nuclear localization as a surrogate for YAP activation.^19^ Consistent with its physiologic role in stem cell maintenance,^9,20^ nuclear YAP was detected primarily in basal cells in normal oral epithelial tissue. Conversely, YAP activated cells were distributed throughout tumor tissue in the majority of HPV- and HPV+ HNSC lesions (**Fig. 1a-c, Extended Data Fig. 1a-c**).

**Figure 1.**
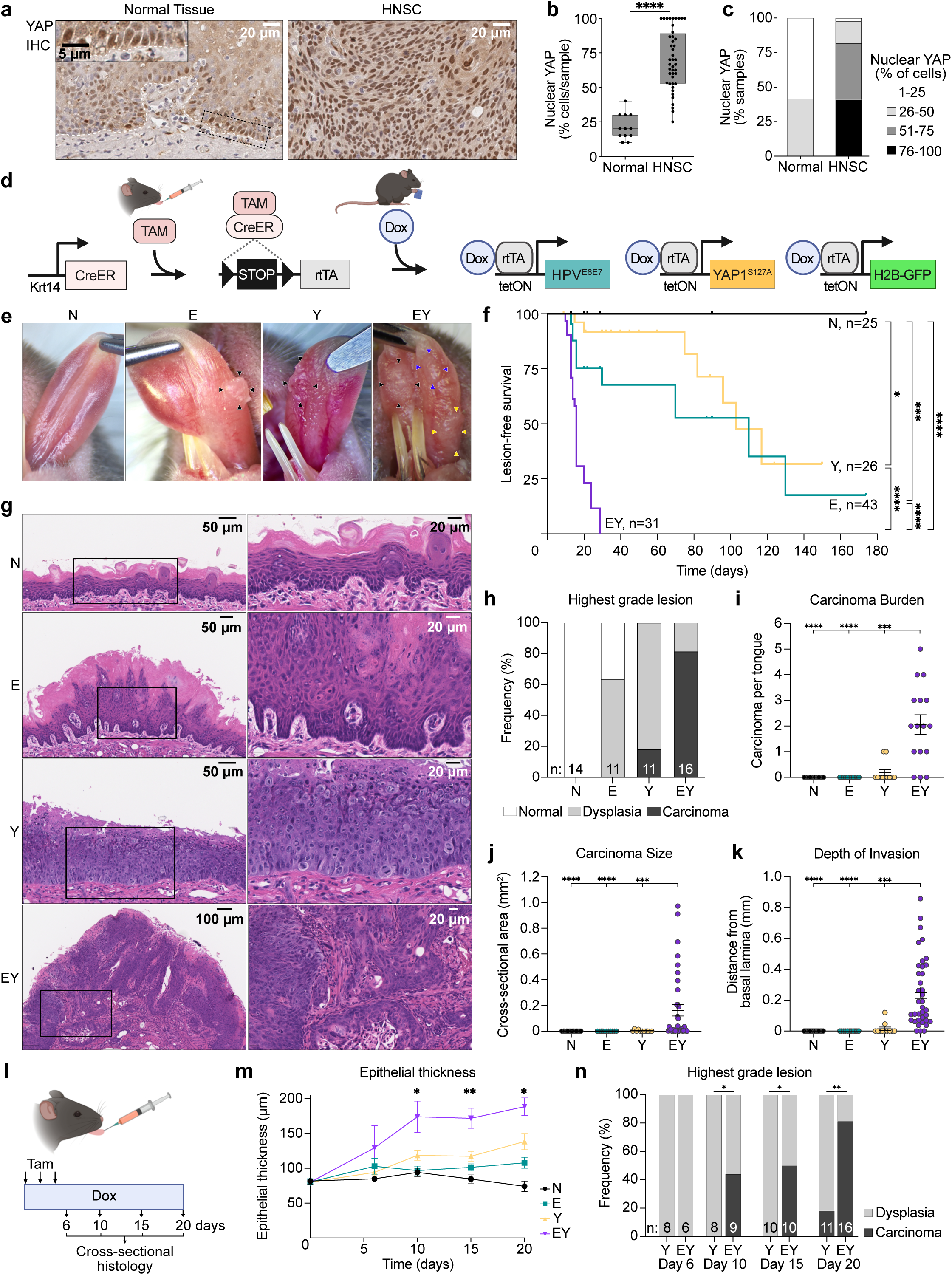
YAP and E6-E7 activation is sufficient to induce rapid tumor initiation in OEPCs. (a) Representative images of nuclear YAP protein staining in tissues by IHC in normal human oral epithelial tissue (left) and malignant human HNSC tissue (right). (b) Percent of cells with nuclear YAP protein by IHC in normal human tissue (n=12) and malignant HNSC (n=44) from tissue microarrays. (c) Percent of samples with positive nuclear YAP IHC staining in normal human tissues and malignant HNSC from the same tissue microarray samples as Panel b. (d) Schematic depicting spatiotemporally controlled activation of YAP and HPV oncogenic pathways in oral epithelial progenitor cells. A tissue-specific inducible Cre recombinase (CreERT) expressed in *Krt14*-expressing cells is activated by intralingual injection of tamoxifen (Tam), resulting in recombination of the Lox_p_-STOP-Lox_p_ cassette (LSL) and expression of the *rtTA* transgene. Feeding tamoxifen-treated mice doxycycline chow (Dox) activates transcription of the tetracycline-inducible transgenes *HPV16^E6-E7^*, *YAP^S127A^*, and *H2B-GFP* reporter. (e) Representative images of gross tongue lesions 20 days after transgene induction. (f) Kaplan-Meier plot showing the kinetics of tongue lesion formation upon transgene induction. Bonferroni-corrected Mantel-Cox log-rank test. (g) Hematoxylin and eosin (H&E) stained tongue sections demonstrating epithelial changes 20 days after transgene activation. (h) Histopathologic evaluation and scoring of mouse tongue epithelia; n indicates number of tongues examined for each transgenic condition. Dysplasia: any dysplastic changes present. Carcinoma: infiltrative lesions invading beyond the basement membrane. (i) Number of infiltrative carcinoma lesions per examined tongue. (j) Cross-sectional area of infiltrative carcinoma lesions. (k) Depth of carcinoma invasion as measured by the longest plumb line orthogonal to the tangent of the nearest intact basement membrane. (l) Schematic showing pulse-chase strategy of transgene induction and time points for histological analysis of tongue epithelia. (m) Longitudinal measurement of tongue epithelial thickness. n=3-7 mice per condition per time point. ANOVA with Tukey correction for multiple comparisons; means with standard errors of the mean (SEM) are shown. (n) Highest grade lesion identified per tongue for the carcinogenic conditions Y and EY. Two-sided Fisher’s exact test. Panels i-k: n_Y_=2, n_EY_=34 carcinomata. ANOVA with Tukey correction for multiple comparisons; means with SEM are shown. For all panels with asterisks denoting significance: *p<0.05, **p<0.01, ***p<0.001, ****p<0.0001.

To investigate tumorigenesis in the context of a minimum complement of pathway alterations, we employed transgenic expression of the HPV16 E6-E7 oncogene, which inhibits *TP53* and *CDKN2A* tumor suppressors,^21^ and the constitutively active *YAP1^S127A^* allele.^22^ The latter enables direct YAP activation rather than through *FAT1* gene disruption, as *FAT1* exerts other functions, including activation of Wnt^23^ and CDK6 signaling,^24^ the EGFR/ERK axis,^25^ the CAMK2/CD44/SRC axis,^26^ and recruits the E3 ligase MIB2,^27^ whose disruption may have confounded the emerging results. Keratin 14 (KRT14) is expressed in the basal layer of oral epithelia, which contains OEPCs that may represent the cell of origin for HNSC.^28^ Utilizing a tamoxifen-inducible Cre-recombinase (CreERT) driven by the *Krt14* promoter, genomic alterations were targeted to KRT14^+^ OEPCs.^29^ We bred mice bearing *Krt14-CreERT* and *LSL-rtTA* regulatory transgenes, an *H2B-GFP* reporter, and E6-E7 (“E”), *YAP1^S127A^* (“Y”), or both transgenes (“EY”). Littermates lacking the E6-E7 or *YAP1^S127A^* alleles but possessing regulatory and reporter transgenes served as normal controls (“N”). Intralingual administration of tamoxifen activated CreERT-mediated recombination of a floxed STOP cassette (LSL) and enabled transcription of the reverse tetracycline-controlled transactivator *(rtTA)* in KRT14^+^ OEPCs. Administration of doxycycline chow then induced expression of the tetracycline response element-regulated *HPV16^E6-E7^*, *YAP1^S127A^*, and *H2B-GFP* transgenes (**Fig. 1d, Extended Data Fig. 1d-g**).^21,22,30^

Longitudinal examination of mouse tongues identified macroscopic lesions as early as 8 days after transgene induction in EY mice. By 20 days, the majority (65%) of EY mice bore at least two lesions, while few E or Y mice, and no N mice bore any gross lesions (**Fig. 1e,f, Extended Data Fig. 1h**). Histopathology showed invasive carcinoma in 81% of EY mice, compared to 18% of Y mice, and no E or N mice (**Fig. 1g-h**, **Extended Data Fig. 1i**). Most carcinoma-bearing EY mice had multiple independent carcinomas, which were more abundant, larger, and more deeply invasive than carcinoma in Y mice (**Fig. 1i-k**). We next investigated tumor initiation at higher temporal resolution using a pulse-chase strategy (**Fig. 1l**). At day 10, we observed a marked increase in EY epithelial thickness and invasive carcinoma occurred in 44% of EY mice. No carcinoma in Y epithelia were observed until day 20 (**Fig. 1m-n**). These findings suggest that unrestrained YAP activation in the context of E6-E7 expression in KRT14^+^ OEPCs is sufficient to induce oral carcinoma with high penetrance and rapid kinetics.

To test tumor initiating capacity, we orthotopically implanted cell suspensions generated from E, Y, and EY transgene-induced epithelia into NOD-SCID-gamma (NSG) mice. EY-induced cells formed large tumors in all mice, and Y-induced cells formed small tumors in a few mice, demonstrating the tumorigenic potential of these YAP-activated cells (**Extended Data Fig. 2a-e**). Primary N and EY cell cultures were then developed to further investigate the roles of *YAP^S127A^* and *HPV^E6E7^* transgene expression in tumor initiation. H2B-GFP positive cells were isolated from N and EY-induced lingual epithelia and subjected to fluorescence activated cell sorting (FACS) to enrich for transgene activated cells. FACS-sorted EY cells maintained *HPV16^E6-E7^* and *YAP1^S127A^* expression in culture, and displayed tumorigenicity upon implantation of as few as 5,000 cells (**Extended Data Fig. 2f-h**).

### YAP and E6-E7 activation induces cell cycle activation and loss of normal OEPC identity

To understand transcriptome-wide changes attributable to transgene expression, we performed bulk RNA sequencing (RNAseq) on microdissected tongue epithelia 15 days post-induction, because at this time point approximately half of EY mice were observed to have invasive carcinoma (**Extended Data Fig. 3a**). As expected, *E6* and *E7* were detected in E and EY epithelia. *YAP^S127A^* and YAP target gene^17^ upregulation were observed in Y and EY epithelia (**Extended Data Fig. 3b**). Transgene activation resulted in significant transcriptional differences across groups (**Extended Data Fig. 3c**). Venn analysis revealed 2,318 genes differentially expressed genes (DEGs) solely in EY epithelia (EY-unique DEGs, **Fig. 2a, Supplementary Tables 1,2**). GSEA of Molecular Signatures Database (MSigDB) Hallmark pathways^31,32^ and gene ontology (GO)^33,34^ identified enrichment among EY-unique DEGs for processes underlying cell proliferation, epithelial cell development and identity, and inflammatory responses (**Fig. 2b, Extended Data Fig. 3d**).

**Figure 2.**
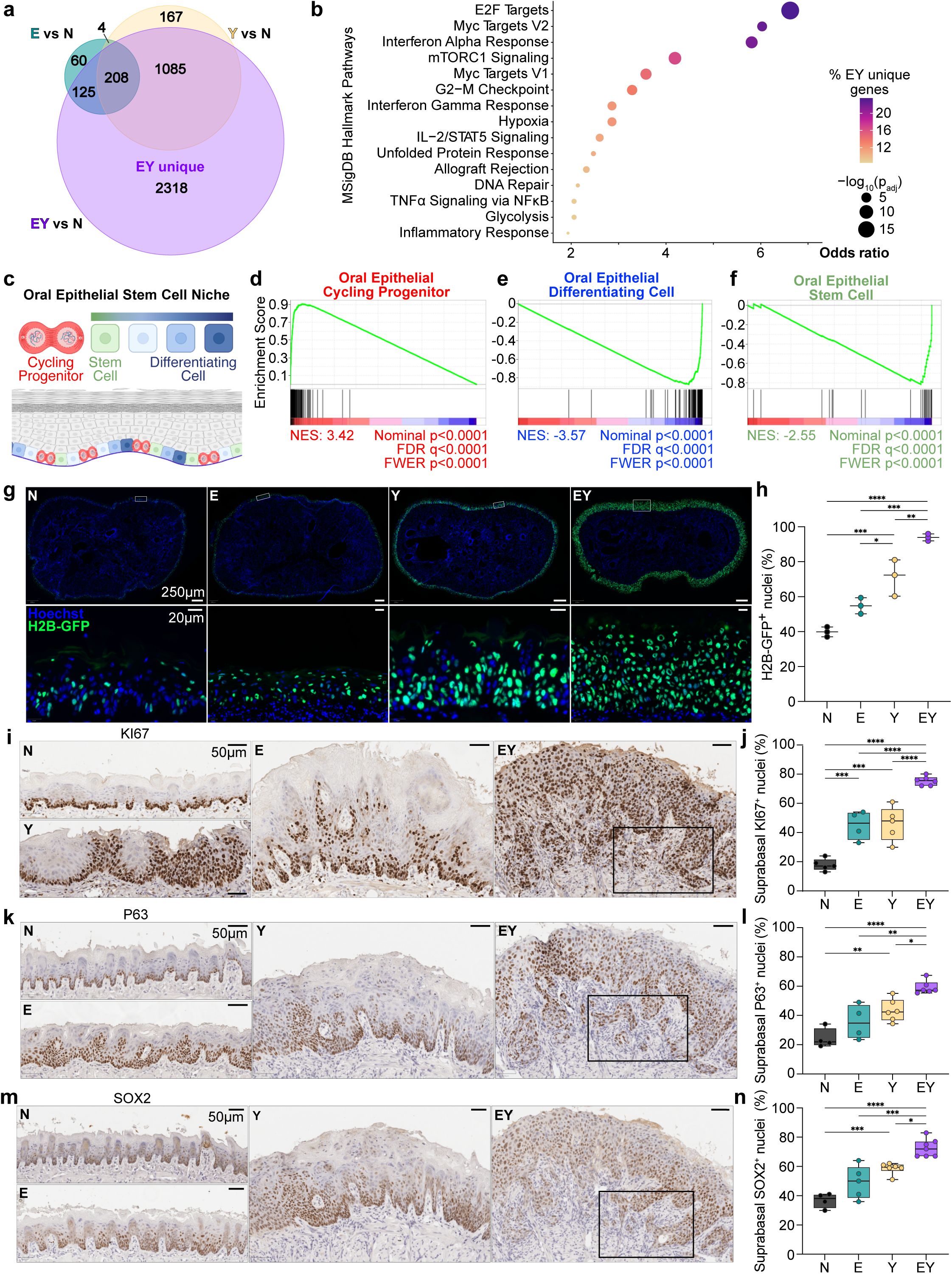
Combined YAP and E6-E7 activation drives loss of normal OEPC identity. (a) Venn analysis of shared and unique differentially expressed genes for each transgenic condition versus control (N). ‘EY unique’ denotes the 2,318 genes uniquely dysregulated in EY. Log_2_FC>1 and p_adj_<0.01. (b) MSigDB Hallmark Pathways enriched in upregulated EY-unique DEGs. (c) Model of the basal layer oral epithelial stem cell niche depicting physiologic cell states including stem cells, cycling progenitor cells, and differentiating cells. Adapted from Jones et al.^9^ (d-f) GSEA analysis enrichment plots of EY vs N mouse tongue epithelium DEGs for (d) cycling progenitor cell (G1/S, G2/M), (e) differentiating cell, and (f) stem cell oral epithelial cell programs. (g-h) Lineage tracing by fluorescent microscopy using the *H2B-GFP* reporter to track and quantify GFP^+^ nuclei. (g) Representative axial tongue sections of H2B-GFP^+^ nuclei 6 days after transgene induction. (h) Percent H2B-GFP^+^ nuclei in g. (i-n) Representative IHC images 20 days after transgene induction of (i) KI67^+^ nuclei, (k) P63^+^ nuclei, (m) SOX2^+^ nuclei. Percent of positively stained nuclei in the suprabasal epidermal strata for: (j) KI67, (l) P63, (n) SOX2; ANOVA with Tukey correction for multiple comparisons, *p<0.05, **p<0.01, ***p<0.001, ****p<0.0001.

Extending our transcriptional analysis to OEPC gene programs, we focused on previously described murine oral epithelial basal layer cell states defined as stem, cycling progenitor, and differentiating cells (**Fig. 2c, Supplementary Table 3**).^12^ Using signatures of these physiologic cell states, we observed pronounced enrichment of the cycling progenitor signature and dramatic depletion of the differentiation signature in EY epithelia (**Fig. 2d-f** and **Extended Data Fig. 3e**). At the gene level, we observed EY-mediated upregulation of OEPC stemness factors and downregulation of differentiation and apicobasal polarity factors. Several basal progenitor state factors, however, displayed paradoxical downregulation (**Extended Data Fig. 3f**). Notably, EY transcriptomes showed enrichment for epithelial to mesenchymal transition (EMT) signatures (**Extended Data Fig. 3g**). These findings indicate that EY-driven transcriptional changes do not reflect a physiologic OEPC state, but rather a unique transcriptional state related to tumor initiation.

We next performed lineage tracing of transgene-activated cells using the *H2B-GFP* reporter to evaluate if transgene activation drives cellular proliferation *in vivo*. Thirty-six hours after a single dose of tamoxifen, we observed a similar distribution of H2B-GFP^+^ basal cells across all groups (**Extended Data Fig. 4a-b**). By 6 days, H2B-GFP^+^ cells had expanded throughout the full thickness of EY epithelia (**Fig. 2g,h**). KI67 showed that while mitotically active cells were restricted to the basal layer in N epithelia, the expression of E, Y, or EY transgenes resulted in suprabasal extension of KI67^+^ cells (**Fig. 2i,j and Extended Data Fig. 4c,d**).

To examine the impact of transgene activation on cellular identity, we evaluated the expression of factors involved in OEPC identity maintenance: P63, SOX2, and ITGA6.^14,35–37^ In N epithelia, P63 and SOX2 expression was restricted to the basal layer, and ITGA6 to basal cells in contact with the basement membrane. YAP activation resulted in ectopic extension of P63^+^ cells into suprabasal layers, while EY expression induced the P63^+^ compartment to occupy the entire epithelial depth (**Fig. 2k,l, Extended Data Fig. 4e,f**). Similarly, EY expression drove expansion of SOX2^+^ cells into suprabasal layers (**Fig. 2m,n**). While E6-E7 expression alone did not alter ITGA6 compartmentalization, Y and EY expression diminished ITGA6 basement membrane localization and resulted in diffuse low (Y) to high (EY) ITGA6 expression throughout suprabasal strata (**Extended Data Fig. 4e,g**). Strikingly, most cells at the EY invasive front were positive for KI67^+^, TP63^+^, and SOX2^+^ (**Extended Data Fig. 4h-j**). Together, these findings support that concomittant YAP and E6-E7 activation induces rapid expansion of a highly proliferative stem cell-like population, which disrupt epithelial tissue architecture.

### YAP-driven epigenetic reprogramming promotes proliferation, invasion, and inflammation

To examine gene expression programs directly regulated by YAP, we performed transcriptional profiling by RNAseq, mapped YAP genome-wide binding to native chromatin by CUT&Tag, evaluated promoter and enhancer activity by H3K27ac by CUT&Tag, and explored chromatin accessibility by ATACseq in N and EY primary cell cultures. Comparing transcriptomes, we observed high concordance (~90%) among EY DEGs across transgene-activated primary cultured cells and whole epithelia (**Extended Data Fig. 5a**). Transcripts detected exclusively in tissue included stromal and immune transcripts and signatures not expected to be observed in FACS-enriched cultured cells (**Extended Data Fig. 5b-c**).

YAP CUT&Tag identified 38,002 YAP binding sites (‘peaks’). Consistent with published YAP ChIPseq data, YAP CUT&Tag peak distribution showed that ~43% of YAP peaks occurred in intergenic regions, and approximately half of YAP peaks occurred 10-100kb from transcription start sites (TSS, **Extended Data Fig. 5d**).^38^ Compared to N, EY cells had 11,169 (29%) gained, 3109 (8%) lost, and 23,724 unchanged YAP peaks (**Fig. 3a**). Transcription factor motifs enriched at YAP gained peaks in EY included TEAD, AP-1, Sp2, KLF, p63, and NRF2, suggesting that these factors may cooperatively regulate transcription with YAP (**Extended Data Fig. 5e**). Indeed, TEAD family DNA binding proteins are required for YAP-mediated gene expression,^39^ and YAP/TEAD complex with AP-1 factors, KLF4, and p63 to regulate transcriptional programs.^38,40,41^

**Figure 3.**
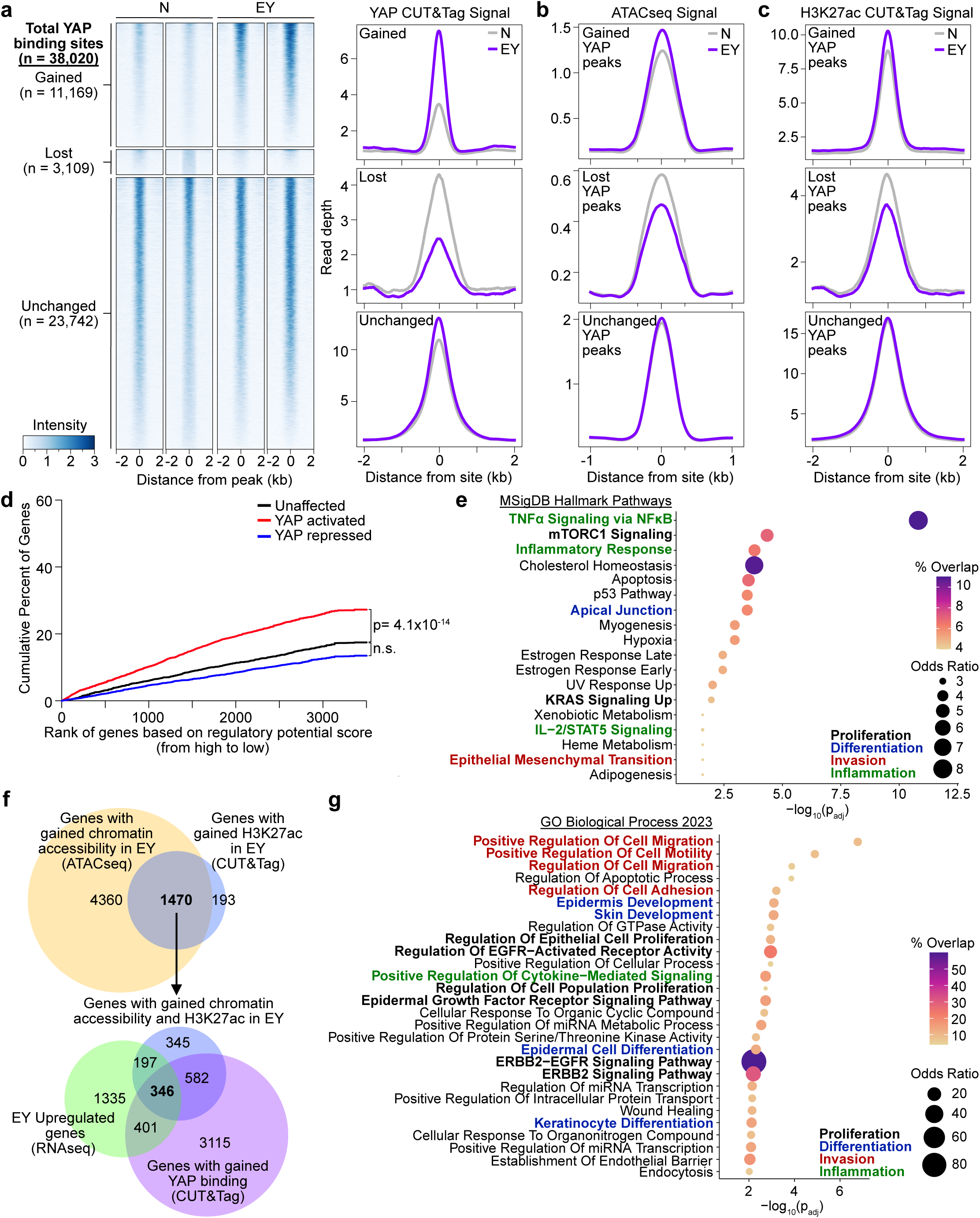
YAP-driven epigenetic reprogramming of oral epithelial progenitor cells. (a) Left: Heatmaps of YAP CUT&Tag signal in primary cell cultures from N and EY epithelia. YAP binding sites were characterized as gained, lost, and unchanged based on the comparison of YAP CUT&Tag results from EY and N cells. Sites were ranked from the strongest to weakest YAP binding for each category, shown in ±2-kb windows centered at YAP binding sites. At least two independent biological replicates were performed and two are shown for each condition. Right: Averaged YAP CUT&Tag signal centered at gained, lost, and unchanged YAP binding sites, as determined above, across EY and N. Averaged signals of at least two biological replicates are shown. (b) ATACseq signals at gained, lost, and unchanged YAP binding sites, as determined above. Averaged signals of at least four biological replicates are shown. (c) H3K27ac CUT&Tag signal centered at gained, lost, and unchanged YAP binding sites, as determined above. Averaged signals of at least two biological replicates are shown. (d) Binding and expression target analysis (BETA) of activating and repressive function of the gained YAP binding sites. The red, blue and black lines represent cumulative fractions of genes that are activated, repressed, or unaffected by YAP. P values were calculated by two-sided Kolmogorov–Smirnov tests. (e) MSigDB Hallmark Pathways enriched in the top 300 YAP-activated genes based on BETA. (f) Top Venn: Overlap of genes near EY-gained chromatin accessibility by ATACseq and EY-gained H3K27ac CUT&Tag peaks. Bottom Venn: Genes with EY-gained chromatin accessibility and H3K27ac peaks overlapped with EY-upregulated genes by RNAseq and genes near EY-gained YAP CUT&Tag peaks. (g) Gene Ontology biological processes enriched in EY-upregulated genes with EY-gained YAP CUT&Tag, H3K27ac, and ATACseq peaks.

We next evaluated functional chromatin elements co-localizing with YAP peaks. Globally, 28,986 (76%) of YAP CUT&Tag peaks overlapped with ATACseq peaks. Approximately 8% of EY gained YAP peaks overlapped with EY gained ATAC sites, indicating that EY expression resulted in the emergence of a subset of YAP binding sites associated with newly opened chromatin regions (**Fig. 3b**). Examining transcriptional regulatory activity at YAP binding sites, we observed a subtle increase in H3K27ac CUT&Tag signal intensity at gained YAP binding sites in EY (**Fig. 3c**). Together, these findings suggest that EY expression leads to increased chromatin accessibility and activation in a subset of YAP-regulated genes without exerting strong global effects on chromatin state.

To gain insight into specific genes and pathways regulated by YAP, RNAseq and YAP CUT&Tag data were integrated using Binding and Expression Target Analysis (BETA), showing that YAP predominantly acts as a transcriptional activator (**Fig. 3d, Supplementary Table 4**). Intriguingly, the top 200 YAP-activated genes showed marked enrichment for three MSigDB Hallmark Pathways: TNFa signaling, mTORC1 signaling, and inflammatory response (**Fig. 3e**), highlighting that mTOR signaling and immune cell related processes may both be important in EY tumor initiation.

Integrating H3K27ac CUT&Tag and ATAC, we found that 88% of genes with gained H3K27ac peaks also gained chromatin accessibility in EY cells. Integrating these genes with YAP CUT&Tag and RNAseq data delineated 346 genes with gained YAP binding, local promoter/enhancer activation by H3K27ac, chromatin accessibility, and transcriptional upregulation in EY cells (**Fig. 3f, Supplementary Table 4**). GO analysis of these genes showed strong enrichment for pathways associated with invasion, epithelial cell fate determination, proliferation, and ERB2/EGFR signaling (**Fig. 3g**). These findings raised the possibility that beyond activating transcriptional programs driving proliferation, invasion, and inflammation, YAP may promote mTOR and EGFR signaling, which are among the most frequently activated signaling mechanisms in HNSC.^42,43^

### Oncogenic transcriptional programs define tumor initiating cells

Our bulk transcriptional analyses shed insight into the processes occurring in transgene activated epithelial tissue undergoing malignant conversion. To identify transcriptional programs in nascent tumor initiating cells we performed single cell RNAseq (scRNAseq) of oral epithelia at the same time point as RNAseq. In total, 12,771 epithelial cells were identified across 8 clusters, which could broadly be divided into physiologic^12^ and transgene-associated cell states (**Fig. 4a-d and Extended Data Fig. 6a-b**).

**Figure 4.**
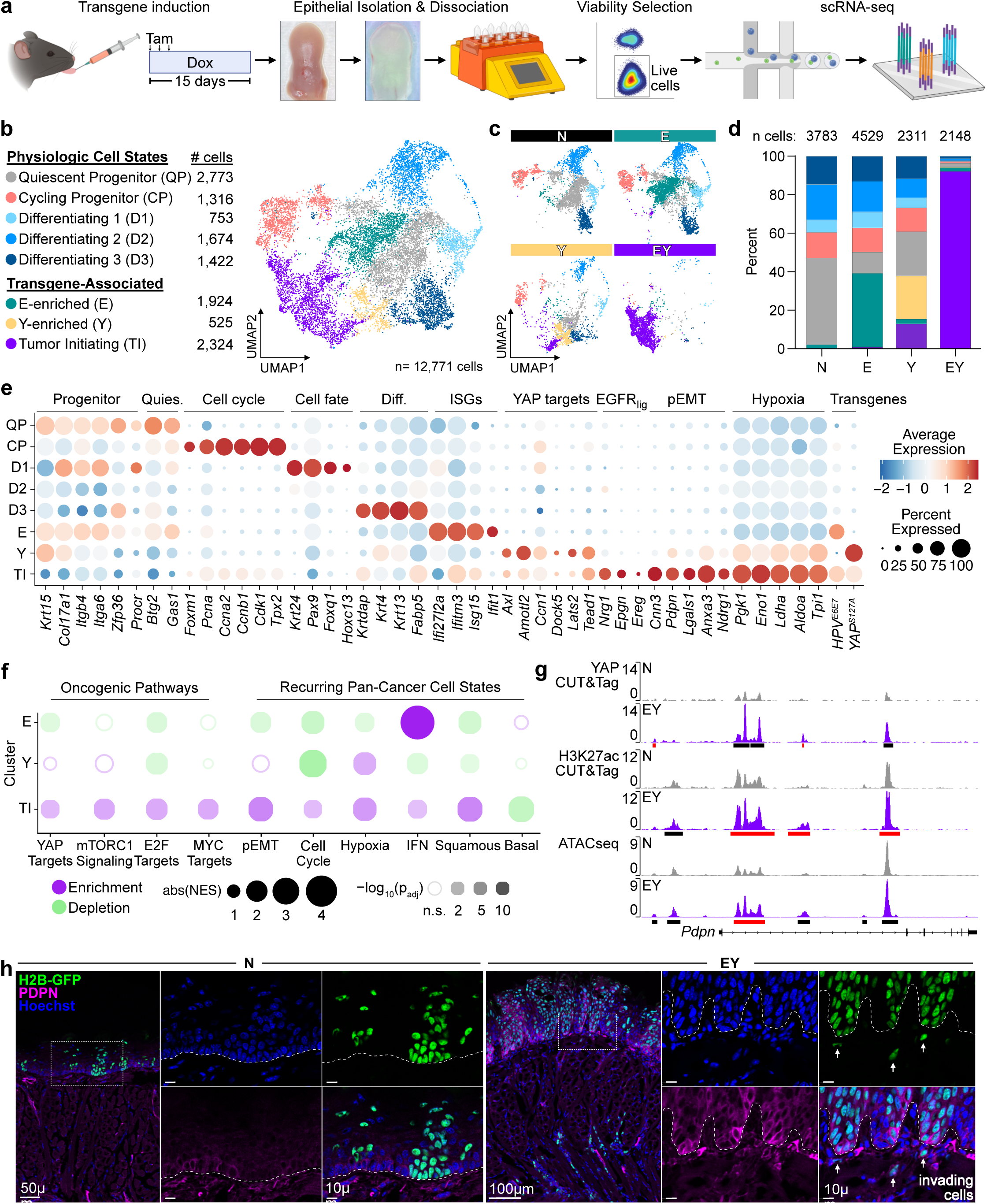
Transcriptional programs in nascent carcinoma at single cell resolution. (a) Experimental approach for scRNAseq of tongue epithelia. (b) UMAP of physiologic and transgene-associated epithelial cell states. (c) Individual UMAPs by transgenic condition demonstrating contributions to overall epithelial cell UMAP shown in b. n=2 mouse tongue epithelia per transgenic condition. (d) Distribution of cell states in tongue epithelial cells from each transgenic condition. n, total cells per transgenic condition. (e) Relative expression of representative cell state associated genes and the *HPV^E6E7^* and *YAP^S127A^* transgenes by cluster. (f) GSEA of molecular signatures and recurring cancer cell states across the transgene-enriched clusters. Dot color indicates enrichment (purple) or depletion (green). Dot size encodes the absolute value (abs) of the normalized enrichment score (NES). Circle opacity represents −log_10_ of the adjusted p-value (p_adj_); circle are hollow if p_adj_>0.05. For gene set details, please see Supplementary Table 3. (g) IGV tracks of YAP CUT&Tag, H3K27ac CUT&Tag, and ATACseq peaks at the *Pdpn* gene locus. Black bars indicate significant peaks. Red bars indicate EY-gained peaks. (h) Representative images of H2B-GFP and PDPN expression in N and EY epithelia 10 days after transgene induction. Dashed white line indicates the basement membrane.

We examined DEGs and performed GSEA comparing epithelial clusters to published transcriptional signatures for OEPC states^12,44–46^ to assign physiologic clusters with Quiescent Progenitor (QP), Cycling Progenitor (CP), and Differentiating (D1-D3) phenotypes. QP displayed enrichment of basal/progenitor markers, stem cell maintenance factors, and antiproliferative factors. CP cells showed enrichment for cell cycle transition factors, and depletion of differentiation markers. Differentiating cells exhibited a continuum of maturation states from D1 to D3. D1 cells were enriched for markers of lineage commitment,^47^ and factors required for transit amplifying cell proliferation^46,48^ and cell fate determination.^49,50^ D2 and D3 cells showed increasing expression of terminal differentiation markers (**Fig. 4e, Extended Data Fig. 6c-e, and Supplementary Table 5**).

Transgene expression resulted in the emergence of three clusters not observed in normal epithelia. E-enriched cells displayed high expression of *E6-E7* and interferon-stimulated genes known to be upregulated by *E6-E7*.^51,52^ Y-enriched cells showed high expression of *YAP1^S127A^* and YAP target genes (**Fig. 4e**). Interestingly, EY epithelial cells were found primarily within a unique cluster most consistent with tumor initiating cells (TIC). TIC expressed *E6-E7* and *YAP1^S127A^*, and displayed high expression of pEMT and hypoxia transcripts (**Fig. 4e**).^53,54^ The TIC cluster was markedly enriched for YAP, mTORC1, E2F, and MYC-driven programs, and modules defining pan-cancer cell states^53^ including pEMT, hypoxia, cell cycle, interferon response, and squamous differentiation (**Fig. 4f, Extended Data Fig. 6f**), suggesting key oncogenic pathways were exclusively activated in TIC.

TIC also highly expressed the pEMT transcript *Pdpn*, which we found to be regulated by YAP, demonstrating gained YAP and H3K27ac binding sites, and increased chromatin accessibility in EY cells (**Fig. 4g**). Importantly, PDPN was highly expressed in epithelial cells at the invasive front in EY epithelia (**Fig. 4h**). Other TIC cluster defining genes, including *Slpi*, *Anxa3*, *Klk10*, and *Eno1* also displayed direct YAP-mediated activation (**Extended Data Fig. 6g-j**). Multiple lines of evidence thus suggest that YAP drives OEPC to TIC reprogramming and tumor initiation.

### Tumor initiating cells co-opt collagenase-expressing G-MDSCs to facilitate tumor invasion

Our integrated multiomic analyses indicated that YAP promotes inflammatory responses, which may contribute to tumor initiation. Analysis of bulk transcriptomes showed granulocyte-specific markers and granulocyte-recruiting chemokines and cytokines ranked among the most highly upregulated genes in EY epithelia (**Extended Data Fig. 7a**). Integrative transcriptomic analysis of cultured cells and epithelial tissues identified tissue-specific transcripts, which included myeloid and lymphoid cell-specific transcripts, suggesting that EY activation induces immune cell infiltration (**Extended Data Fig. 7b,c**). Accordingly, YAP-activated epithelia showed a marked increase in infiltrating CD45^+^ immune cells by flow cytometry (**Fig. 5a**). Differences in immune cell types were further analysed by scRNAseq, identifying 11,286 immune cells distributed across 13 clusters (**Fig. 5b**, **Extended Data Fig. 7d,e; Supplementary Table 6**). Remarkably, myeloid derived suppressor cells (MDSCs) comprised 65% of immune cells in EY epithelia, with approximately equal proportions of monocytic (M-MDSC) and granulocytic (G-MDSC) MDSCs (**Fig. 5c,d**). Consistently, immunofluorescent microscopy demonstrated Ly6G^+^ G-MDSCs infiltration in close proximity to invading tumor cells (**Fig. 5e**).

**Figure 5.**
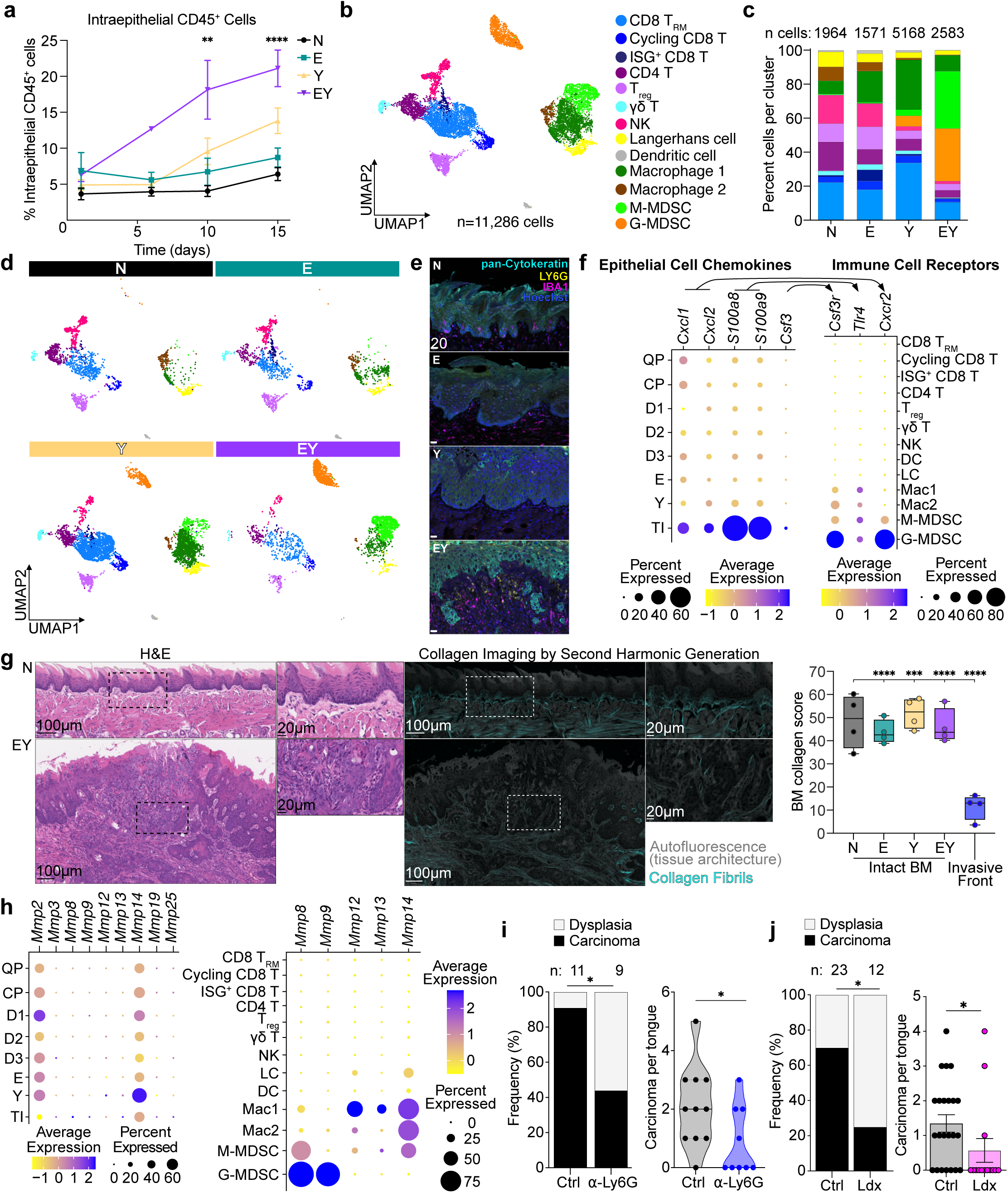
TICs co-opt collagenase-expressing G-MDSCs to facilitate tumor invasion. (a) Percent of CD45^+^ cells present in tongue epithelia at 1, 6, 10, and 15 days after transgene induction by flow cytometry. Each time point was analyzed individually by ANOVA with Tukey correction for multiple comparisons; means with SEM are shown: **p<0.01, ****p<0.0001. (b) UMAP of immune cell clusters identified by scRNAseq. (c) Immune cell types stratified by transgenic condition. (d) Distribution of immune cell types by transgenic condition. (e) Representative immunofluorescence image of pan-Cytokeratin, LY6G, and IBA1 expression in N, E, Y, and EY tongue epithelia 20 days after transgene induction. (f) Expression of chemokine genes among epithelial cell clusters and corresponding receptor genes among immune cell clusters. (g) Representative H&E-stained sections (left) and collagen imaging by second harmonic generation (middle) in control epithelia (N) and an infiltrative carcinoma lesion (EY) 20 days after transgene induction. Right: fluorescent intensity of second harmonic generation signal at intact basement membrane of transgenic tongue epithelia and at the invasive front of EY carcinoma 20 days after transgene induction. ANOVA with Tukey correction for multiple comparisons. Boxplots show median, interquartile range (IQR), and range. ***p<0.001, ****p<0.0001. (h) Collagenase gene expression across epithelial and immune cell states. (i) Frequency of carcinoma (left) and carcinoma burden per tongue (right) after treatment of transgened induced EY mice with vehicle or anti-LY6G depleting antibody. (j) Frequency of carcinoma (left) and carcinoma burden per tongue (right) after treatment of transgened induced EY mice with vehicle or the CXCR1/2 dual inhibitor ladarixin (Ldx).

As noted above, TNF signaling and inflammatory response represented highly enriched YAP-regulated pathways. Further investigation of these pathways revealed *Tnf*, *Csf3*, *Cxcl1*, and *Cxcl2* represented YAP-regulated genes (**Extended Data Fig. 8a**). In oral epithelia, TNF, G-CSF, IL-23, and IL-17 initiate cytokine-chemokine cross-talk which induces sustained granulocyte recruitment.^55^ In line with this model, EY epithelia showed increased abundance of transcripts for *Tnf*, *Csf3*, and the granulocyte-specific chemokines *Cxcl1* and *Cxcl2* (**Extended Data Fig. 8b**). Accordingly, TNF, IL-17, G-CSF, CXCL1, and CXCL2 proteins were also elevated in EY epithelia (**Extended Data Fig. 8c**). Ligand-receptor analysis of scRNAseq data revealed that TIC express chemokine ligands whose cognate receptors are expressed by G-MDSCs (**Fig. 5f**, **Extended Data Fig. 8d**). The above findings suggest that TIC may promote G-MDSC recruitment.

The basement membrane represents an anatomic barrier against invasive carcinoma. Evaluation of transgene-induced EY epithelia by second harmonic generation microscopy^56^ demonstrated a dramatic reduction in fibrillar collagen at the invasive front of nascent invasive carcinoma (**Fig. 5g**). Given that G-MDSCs were present at the EY invasive front, we investigated expression of basement membrane extracellular matrix (ECM) remodeling enzymes by TIC and immune cells. While TIC expressed genes associated with cell motility and invasion, they did not express collagenases. Conversely, G-MDSCs highly expressed collagenases *Mmp8* and *Mmp9,* and MMP-8 and proMMP-9 proteins were detected in EY epithelia (**Fig. 5h, Extended Data Fig. 8e**). Treatment of transgene-induced EY mice with anti-LY6G depleting antibody significantly reduced the number of G-MDSCs at the invasive front and diminished carcinoma formation (**Fig. 5i** and **Extended Data Fig. 8f**). Similarly, CXCR1/CXCR2 inhibition with the small molecule inhibitor ladarixin diminished the presence G-MDSCs near EY epithelia basement membranes and decreased carcinoma incidence and burden in EY mice (**Fig. 5j** and **Extended Data Fig. 8g**). Overall, these findings support that YAP-activated TIC produce cytokines and chemokines that recruit G-MDSCs, which in turn may help facilitate tumor invasion.

### YAP promotes mTOR signaling

We next explored the role of mTOR and EGFR pathway activation in EY tumor initiation. Enrichment of YAP and mTOR transcriptional signatures by GSEA^57^ was observed in YAP-expressing epithelia (**Fig. 6a, Extended Data Fig. 9a, Supplementary Table 3**). Consistently, EY epithelia showed a pronounced increase in phospho-S6 (pS6), a downstream marker of mTOR activity^58^ (**Fig. 6b,c**). Consistent with the mutually compensatory functions of YAP and TAZ,^59^ combined knockdown of *YAP* and *TAZ* was required to diminish YAP target (CYR61) and pS6 abundance in Cal27 and Cal33 cells (**Fig. 6d**, **Extended Data Fig. 9b**). Intriguingly, *YAP/TAZ* knockdown also resulted in diminished pEGFR. Together, these findings, and the enrichment of YAP and mTOR transcriptional signatures in EY tumors, suggested a potential mechanistic link between YAP activation and mTOR signaling.

**Figure 6.**
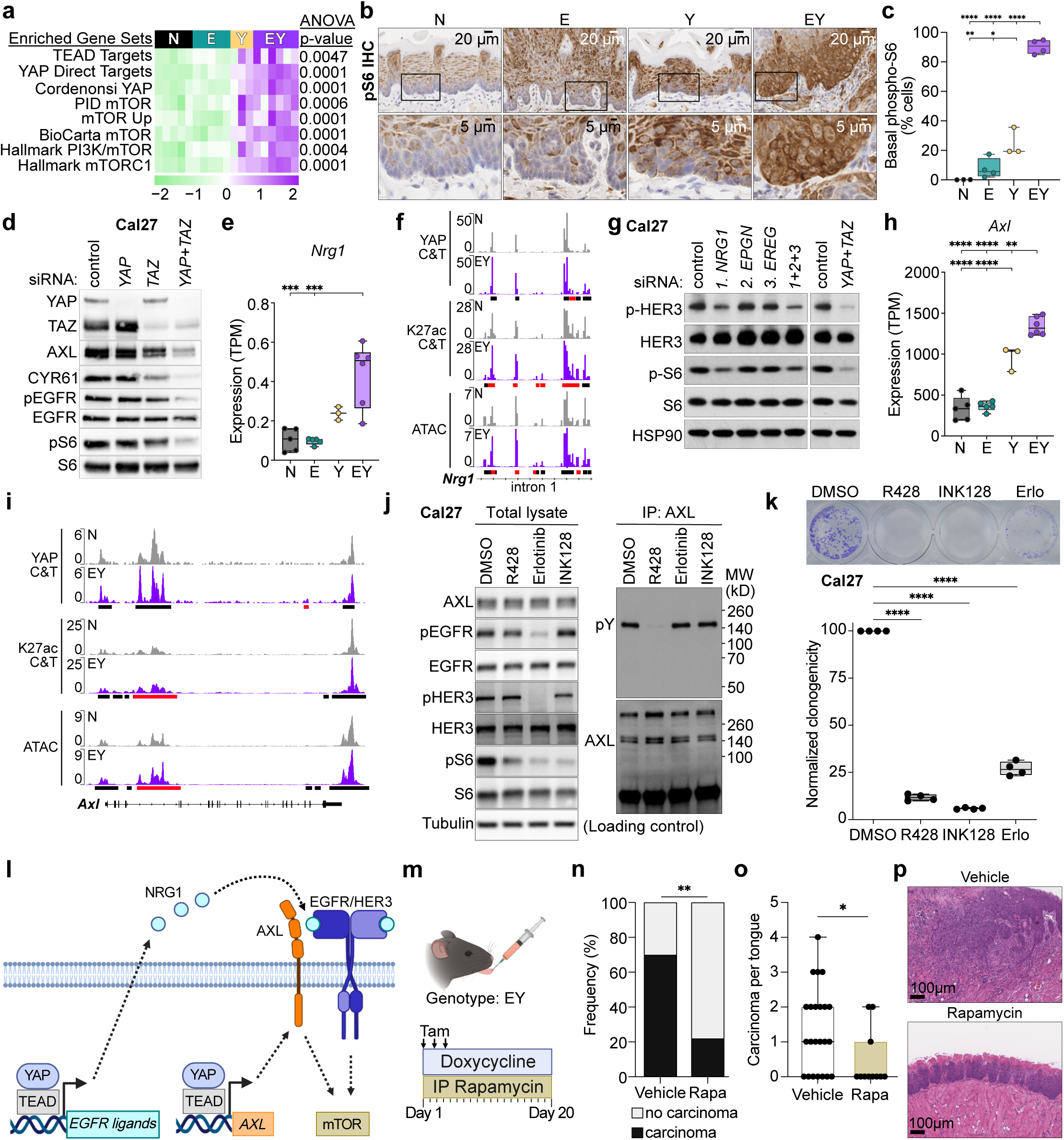
YAP promotes mTOR signaling. (a) GSEA for gene sets associated with TEAD- and YAP-target gene expression, and mTOR activation. For gene set details, please see Supplementary Table 3. (b) Representative IHC and (c) quantification of phospho-S6-positive cells in basal epithelial cells. (d) The effects of siRNA-mediated knockdown of *YAP*, *TAZ*, and *YAP*+*TAZ* on phospho- and total EGFR and S6 expression in Cal27 whole cell lysate. (e) *Nrg1* expression in transgenic tongue epithelia by RNAseq. (f) IGV tracks of YAP CUT&Tag, H3K27ac CUT&Tag, and ATACseq peaks at a segment of the *Nrg1* gene locus. IGV tracks of full *Nrg1* locus shown in Extended Data Fig. 9. Black bars indicate significant peaks. Red bars indicate EY-gained peaks. (g) The effects of siRNA-mediated knockdown of *NRG1*, *EREG*, *EPGN* on phospho- and total HER3, and S6 expression in Cal27 whole cell lysate. (h) *Axl* expression in transgenic tongue epithelia by RNAseq. (i) IGV tracks of YAP CUT&Tag, H3K27ac CUT&Tag, and ATACseq peaks at the *Axl* gene locus. Black bars indicate significant peaks. Red bars indicate EY-gained peaks. (j) Left: Immunoblots of Cal27 whole cell lysate showing effects of AXL (R428), EGFR (Erlotinib), and mTOR (INK128) inhibitors on phospho- and total EGFR, HER3, and S6. Right: immunoprecipitation of AXL from Cal27 whole lysates followed by immunoblot for phospho-tyrosine (pY) showing effects of R428, Erlotinib, or INK128 on phospho- and total AXL. (k) Representative wells (top) and quantification (bottom) of clonogenic assays in Cal27 cells treated with DMSO, R428, INK128, or Erlotinib (Erlo). (l) Proposed model for YAP-mediated transcriptional activation of mTOR via *AXL*, *NRG1*, and the EGFR/HER3 axis. (m) Schematic of *in vivo* mTOR inhibition with rapamycin. (n) Carcinoma burden after treatment of transgene-induced EY mice with vehicle or rapamycin. Fisher’s exact test: **p<0.01 (o) Frequency of carcinoma after treatment of transgene-induced EY mice with vehicle or rapamycin. Two-tailed Mann-Whitney test: *p<0.05 (p) Representative H&E staining of vehicle- and rapamycin-treated EY mouse tongues. Panels a, c, e, h, and k were analyzed by ANOVA with Tukey correction for multiple comparisons. For all panels with asterisks denoting significance: *p<0.05, **p<0.01, ***p<0.001, ****p<0.0001.

In search of underlying mechanisms, we interrogated transcriptomic, CUT&Tag, and ATAC data for YAP-regulated genes that may induce EGFR signaling. We identified multiple effectors and ligands of EGFR signaling to be directly regulated by YAP. The HER3 ligand *Nrg1* and EGFR ligands *Ereg* and *Epgn* were upregulated and showed gains in YAP binding, H3K27ac, and chromatin accessibility in EY cells (**Fig. 6e-f, Extended Data Fig. 9c-f**). Knockdown of *NRG1* but not *EREG* or *EPGN* resulted in decreased pS6 in representative HNSC cell lines, Cal27 and HN12 cells, which we have previously shown to be dependent on YAP for survival and proliferation^16^ (**Fig. 6g, Extended Data Fig. 9g-h**). Furthermore, the EGFR activator^60^ *AXL* demonstrated YAP-mediated transcriptional regulation (**Fig. 6h-i**). Small molecule inhibition of the EGFR/HER3 axis with erlotinib and AXL inhibition with R428, which resulted in reduced tyrosine phosphorylation of the corresponding receptors, as well as direct mTOR inhibition with INK128 reduced pS6 in Cal27 cells (**Fig. 6j**). Finally, treatment with erlotonib or R428 diminished tumor cell clonogenicity *in vitro* in Cal27 cells (**Fig. 6k, Extended Data Fig. 9i**). Taken together, these data provide a mechanistic framework by which YAP may induce mTOR activation via direct transcriptional activation of *NRG1* and *AXL* (**Fig. 6l**).

To investigate whether YAP-mediated mTOR activation contributed to YAP-mediated tumor initiation, we treated KEY primary cells with siRNAs targeting *Yap/Taz* and *Tead1/Tead4* as well as small molecule mTOR (INK128) and YAP-TEAD (VT104) inhibitors, and observed significant reductions in clonogenic growth (**Extended Data Fig. 9j-k**). In light of these findings, we treated transgene-induced EY mice with the mTOR inhibitor rapamycin. mTOR inhibition resulted in a remarkable decrease in carcinoma formation (**Fig. 6m-p**), thus supporting that YAP-mediated mTOR activation contributes to tumor initiation.

### TIC transcriptional programs are enriched in HNSC and associated with poor prognosis

We next tested the significance of YAP and mTOR signaling and TIC programs in human HNSC tumors. We examined human subjects with HNSC from a molecularly and clinically well-defined cohort, The Cancer Genome Atlas HNSC cohort (TCGA-HNSC). Among 43 TCGA-HNSC subjects with RNAseq data for paired tumor and normal tissue, we found we observed that both YAP and mTOR signaling were enriched in tumor compared to normal tissue (**Fig. 7a**). Intriguingly, we found a strong positive relationship between YAP and mTOR pathway activation across TCGA-HNSC (**Fig. 7b**). Subjects with above-median YAP or mTOR pathway activation displayed worse overall and disease-free survival (**Fig. 7c-f**), suggesting that beyond tumor initiation, YAP and mTOR pathway activation may contribute to tumor progression and influence prognostic outcomes.

**Figure 7.**
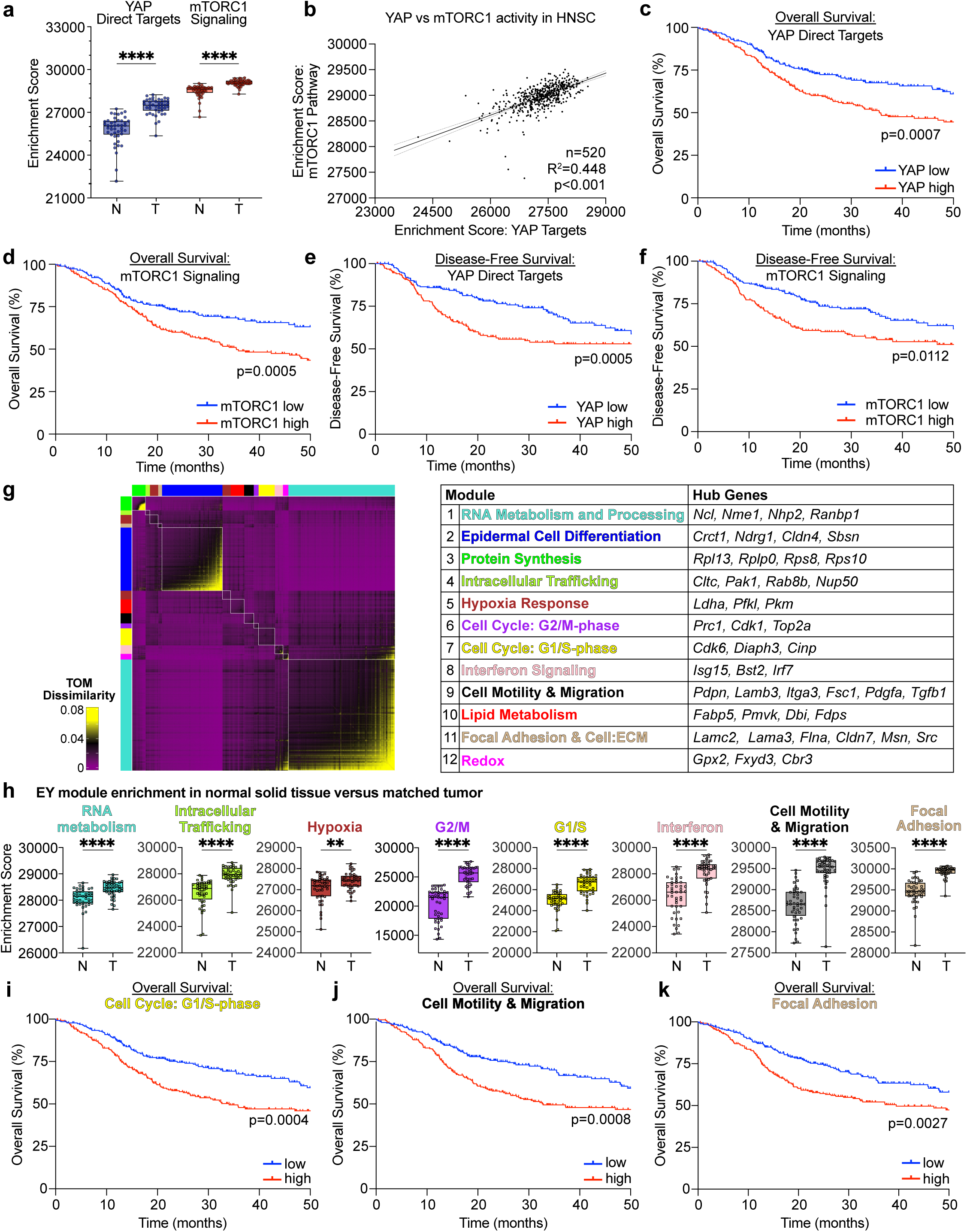
TIC programs are enriched in HNSC and associated with poor prognosis. (a) mTORC1 and YAP pathway enrichment in malignant tumors (T) compared to matched normal solid tissues (N) by single sample GSEA among subjects in the TCGA-HNSC cohort (n=43 subjects). ****p<0.0001. Two-tailed Mann-Whitney test: ****p<0.0001. (b) Correlation of YAP and mTORC1 pathway activity by ssGSEA among TCGA-HNSC samples (n=520). (c-d) Kaplan-Meier plots demonstrating worse overall survival (n=520) among TCGA-HNSC subjects with greater than median (c) mTORC1 and (d) YAP pathway activity. (e-f) Kaplan-Meier plots demonstrating worse disease-free survival (n=393) among TCGA-HNSC subjects with greater than median (e) mTORC1 and (f) YAP pathway activity. For gene set details, please see Table S1. (g) Left: Weighted gene co-expression network analysis (WGCNA) heatmap displaying topological overlap matrix (TOM) dissimilarity indices among genes in TP cluster cells. Right: Table of WGCNA modules and selected genes identified by Metascape and Enrichr. See also Supplementary Table 6. (h) Module enrichment in malignant tumors (T) compared to matched adjacent normal solid tissues (N) by single sample GSEA among TCGA-HNSC subjects (n=43 subjects with matched T and N samples). Boxplots show median, interquartile range (IQR), and range. Two-tailed paired T-test: **p<0.01, ****p<0.0001. (i–k) Kaplan-Meier plots demonstrating worse overall survival (n=393) among TCGA-HNSC subjects with greater median enrichment for the (c) G1/S, (d) Cell Motility & Migration, and (e) Focal Adhesion EY-modules.

We next decomposed the TIC cluster into co-expressed transcriptional modules using high dimensional weighted gene co-expression network analysis (hdWGCNA)^61,62^ (**Fig. 7g, Supplementary Table 7**). We noted enrichment of 8 of 12 modules in tumor compared to normal tissue, unveiling RNA metabolism and processing, intracellular trafficking, hypoxia response, G1/S and G2/M cell cycle progression, interferon response, motility and migration, and cytoskeleton and cell polarity as distinguishing features of the HNSC transcriptome (**Fig. 7h, Extended Data Fig. 10a**). Among all modules, enrichment for the G1/S cell cycle, motility and migration, and focal adhesion modules were associated with worse disease-free and overall survival (**Fig. 7i-k, Extended Data Fig. 10b-d**). These findings suggest that tumor initiating cells display coherent transcriptional programs, which are enriched in aggressive HNSC and associated with worse HNSC outcomes.

## DISCUSSION

The overwhelming majority of work investigating cancer-driving mechanisms has relied on established tumors, which limits distinction between processes governing tumor initiation and progression. Using a spatiotemporally controlled *in vivo* system targeting genomic alterations to a single pool of epithelial progenitor cells, we show that unrestrained YAP activation in the context of HPV oncoprotein-mediated *TP53* and *CDKN2A* inhibition induces carcinoma with rapid kinetics and high penetrance. This system enabled multi-modal genome-wide exploration of YAP-mediated processes driving OEPC reprogramming into tumor initiating cells.

Tumor initiating cells were endowed with hallmarks driving invasion, including the activation of invasive (pEMT) and inflammatory (G-MDSC recruitment) programs, early in carcinogenesis. Our findings suggest pEMT is not merely an element driving cancer progression, but a defining feature of tumor initiating cells. Furthermore, EY carcinoma displayed tumor invasive fronts with extensive basement membrane collagen remodeling, concordant with the perspective that ECM modification and invasion are defining features of premalignancy-to-cancer transition. Remarkably, single cell analysis revealed that TIC do not express collagenases, which are important drivers of ECM remodeling. This suggests that TIC are not endowed with an intrinsic ability to intitate invasion of surrounding tissues. Mechanistically, TIC may instead express multiple cytokines and chemokines, which in turn promote the recruitment of collagenase-expressing G-MDSCs to the invasive front, thus facilitating tumor infiltration. This paracrine mechanism supports that cell-cell communication networks between precancer and myeloid cells may ultimately enable cancer initiation, thereby providing an opportunity to halt the progression of premalignant disease by pharmacologically interfering with TIC-myeloid cell crosstalk.

Another unexpected finding of our *in vivo* system was that carcinogenesis did not appear to require genomic alterations in the PI3K/AKT/mTOR signaling axis.^42,63–65^ However, >70% of human HNSCs exhibit widespread activation of YAP^10,16^ and mTOR^65^ in the absence of genomic alterations in components of the PI3K-mTOR pathway. Mechanistically, our findings support a novel model in which YAP mediates transcription of *NRG1* and *AXL*, which in turn converge to activate the mTOR signaling network via HER3 and EGFR signaling. HER3/EGFR involvement in mTOR pathway activation is in line with our prior findings that persistent tyrosine phosphorylation of HER3 underlies aberrant PI3K/AKT/mTOR signaling in *PIK3CA* wildtype HNSC^66,67^, albeit what leads to HER3/EGFR activation was not known. We now provide evidence that dysregulated YAP-driven cis-regulatory activation of transcription and chromatin accessibility may underlie mTOR activation in HNSCs lacking genomic alterations in the PI3K/AKT/mTOR axis. Furthermore, mTOR inhibition with well-known pharmacological agents revealed that mTOR activation is required for YAP-mediated tumor progression. Indeed, this YAP-mediated autocrine loop initiating NRG1/HER3/EGFR-mTOR signaling, concomitant with AXL expression, may provide actionable targets for future clinical investigation.

In summary, we demonstrate that a genetically-defined, traceable system simultaneously activating oncogenic pathways and disabling tumor suppressive mechanisms in normal oral epithelial progenitor cells induces the emergence of a distinct cancer initiating stem-like cell state. Through multimodal analysis of nascent TIC at the single cell level *in vivo*, we define tumor-autonomous transcriptional programs and TIC-TME cross-talk as tumor initiating events during invasive carcinoma formation (**Fig. 8**). This conceptual framework of cancer initiation has the potential to open multiple novel avenues for early intervention, including precision targeting of tumor cell-autonomous cancer initiating signaling pathways, and disrupting TIC-TME networks mediating the development of invasive carcinoma.

**Figure 8.**
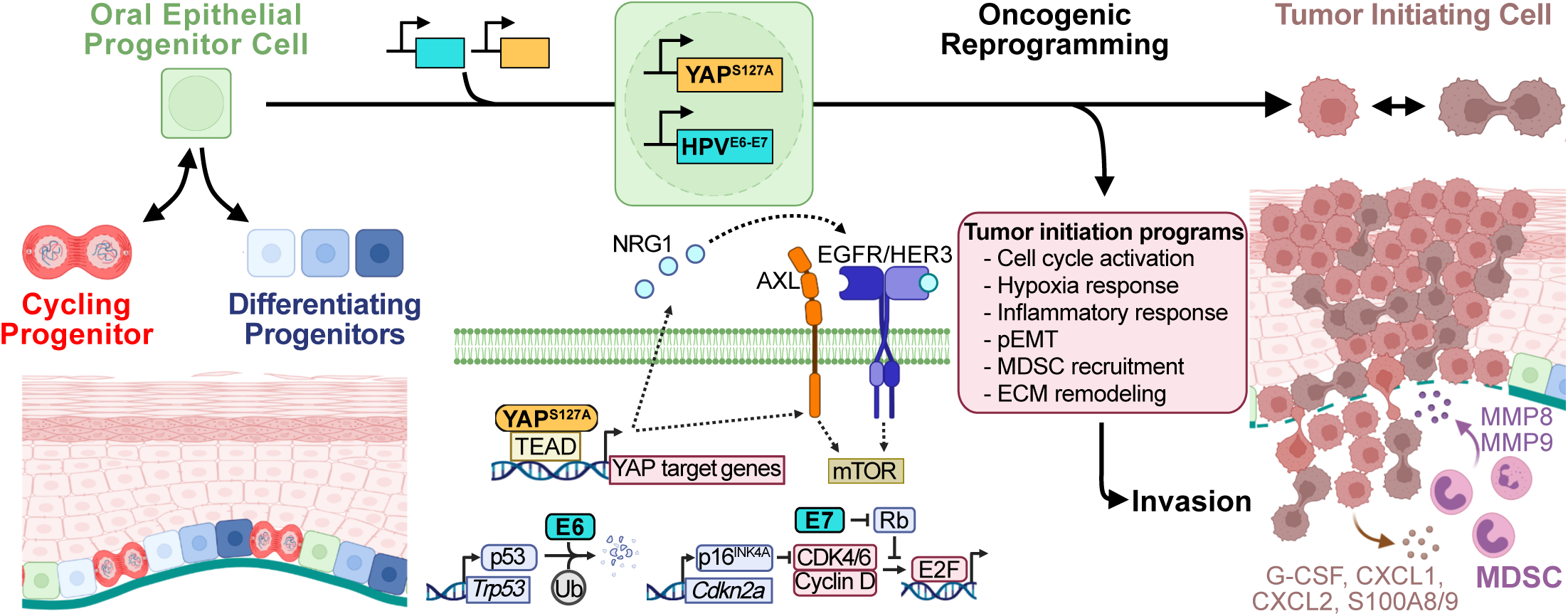
Genetically-defined oncogenic and tumor suppressive pathway alteration in normal oral epithelial progenitor cells defines tumor initiating events.

## ACKNOWLEDGEMENTS

This work was supported by Stand Up To Cancer No. 308268 (FF & JSG); NIDCD T32DC000028 (FF); American Head and Neck Society Pilot Grant No. 1061310 (FF); NIAID T32AI007036 & AP Giannini Postdoctoral Research Fellowship and Leadership Award (SIR); Burroughs Welcome Fun Career Award for Medical Scientists (SIR); National Defense Sciences and Engineering Graduate (NDSEG) Fellowship Program (LMC); NCI R25CA221779 (PYAQ); Tobacco-Related Disease Research Program Pre-Doctoral Award T32DT4965 (TSH); Howard Hughes Medical Institute Hanna H. Gray Fellowship (QS); State of California Initiative to Advance Precision Medicine Award OPR18112 & GCBSR shared resources at the UCSD Moores Cancer center NCI P30CA023100 (JAC, KME, & PT); NIDCR R01DE030497 & NIDCR R01DE026644 (JSG); NCI R01CA257505, NIDCR U01DE028227, NIDCR R01DE026870 (PT & JSG).

We are thankful for Katarzyna Dobaczewska, Sarah McArdle, and Brett Laffey of the LJI Microscopy & Histology Core and Kersi Pestonjamasp of the Moores Cancer Center Microscopy Core for helpful insights and guidance on experimental design; Cheryl Kim, Emily Von Gerichten, and Semra Sehic of the LJI Flow Cytometry Core for assistance with FACS experiments; Suzie Alarcon and Hannah Dose of the LJI NGS Core for assistance with scRNAseq experiments. The BD FACSAria II at LJI was funded by NIH equipment grant S10RR027366. The NovaSeq6000 at LJI was acquired through the Shared Instrumentation Grant Program S10OD025052. The results shown here are in part based upon data generated by the TCGA Research Network: https://www.cancer.gov/tcga. We appreciate the generous contribution of the TCGA-HNSC donors and participating research groups who made this resource available.

## AUTHOR CONTRIBUTIONS

Conceptualization: FF, JSG; Methodology: FF; Validation: FF, SIR, LMC; Formal Analysis: FF, SIR, LMC, PM, K.Sato; Investigation: VB, FF, SIR, LMC, K.Sato, PAQ, WMGG, TSH, KME, AO, AAM, KK, K.Sakaguchi, ZM; Resources: JSG, PT, PM, ODK, QS, JAC; Data Curation: FF, SIR, LMC, K.Sato; Writing, Original Draft: FF, SIR, JSG; Writing, Review & Editing: FF, SIR, JSG, ODK, QS, KSato, PT, ZM; Visualization: FF; Supervision: JSG, PT, QS, ODK, JAC; Project Administration: FF, JSG; Funding Acquisition: FF, JSG.

## DECLARATION OF INTERESTS

J.S.G. has received other commercial research support from Kura Oncology, Mavupharma, Dracen, Verastem, and SpringWorks Therapeutics, and is a consultant/advisory board member for Domain Therapeutics, Pangea Therapeutics, and io9, and founder of Kadima Pharmaceuticals. The remaining authors declare no competing interests.

## EXTENDED DATA FIGURE LEGENDS

**Extended Data Figure 1. Spatiotemporally controlled YAP and E6-E7 activation in OEPCs**

(a) Representative images of nuclear YAP protein in tissues by IHC in HPV negative (left) and HPV positive (right) malignant human HNSC tissue.

(b) Percent of cells with nuclear YAP in normal human tissue (n=12) and malignant HNSC (n=44) from tissue microarrays stratified by HPV status. Two-tailed Mann-Whitney test: **p<0.01, ****p<0.0001.

(c) Percent of samples with positive nuclear YAP in normal human tissues (n=12) and malignant HNSC (n=44) stratified by HPV status.

(d) Overview of local transgene induction in lingual epithelia and strategies for validation of transgene activation at genomic and transcript levels, and *in situ*.

(e) Left: Genomic DNA extracted from tongue epithelia 10 days after transgene induction with tamoxifen and doxycycline treatment. Top three gel images show presence of transgene and internal control (ctrl) PCR products. Bottom gel image confirms Lox_p_-Stop-Lox_p_ (LSL) recombination for *LSL-rtTA*. Lane 1 mouse tongue epithelial gDNA from a tamoxifen and doxycycline untreated mouse bearing all transgenes. NTC, no template control. Right: Schematic of oligonucleotide primer design to assay LSL excision.

(f) qRT-PCR quantification of *HPV^E6^*, *HPV^E7^*, and *YAP^S127A^* expression in tongue epithelia 10 days after transgene induction. Boxplots show median, interquartile range (IQR), and range.

(g) Immunofluorescence image of tongue epithelia from a Krt14-CreER^+^LSL-rtTA^+^tetON_H2B-GFP^+^ mouse 6 days after transgene induction showing basal localization of KRT14 and ITGA6, and nuclear expression of H2B-GFP. Nuclei are counterstained with Hoescht.

(h) Total number of lesions per tongue 20 days after transgene induction. Means with SEM are shown.

(i) Hematoxylin and eosin (top) and pan-cytokeratin (pan-CK) IHC (bottom) stained tongue epithelia demonstrating infiltrative carcinoma in Y (left panels) and EY (right panels) mice. **Related to Fig. 1**.

**Extended Data Figure 2. YAP and E6-E7 activation in OEPCs induces tumor initiating cells**

(a) Experimental approach for the generation and orthotopic implantation of transgene-induced epithelial cell suspensions.

(b) Representative photographs of NSG mouse tongues after orthotopic implantation transgene-induced epithelial cell suspensions.

(c) Representative H&E stained section of an NSG mouse tongue after orthotopic implantation with transgene-induced EY epithelial cell suspension.

(d) Implanted tumor outgrowth frequency; n, number of orthotopically implanted mouse tongues. Between group comparisons were made using Fisher’s exact test with Bonferroni correction.

(e) Tumor volumes 10 days after implantation. Mean with SEM is shown. ANOVA with Tukey correction for multiple comparisons used for between group comparisons.

(f) Left: Gating strategy for FACS enrichment of H2B-GFP transgene positive cells derived from EY tongue epithelium based on GFP expression. Right: Representative fluorescence microscopy of primary EY cell cultures pre- and post-FACS enrichment.

(g) qRT-PCR quantification of *HPV^E6^*, *HPV^E7^*, and *YAP^S127A^* expression in FACS-purified N and EY cell cultures. Barplots show mean and SEM.

(h) Representative gross (left) and H&E stained sections (right) after implantation of 5,000 FACS-enriched EY cells in NSG mouse tongues.

For all panels with asterisks denoting significance: *p<0.05, **p<0.01, ***p<0.001, ****p<0.0001.

**Related to Fig. 1**.

**Extended Data Figure 3. Oncogenic transcriptional reprogramming defines YAP and E6-E7 activated epithelia**

(a) Schematic demonstrating strategy for bulk RNAseq of microdissected tongue epithelia at 15 days post-transgene induction.

(b) *HPV^E6E7^*, *YAP^S127A^*, and YAP target gene expression in transgenic epithelia transcriptomes.

(c) Two-dimensional principal component analysis of individual N, E, Y, and EY transcriptome profiles.

(d) Enriched cellular processes among EY unique DEGs by Gene Ontology.

(e) Single sample GSEA showing enrichment or depletion of oral epithelial stem cell states including Cell Cycle Progression (G1/S, G2/M); Stem Cell; and Differentiation (Diff 1 and Diff 2). These gene signatures were derived from cluster defining genes from Jones et al. For gene set details, please see Supplementary Table 3.

(f) Relative expression of EY-unique DEGs regulating oral epithelial cell identity.

(g) Single sample GSEA for all transgenic conditions for gene signatures related to cell cycle, differentiation, stemness, and EMT. For gene set details, please see Supplementary Table 3. Each row in panels b, e, f, and g was analyzed by ANOVA with Tukey correction for multiple comparisons: *p<0.05, **p<0.01, ***p<0.001, ****p<0.0001.

**Related to Fig. 2**.

**Extended Data Fig. 4. Invasive carcinoma is preceded by the expansion of a stem-like cell population**

(a-b) Lineage tracing by fluorescent microscopy using the *H2B-GFP* reporter transgene to track and quantify GFP^+^ nuclei. (a) Representative fluorescence images of and (b) percent basal H2B-GFP^+^ nuclei in tongue epithelial basal layer whole mounts 36 hours after transgene induction.

(c) Representative immunofluorescence images of H2B-GFP, KI67, and KRT15 expression in tongue epithelia 15 days after transgene induction. White dashed line represents the basement membrane.

(d) Related to c. Percent KI67^+^GFP^+^ nuclei in tongue epithelia.

(e) Representative immunofluorescence images of H2B-GFP, P63, and ITGA6 expression in tongue epithelia 15 days after transgene induction. White dashed line represents the basement membrane.

(f) Related to e. Percent P63^+^GFP^+^ nuclei in tongue epithelia.

(g) Related to e. Quantitative spatial analysis showing normalized fluorescent intensity of ITGA6 signal from basement membrane to epidermal surface. Probability cloud shows 95% confidence interval.

(h-j) Magnified images of cells at the invasive front from Fig. 2m demonstrating (h) KI67^+^ nuclei in Fig. 2i, (i) P63^+^ nuclei in Fig. 2k, (j) SOX2^+^ nuclei in Fig. 2m.

Panels b, d, and f were analyzed by ANOVA with Tukey correction for multiple comparisons: *p<0.05, **p<0.01, ***p<0.001, ****p<0.0001. Boxplots show median, interquartile range (IQR), and range.

**Related to Fig. 2**.

**Extended Data Figure 5. Epigenetic reprogramming in YAP and E6-E7 activated epithelia and EY cell lines**

(a) Left: DESeq interaction model comparing genes enriched in EY compared to N tissue versus in EY compared to N primary culture cells. Discordant genes between epithelial tissue and cell lines are shown in red, and concordant genes (88%) in blue. Concordance threshold based on log fold difference, p > 0.05. Right: Frequency of genes with an observed difference (%) in EY versus N across EY and N tissues and cell lines.

(b) Left two Venn diagrams: overlap of transcripts detected in N (left) or EY (middle) epithelial tissue (top circles) versus primary cell culture (bottom circles). Right Venn diagram: overlap of tissue-specific N or EY transcripts identify 741 transcripts identified only in EY epithelial tissue.

(c) Enrichr analysis of the 2366 tissue-specific transcripts detected in both N and EY tissue found in the Cell Marker 2024 library.

(d) Fraction of YAP CUT&Tag peaks that localize to genomic region annotations defined using HOMER.

(e) Absolute distance of YAP CUT&Tag peaks to the nearest transcription start site (TSS).

(f) Top transcription factor binding motifs enriched in the EYgaine YAP CUT&Tag binding sites. P values were calculated using HOMER.

**Related to Fig. 3**.

**Extended Data Figure 6. Single cell analysis of E6-E7 and YAP activated epithelial cells**

(a) *E6-E7* transgene expression stratified by genotype.

(b) *YAP1^S127A^* transgene expression stratified by genotype.

(c) Expression of the top 6 cluster defining genes among epithelial cell clusters. Dot size denotes the percentage of cells within a cluster expressing each transcript and the color indicates average gene expression across all cells in each cluster.

(d) GSEA of physiologic cell states across epithelial cell clusters. Circle color indicates enrichment (red) or depletion (blue). Circle size encodes the absolute value (abs) of the normalized enrichment score (NES). Circle opacity represents −log_10_ of the adjusted p-value (p_adj_); circles are hollow if p_adj_>0.05. For gene set details, please see Supplementary Table 3.

(e) GSEA enrichment plots for the Jones Stem Cell, Jones Differentiation, and Jones G1/S G2/M physiologic OEPC gene sets across epithelial cell clusters.

(f) Feature plots showing expression of the YAP Direct Target, mTORC1 Signaling, E2F Targets, MYC Targets, and the Barkley *et al.* recurring cancer cell state gene sets: partial epithelial to mesenchymal transition (pEMT), hypoxia, and squamous differentiation in single epithelial cells.

(g) IGV tracks of YAP CUT&Tag, H3K27ac CUT&Tag, and ATACseq peaks at the *Slpi* gene locus.

(h) IGV tracks of YAP CUT&Tag, H3K27ac CUT&Tag, and ATACseq peaks at the *Anxa3* gene locus.

(i) IGV tracks of YAP CUT&Tag, H3K27ac CUT&Tag, and ATACseq peaks at the *Klk10* gene locus.

(j) IGV tracks of YAP CUT&Tag, H3K27ac CUT&Tag, and ATACseq peaks at the *Eno1* gene locus.

For panels g-j: black bars indicate significant peaks. Red bars indicate EY-gained peaks.

**Related to Fig. 4**.

**Extended Data Figure 7. Single cell analysis of E6-E7 and YAP polarized epithelial immune infiltrate**

(a) RNAseq data from bulk epithelia for granulocyte markers and cytokines among EY DEGs. Means with SEM are shown.

(b) Top: Venn diagram showing overlap of EY tissue-specific and EY upregulated genes. Bottom: Volcano plot of epithelial bulk RNAseq data comparing EY vs N highlighting EY upregulated tissue-specific genes. Myeloid cell associated genes enriched in EY epithelial tissue are shown in red and lymphoid cell associated genes enriched in EY epithelial tissue are shown in blue.

(c) Bulk RNAseq expression in TPM in transcripts per million (TPM) of the EY upregulated tissue-specific myeloid cell markers, *Retnlg*, *Cd177*, and *Mmp8*, across N and EY samples, and epithelial tissue and primary culture conditions.

(d) Expression of select cell type defining genes among immune cells by scRNAseq.

(e) Dot Plot demonstrating GSEA of published cell-type defining gene lists across immune cell types. Dot color indicates enrichment (Orange) or depletion (Blue). Dot size represents the absolute value of the normalized enrichment score (NES). Dot opacity represents −log_10_ of the adjusted p-value. For gene set details, please see Supplementary Table 3.

**Related to Fig. 5**.

**Extended Data Figure 8.**

(a) IGV tracks of YAP CUT&Tag, H3K27ac CUT&Tag, and ATACseq peaks at the *Tnf, Csf3, Cxcl1, Cxcl2* gene loci. Black bars indicate significant peaks. Red bars indicate EY-gained peaks.

(b) Bulk RNAseq expression in TPM of *Tnf, Csf3, Cxcl1, Cxcl2* across N and EY samples, and epithelial tissue and primary culture conditions.

(c) G-CSF, IL-17, TNFα, CXCL1, and CXCL2 protein abundance in epithelial lysates from N, E, Y, and EY mice 15 days after transgene induction.

(d) Bulk RNAseq expression in TPM of *Cxcr2, Csf3r* across N and EY samples, and epithelial tissue and primary culture conditions.

(e) MMP8 and pro-MMP9 protein abundance in epithelial lysates from N, E, Y, and EY mice 15 days after transgene induction.

(f) Left: Experimental approach for depletion of LY6G^+^ G-MDSCs in transgene induced EY mice. Right: Representative images of CD45^+^ and Ly6G^+^ immune infiltrates in EY mouse tongue epithelia 20 days after transgene induction after treatment with vehicle (top) or anti-Ly6G depleting antibody (bottom).

(g) Left: Experimental approach for treatment with CXCR1/2 dual inhibitor ladarixin in transgene induced EY mice. Right: Representative images of CD45^+^ and Ly6G^+^ immune infiltrates in EY mouse tongue epithelia 20 days after transgene induction after treatment with vehicle (Left) or ladarixin (Right).

Panels b-e were analyzed by ANOVA with Tukey correction for multiple comparisons. Boxplots show median, interquartile range (IQR), and range. *p<0.05, **p<0.01

**Related to Fig. 5**.

**Extended Data Figure 9. YAP-mediated transcriptional activation of mTOR signaling**

(a) GSEA enrichment plots for the HALLMARK_MTORC1_SIGNALING gene set for E-, Y-, and EY-specific differentially expressed genes. Right: GSEA normalized enrichment scores and statistics for E-, Y-, and EY-specific differentially expressed genes.

(b) Western blot showing effect of siRNA-mediated knockdown of YAP, TAZ, and AXL on YAP target gene abundance and the abundance and phosphorylation of EGFR and S6 ribosomal protein in Cal33 cells.

(c) IGV tracks of YAP CUT&Tag, H3K27ac CUT&Tag, and ATACseq peaks at the *Nrg1* gene locus. Dotted line box indicates segment displayed in Fig. 6f.

(d) *Epgn* expression in transgenic tongue epithelia by RNAseq.

(e) IGV tracks of YAP CUT&Tag, H3K27ac CUT&Tag, and ATACseq peaks at the *Epgn* gene locus.

(f) IGV tracks of YAP CUT&Tag, H3K27ac CUT&Tag, and ATACseq peaks at the *Ereg* gene locus.

(g) The effects of siRNA-mediated knockdown of *NRG1*, *EREG*, *EPGN* on phospho- and total HER3, and S6 expression in HN12 whole cell lysate.

(h) The effects of siRNA-mediated knockdown of *NRG1*, *EREG*, *EPGN* on *NRG1*, *EREG*, *EPGN* transcript expression in Cal27 and HN12 cells. *EPGN* expression was undetectable in Cal27 cells (not shown).

(i) Representative wells (left) and quantification (right) of clonogenic assays in HN12 cells treated with DMSO, R428, INK128, or Erlotinib (Erlo).

(j) Representative wells (left) and quantification (right) of clonogenic assays in EY primary cultured cells cells treated with siRNA targeting, *YAP+TAZ* or *TEAD1+TEAD4*

(k) Representative wells (left) and quantification (right) of clonogenic assays in EY primary cultured cells cells treated with the mTOR inhibitor rapamycin or the YAP-TEAD inhibitor VT104. For panels c, e, f black bars indicate significant peaks. Red bars indicate EY-gained peaks.

For panels d, i, j, k boxplots show median, interquartile range (IQR), and range; ANOVA with Tukey correction for multiple comparisons. *p<0.05, **p<0.01, ***p<0.001, ****p<0.0001.

**Related to Fig. 6**.

**Extended Data Figure 10. TIC programs are enriched in HNSC and associated with disease-free survival**

(a) Related to Figure 7i. EY-module enrichment in malignant tumors (T) compared to matched normal solid tissues (N) by single sample GSEA among subjects in The Cancer Genome Atlas (TCGA) Head and Neck Squamous Carcinoma (HNSC) cohort (n=43 subjects with matched T and N samples). Two-tailed paired T-test: **p<0.01, ****p<0.0001.

(b-d) Kaplan-Meier plots demonstrating worse disease-free survival (n=393) among TCGA-HNSC subjects with greater median enrichment for the (b) G1/S, (c) pEMT, and (d) Focal Adhesion EY-modules.

**Related to Fig. 7**.

## SUPPLEMENTARY TABLES

**Supplementary Table 1**. Differentially expressed genes on bulk RNAseq

**Supplementary Table 2**. EY-unique DEGs and gene ontology analysis

**Supplementary Table 3**. Gene sets used for GSEA analyses

**Supplementary Table 4**. EY activated YAP transcriptional targets

**Supplementary Table 5**. Epithelial cluster marker genes

**Supplementary Table 6**. Immune cluster marker genes

**Supplementary Table 7**. EY-cluster WGCNA module genes and gene ontology analysis

## ONLINE METHODS

### Genotyping PCR primers

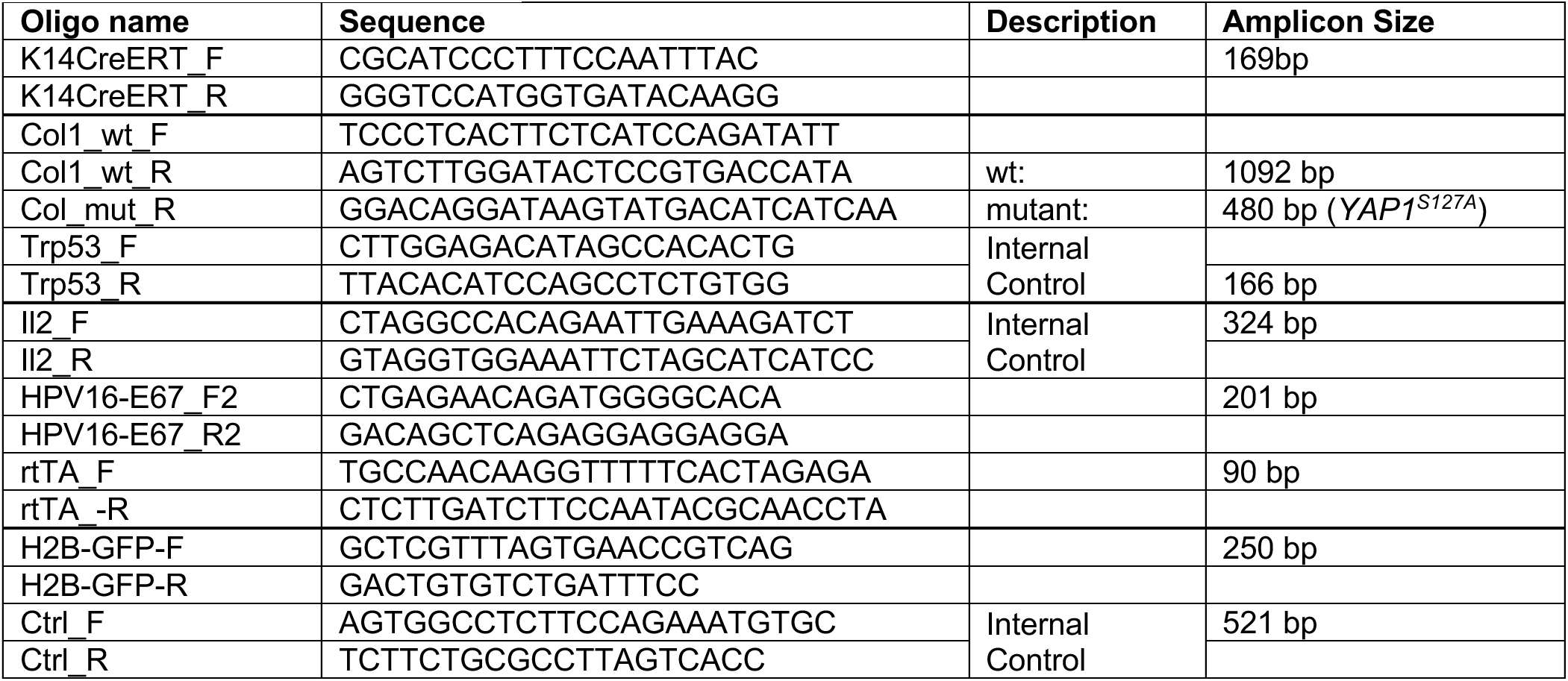

### Loxp–STOP–Loxp excision assay PCR primers

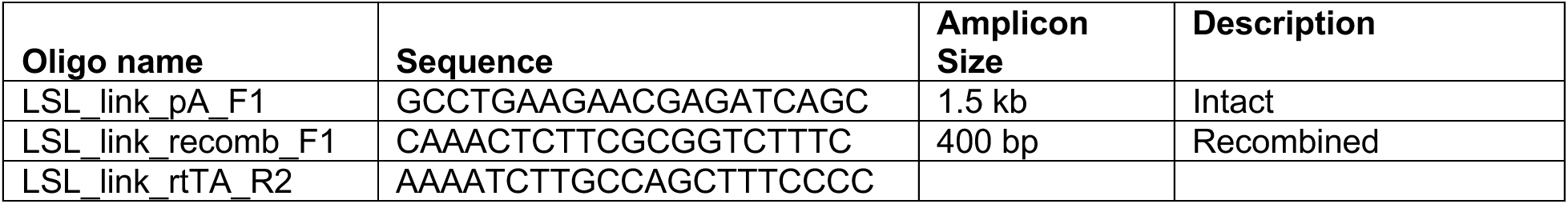

### RT-qPCR primers

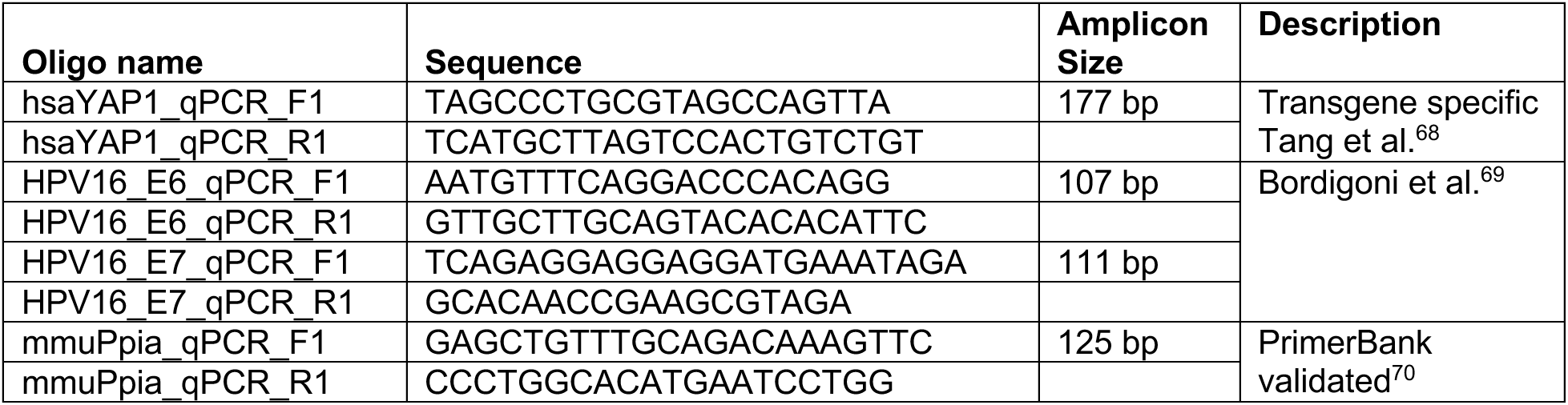

### IHC antibodies

| Primary Antibody | Clone | Dilution | Species | Supplier |
| --- | --- | --- | --- | --- |
| Pan-cytokeratin | ab9377 | 1/200 | Rabbit | Abcam |
| phospho-S6 | CST2211 | 1/400 | Rabbit | Cell Signaling Technology |
| KI67 | ab15580 | 1/400 | Rabbit | Abcam |
| P63 | CST39692 | 1/900 | Rabbit | Cell Signaling Technology |
| SOX2 | CST14962 | 1/300 | Rabbit | Cell Signaling Technology |

| Secondary Antibody | Dilution | Catalog # | Supplier |
| --- | --- | --- | --- |
| Goat Anti-Rabbit IgG Antibody (H+L),<br>Biotinylated | 1/200 | BA-1000 | Vector Laboratories |

### Immunofluorescence antibodies

| Primary Antibody | Clone | Dilution | Species | Supplier | Secondary Antibody | Secondary dilution | Secondary catalog # |
| --- | --- | --- | --- | --- | --- | --- | --- |
| KRT14 | poly19053 | 1/200 | Rabbit | BioLegend | AF568 goat anti-rabbit | 1/1000 | Thermo A11036 |
| PDPN-biotin | 8.1.1 | 1/100 | Syrian Hamster | BioLegend | AF647 streptavidin | 1/1000 | Thermo A78962 |
| KRT15 | Poly18339 | 1/100 | Chicken | BioLegend | AF674 goat anti-chicken | 1/1000 | Thermo A11036 |
| KI67 | ab15580 (poly) | 1/200 | Rabbit | Abcam | AF568 goat anti-rabbit | 1/1000 | Thermo A32933 |
| ITGA6 | GoH3 | 1/200 | Rat | BioLegend | AF647 goat-anti-rat | 1/1000 | Thermo A21247 |
| P63 | D9L7L | 1/200 | Rabbit | CST | AF568 goat anti-rabbit | 1/1000 | Thermo A32933 |
| IBA1 | E4O4W | 1/200 | Rab | CST | AF568 goat anti-rabbit | 1/1000 | Thermo A32933 |
| LY6G | 1A8 | 1/100 | Rat | BioLegend | AF647 goat-anti-rat | 1/1000 | Thermo A21247 |
| Broad Spectrum Cytokeratin | poly ab86734 | 1/200 | Mouse | Abcam | AF488 goat-anti-mouse | 1/1000 | Thermo A21121 |

### siRNAs

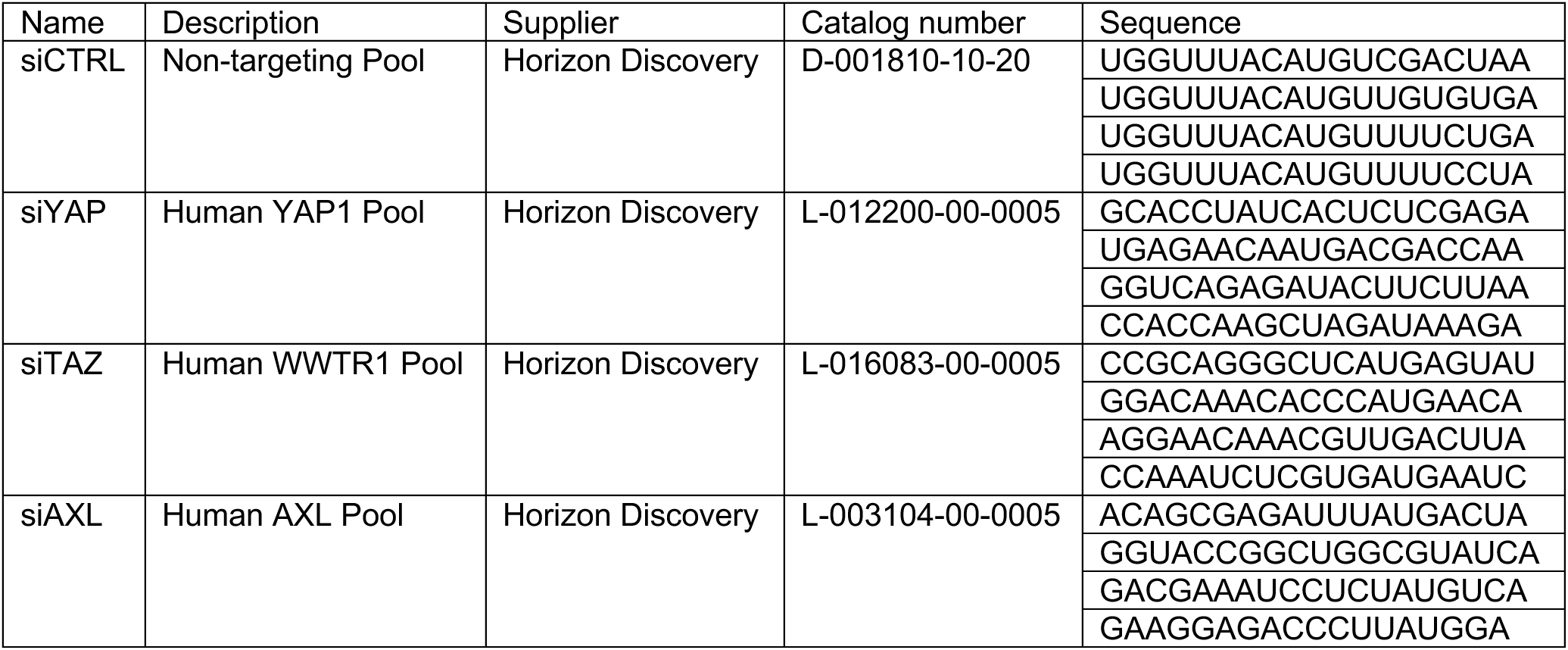

### Immunoblot antibodies

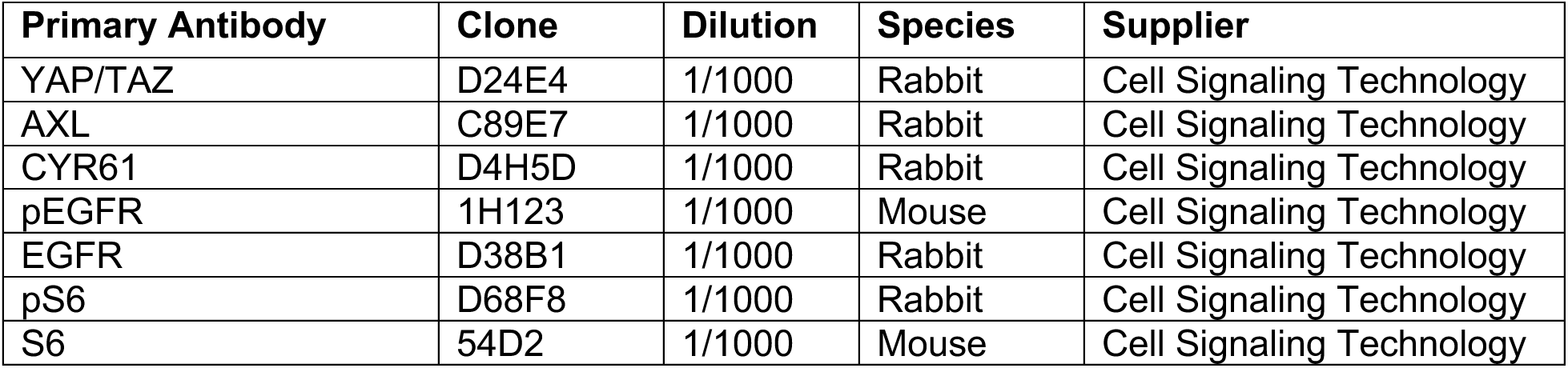
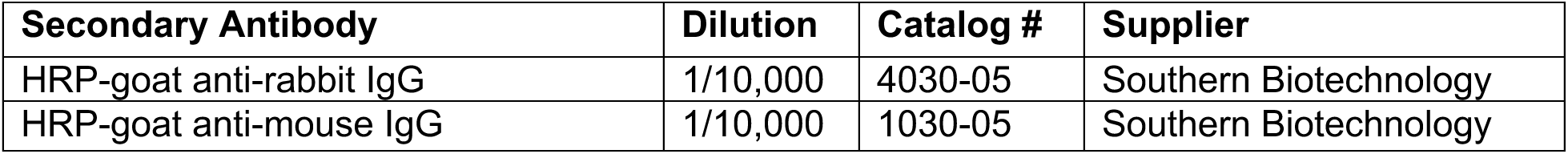

### CUT&Tag antibodies

| Primary Antibody | Cat no. / Clone | Dilution | Species | Supplier |
| --- | --- | --- | --- | --- |
| Isotype | DA1E | 1/50 | Rabbit | Cell Signaling Technology |
| YAP | D8H1X | 1/50 | Rabbit | Cell Signaling Technology |
| H3K27ac | D5E4 | 1/50 | Rabbit | Cell Signaling Technology |
| H3K27me3 (pAb) | Cat no. 39157 | 1/50 | Rabbit | Active Motif |

## EXPERIMENTAL MODEL AND SUBJECT DETAILS

### Mouse lines

Mice were housed in accordance with University of California San Diego Institutional Animal Care and Use Committee (IACUC) guidelines. The UCSD IACUC approved all mouse experiments (Protocol S15195). The following mouse lines were kindly provided by Dr. Elaine Fuchs (The Rockefeller University): Tg(KRT14-cre/ERT)^20Efu^ and Tg(tetO-HIST1H2BJ/GFP)^47Efu^.^29,30^ The Col1a1tm1(tetO-Yap1*)^Lrsn^ mouse was kindly provided by Dr. Fernando Camargo (Harvard University).^22^ The B6.Cg-Gt(ROSA)26Sor^tm1(rtTA,EGFP)Nagy/J^ mouse was obtained from The Jackson Laboratory.^71^ The Tg(tetO-HPV16-E6E7)^SGu^ mouse was designed by the Gutkind laboratory and generated in house.^21^ All transgenic mouse experiments were performed in age- and sex-balanced groups of 8-16 week old littermates. NSG™ mice (NOD.Cg-*Prkdc^scid^ Il2rg^tm1Wjl^*^/SzJ^) mice were originally obtained from The Jackson Laboratory and propagated at the Moores Cancer Center. Implantation of transgene epithelial cell suspensions were performed in 8-week-old female NSG mice.

### Human HNSC cell lines

CAL27 and CAL33 cell lines were obtained from the NIDCR Oral and Pharyngeal Cancer Branch cell collection.^72^ Cell identity was confirmed by STR profiling. CAL27 (CVCL_1107) was derived from a 56 year old male with tongue adenosquamous carcinoma. CAL33 (CVCL_1108) was derived from a 69 year old male with tongue squamous cell carcinoma. Both cell lines were cultured in DMEM (D-6429, Sigma-Aldrich, St. Louis, MO), 10% fetal bovine serum, 5% CO_2_, at 37 °C, and both tested free of Mycoplasma infection directly prior to experimentation.

### TCGA-HNSC

Transcriptome profiling, biospecimens, and clinical data from The Cancer Genome Atlas Program (TCGA) was downloaded from the National Cancer Institute GDC Data Portal for patients with the cancer type head and neck squamous cell carcinoma (HNSC) (https://portal.gdc.cancer.gov/projects/TCGA-HNSC)]^7^ Additional clinical data for this project was downloaded from cBioPortal (https://www.cbioportal.org/study/summary?id=hnsc_tcga_pan_can_atlas_2018).^73,74^

## METHOD DETAILS

### Husbandry and genotyping

We bred mice expressing E6-E7 (*Krt14-CreERT/LSL-rtTA/tetON_H2B-GFP/tetON_E6-E7*, “**E**”), *YAP1^S127A^* (*Krt14-CreERT/LSL-rtTA/tetON_H2B-GFP/tetON_YAP1^S127A^*, “**Y**”), or both transgenes (*Krt14-CreERT/LSL-rtTA/tetON_H2B-GFP/tetON_E6-E7/tetON_YAP1^S127A^*, “**EY**”). Littermates that bore neither *tetON_E6-E7* nor *tetON_YAP1^S127A^*effector transgenes but possessed the *Krt14-CreERT* and *LSL-rtTA* regulatory transgenes were used as the normal condition (*Krt14-CreERT/LSL-rtTA/tetON_H2B-GFP*, “**N**”). Intralingual injection of tamoxifen was performed to achieve reliable transgene induction. Mice were started on a doxycycline-containing diet on the first day of tamoxifen treatment. This treatment regimen resulted in consistent CreERT-mediated excision of the floxed STOP cassette and expression of effector and reporter transgenes in KRT14^+^ basal cells.

*Krt14-CreERT^+/+^/LSL-rtTA^+/+^/H2B-GFP^+/+^/E6-E7^+/-^*mice were crossed to *Krt14-CreERT^+/+^/LSL-rtTA^+/+^/H2B-GFP^+/+^/YAP1^S127A+/-^* resulting in Mendelian proportions of N, E, Y, and EY littermates. At 3-4 weeks of age, a tail fragment was obtained for initial screening genotype confirmation. Directly prior to transgene induction for experiments, mice were assigned to age and sex-balanced groups, and an ear fragment was obtained for confirmatory genotyping. Genomic DNA was isolated by incubating tissue in 25mM NaOH and 0.2 mM EDTA at 100°C for 1 hour, followed by neutralization with an equal volume of 40 mM Tris-HCl (pH 5.5).^75^ Multiplex polymerase chain reaction (PCR)-based genotyping was performed using REDTaq® polymerase per manufacturer recommendations (Millipore Sigma). Oligonucleotides were multiplexed as follows: (1) *LSL-rtTA* and *E6-E7* and *Il2* (positive control), (2) *Yap1^S127A^* and *Trp53* (positive control), (3) *Krt14-CreERT* and *Il2* (positive control). All PCR products were subjected to electrophoresis on 2% agarose gel in Tris acetate EDTA buffer.

### Transgene induction

Mice were anesthetized with isoflurane and 100µL of tamoxifen solution (20 mg/mL in miglyol) was administered into the tongue under stereomicroscopic visualization. One dose of tamoxifen was administered every other day for a total of 3 doses.

### Epithelia isolation

After *in situ* infiltration with 500uL collagenase+dispase solution (1mg/mL, 2.5 mg/mL) (Millipore Sigma), the tongues of euthanized mice were dissected free and incubated for 30 minutes at 37°C. The tongue epithelium was then dissected free from the underlying muscle under stereomicroscopic visualization.

### LoxP–STOP–LoxP excision assay

For floxed stop cassette excision assay, high-quality genomic DNA was isolated from whole epithelia using the DNeasy Blood and Tissue Kit per manufacturer protocol (Qiagen). PCR products were generated using REDTaq® polymerase and LSL excision primers, and were subjected to electrophoresis on 2% agarose gel in Tris acetate EDTA buffer.

### RT-qPCR

RNA was prepared by homogenization of whole tongue epithelia in TRIzol® (Invitrogen) followed by phenol:chloroform extraction and RNeasy Mini Kt based column purification with on-column DNase treatment (Qiagen). For quantitative PCR (qPCR), cDNA library preparation was performed using Bio-Rad iScript™ reverse transcriptase and qPCR was performed using Applied Biosystems Fast SYBR® Green Master Mix per manufacturer’s instructions.

### Evaluation of gross tongue lesions

Following transgene induction, mice were examined under anesthesia using a stereomicroscope every 3-7 days for the appearance of tongue epithelial lesions. Lesion free survival in days was defined as the time from first tamoxifen treatment to the appearance of the first gross lesion.

### Histopathology and immunohistochemical staining

Mouse tongues were transected from the pharynx and floor of mouth, and placed in 10% aqueous buffered zinc formalin for 24-36 hours at room temperature and transferred to 70% ethanol. Tissues were paraffin embedded, sectioned (5µm), and stained with hematoxylin and eosin (H&E) by standard protocols (Histoserv or LJI histology). H&E slides were prepared according to a standard protocol. https://www.protocols.io/view/hematoxylin-amp-eosin-protocol-for-leica-st5020-au-x54v9mozqg3e/v1

Immunohistochemistry (IHC) was performed as previously described.^76^ After deparaffinization, antigen retrieval was performed using IHC Antigen Retrieval Solution (ThermoFisher, 00-4955-58) in a steamer for 40 minutes. Endogenous peroxidases were inactivated using Bloxall Blocking Solution (Vector Labs, SP-6000, 30-min incubation, room temperature). Tissues were incubated with primary antibody overnight at 4°C then exposed to biotinylated anti-rabbit secondary antibody (Vector Labs, BA-1000, 1:400 dilution, 30 min at room temperature followed by avidin-biotin complex formation (Vector Laboratories, # PK-6100), staining with DAB substrate (Vector Laboratories, # SK-4105), and hematoxylin counterstain (Mayer’s Hematoxylin solution (Sigma MHS1-100ML)). All H&E and IHC stained slides were scanned using the Leica Aperio AT2 slide scanner at 40x magnification.

### Fluorescence microscopy

For whole mount fluorescence imaging, epithelial sheets were isolated from mice euthanized 36 hours after a single dose of intralingual tamoxifen, washed in HBSS, stained with NucBlue for 3 hours at room temperature, washed again, mounted immersed in HBSS between two cover glasses, and Z-stacks were acquired with a confocal microscope.

For cross-sectional fluorescence imaging, immediately after euthanasia, mice underwent intracardiac perfusion first with 2mM EDTA in PBS followed by 1.6% paraformaldehyde in PBS. Perfusion fixed tongues were dissected and incubated in 1.6% paraformaldehyde at room temperature overnight, then transferred to 30% sucrose in PBS for 2-3 days at 4°C, then washed in PBS, then embedded in OCT media and snap frozen in cryomolds for frozen section slide preparation. For fluorescent analyses, slides were thawed in the dark, blocked, incubated overnight at 4°C with primary antibodies, and then incubated with fluorophore-conjugated secondary antibodies for 2 hours at room temperature. Nuclei were then stained with Hoechst 33342 in PBS for 15 minutes and slides were mounted with ProLong Diamond mounting medium.

### Second harmonic generation for collagen imaging

The second-harmonic generation imaging was done on an upright Leica SP8 microscope with a resonant scanner and hybrid non-descanned detectors. Ti-Sapphire femtosecond pulsed Chameleon Ultra II (Coherent Inc.) laser was tuned to 855 nm and the beam was focused on the sample with an HC PL APO CS 10x/0.40 dry objective. The light was routed to the detectors with 560 nm, 495 nm, and 640 nm long-pass dichroic mirrors. The SHG signal was recorded with a 425/26 nm bandpass filter, the autofluorescence was recorded with a 650/60 nm bandpass filter. The pixel size was set to 0.746 µm, and 16x line averaging was used to improve the signal-to-noise ratio. Data were digitized in an 8-bit mode. The sample navigator software module was used to create autofocus support points and individual fields of view were tiled and stitched.

### RNAseq

Tongue epithelia were isolated and RNA was prepared as described above. RNA samples passing purity, concentration, and integrity quality metrics by NanoDrop and TapeStation were submitted to Novogene for oligo-dT-based mRNA selection, cDNA library preparation, and sequencing on Illumina NovaSeq6000.

### siRNA transfection in human cell lines

All human cells were transfected at 60% confluency using Lipofectamine RNAiMAX reagent according to the manufacturer’s instructions, using 20nM of each siRNA. Culture media was refreshed at 24 hours after transfection. Cells were placed under serum free conditions at 48 hours, and collected for experimentation at 72 hours post-transfection.

### Immunoblot assay

Cells rinsed with ice cold PBS and lysates were harvested in RIPA buffer (50 mM Tris-HCl, 150 mM NaCl, 1 mM EDTA, 1% NP-40) supplemented with Halt^TM^ Protease and Phosphatase Inhibitor Cocktail (#78440, ThermoFisher Scientific) and cleared by centrifugation for 15 minutes. The concentration of supernatants was measured using Bradford colorimetric assay. Equal amounts of protein were loaded onto 10% polyacrylamide gels, subjected to electrophoresis in Tris/Glycine/SDS buffer, and transferred to PVDF membranes. The membranes were blocked with 5% milk in TBS with 0.1% Tween-20 (TBS-T) buffer for 1 hour, incubated with primary antibodies diluted in 5% BSA overnight at 4°C. After washing 3 times with TBS-T, the membranes were incubated with HRP-conjugated secondary antibodies diluted in 5% milk in TBS-T for 1 h at room temperature. Immobilon Western Chemiluminescent HRP substrate (Millipore, MA) was used for detection.

### Cytokine array

Whole epithelia were homogenized in RIPA lysis buffer with protease and phosphatase inhibitors, snap frozen, and sent to Eve Technologies for Mouse Cytokine/Chemokine 44-Plex and Mouse MMP 5-plex Discovery Assay® Arrays.

### Generation of epithelial cell suspensions

Isolated epithelia were minced in 0.25% trypsin-EDTA (Thermo) and subjected to mechanical dissociation in the gentleMACS dissociator C tubes (Miltenyi #130-095-937) for 12 minutes at 37°C, followed by inactivation of trypsin and filtration.

### Primary epithelial cell culture

Mouse tongue epithelial cells were isolated from mice following transgene induction as described above. Cells were grown on collagen coated plates in complete DermaCult keratinocyte basal expansion medium (STEMCELL Technologies). Medium contained the manufacturer’s provided supplements, plus 5 ng/mL mouse EGF (Gibco), 50 pM cholera toxin (Sigma), 1x antibiotic/antimycotic solution [Gibco], and 2 uM doxycycline hyclate (Sigma, to maintain transgene activation) at 37°C with 5% CO_2_.

### Flow cytometry

Epithelial cell suspensions were stained for viability using LIVE/DEAD™ Fixable Blue Dead Cell Stain Kit (Thermo #L23105) and BUV737 Rat Anti-Mouse CD45 Clone 30-F11 (BD Biosciences #568344) and analyzed using a 5-laser Cytek Aurora.

### Cell sorting

EY epithelia were isolated and maintained in culture as previously above. When the cells were approximately 70-80% confluent, single cell suspensions were generated by subjecting the cells to EDTA and then trypsin, and then mechanically lifting the cells. Cells were counted and viability assessed by trypan blue staining using a Countess III cell counter. The cells were then resuspended in HBSS at a concentration of ~10 million cells/mL and subjected to fluorescence activated cell sorting (FACS) using an Aria II cell sorter. Single cells were identified based on forward and side scatter parameters, and then GFP positive and negative cells were sorted into individual tubes with cell culture medium. Sorted cells were returned to culture and expanded for experimentation and cryopreservation.

### scRNAseq

Single cell suspensions were generated from tongue epithelia, sorted for viability, and subjected to droplet-based single cell cDNA library preparation and sequencing. Two mice per genotype were used to generate tongue epithelial cell suspensions. Cells were sorted on a BD FACSAria-II. Viable single cells were selected by size (FSC x SSC) and viability (double negative for propridium iodide and Fixable Viability Dye eFluor780 (eBioscience) staining) parameters. Sorted cells were then loaded on a Chromium Controller (10x Genomics) using the Chromium Next GEM Single Cell 3’ Kit v3.1 (10x #1000269) according to the manufacturer’s protocol with a target of 10,000 cells per GEM reaction. The resulting cDNA library was sequenced on an Illumina NovaSeq 6000 using the S1 100 cycle Reagent Kit v1.5 (Illumina 200228319), with a targeted read depth of 20,000 reads/cell.

### CUT&Tag Sample Preparation, Data Processing, and Analysis

CUT&Tag assay and library preparation were performed on cell suspensions of cultured primary EY and N cells. Briefly, primary cells cultured on collagen coated plates were sequentially treated with 1 mM PBS-EDTA and 0.25% trypsin-EDTA (Gibco) to generate single cell suspensions. Cell suspensions were counted and viability assessed; 500,000 viable cells were input per condition. Cells were further processed using the CUT&Tag-IT™ Assay Kit (Active Motif, catalog no. 53160) following manufacturer’s specifications without deviation. See *CUT&Tag antibodies* table for antibody specifications. Raw reads were aligned using Bowtie2 (version 2.2.5) to build version mm10 of the mouse genome.^77^ Peaks were called independently in each replicate against the corresponding IgG isotype control using SEACR^78,79^ (version 1.3) in relaxed mode. Peaks with RPKM < 10 were filtered out.^80^ Consensus peaks were merged for each genotype, EY or N, by combining all filtered peaks using bedtools merge (version 2.27.1).^81^ Tornado plots were generated using deeptools (version 3.3.5).^82^ Differential acetylation was called using DESeq2 (version 1.42.0) and apeglm (version 1.24.0) in R (version 4.3.2).^83,84^ Peaks with adjusted p-values less than 0.05 were considered significant. Motif enrichment was performed using the findMotifsGenome.pl script in the HOMER package (version 4.11).^85^ Peaks were annotated using the annotatePeaks.pl script in the HOMER package. Peaks were annotated if they lie within the gene body or closer than 10 kb to the annotated TSS.

### ATAC-seq Sample Preparation, Data Processing, and Analysis

ATACseq assay and library preparation were performed on cell suspensions of cultured primary EY and N cells. Briefly, primary cells cultured on collagen coated plates were sequentially treated with 1 mM PBS-EDTA and 0.25% trypsin-EDTA (Gibco) to generate single cell suspensions. Cell suspensions were counted and viability assessed; 100,000 viable cells were input per condition. Cells were further processed using the ATAC-Seq Kit Assay Kit (Active Motif, catalog no. 53150) following manufacturer’s specifications without deviation. Raw reads were aligned using BWA (version 0.7.17) to build version mm10 of the mouse genome.^86^ Peaks were called using MACS2 (version 2.2.7.1) in narrow peak mode with a False Discovery Rate threshold of less than 0.01.^87^ Consensus peaks were merged for all samples by combining all called peaks using bedtools merge (version 2.27.1). Reads were recounted in consensus peaks using bedtools coverage (version 2.27.1). DESeq2 (version 1.42.0) and apeglm (version 1.24.0) in R (version 4.3.2) were used to call differential chromatin accessibility, peaks with adjusted p-value of less than 0.05 were considered significant. Motifs and peak annotation was performed as with the CUT&Tag data using HOMER.

### Orthotopic implantation

Epithelial cell suspensions were generated as described above. Cell count and viability was performed using trypan blue on the Countess III. Cells were only implanted for epithelia that showed minimum viability of 75%. After one wash in HBSS, 2×10^5^ viable cells were implanted orthotopically into the tongues of NSG mice. After implantation, mice tongues were first evaluated at 5 days after implantation then every other day until endpoint was reached.

### Imaging equipment and software

Gross evaluation of tongue lesions (Fig. 1 & 5) was performed using the Motic K-400P stereo microscope. Fluorescent cross-sectional images of were acquired using the Zeiss LSM780 confocal microscope system with Zeiss Black software (Fig. 2a, S1, S2, 6), Zeiss AxioZ1 slide scanner (Fig. 2c), or Zeiss LSM990 confocal microscope system with Zeiss Blue software (**Extended Data Fig. 5**). Histological images (H&E, IHC, immunofluorescence) were analyzed using QuPath 0.2.3, ImageJ/FIJI, or MATLAB.

## QUANTIFICATION AND STATISTICAL ANALYSIS

### Histopathological analyses

Histopathological changes for each experimental condition were independently evaluated by at least two board-certified veterinary pathologists (KK, KSakaguchi, AAM). Carcinoma was defined as atypical epithelial cells deep to the basement membrane. Average epithelial thickness was determined on mid-tongue axial sections by measuring 8-10 orthogonal lines from the basement membrane to the epithelial surface. Carcinoma burden was defined as the number of independent carcinoma foci identified in individual tongues. Carcinoma size was defined as the cross-sectional area of atypical epithelial cells invading deep to the basement membrane. Carcinoma depth of invasion was measured using a line orthogonal to the basement membrane of the closest adjacent normal mucosa to the deepest point of tumor invasion.

### Quantification of IHC

Basal phospho-S6 staining was quantified in QuPath using a trained pixel classifier applied to manually-segmented epithelial basal layer regions of interest (ROI). At least three ROIs each with a minimum area of 20,000 um^2^ were analyzed per sample. The fraction of phosphoS6 positive pixels for each sample was calculated using the mean of the ROIs after adjusting for relative area per ROI. Suprabasal Ki67, p63, Sox2+ nuclei were quantified in a similar fashion using a trained object classifier applied to manually-segmented epithelial subrabasal layer regions of interest. At least three suprabasal layer ROIs with minimum area of 50,000 um^2^ were selected per sample.The fraction of Ki67, p63, or Sox2+ positive nuclei for each sample was calculated using the mean of the ROIs after adjusting for relative area per ROI.

### IF nuclear segmentation

Instance segmentation was performed using Stardist, a deep-learning based segmentation FIJI plugin. Distinct grayscales values were assigned to each nucleus called by Stardist. MATLAB scripts were then developed to parse the label images, placing the linear indices corresponding to each pixel within a nucleus into a cell array. The relative size of each cell array corresponded to the number of segmented nuclei per image. The Hoechst and GFP channels from each confocal image were separated for independent segmentation and cell array formation. The ratio of GFP+ to Hoechst+ nuclei were calculated by comparing the resulting nuclear pixel cell arrays for the GFP and Hoechst channels for each image. The Hoechst channel was used for normalization and calculation of the mean fluorescence intensity (MFI) in the GFP channel. A threshold value was calculated using this normalized MFI to call Hoechst+/GFP+ and Hoechst+/GFP-nuclei.

### IF spatial context analysis

To determine the spatial fluorescent intensity distribution of ITGA6 as a function of distance from the basement membrane, MFI was calculated along the manually traced basement membrane by calculating vertical shifts from the basement membrane on a per-pixel basis. An array of each pixel location was created, and the MFI along the length of the traced basement membrane was calculated. As epithelial intensity distributions varied based on the length of the basement membrane tracing, they were interpolated using a spline with 50,000 query points, allowing the arrays to be combined and a normalized average intensity distribution to be calculated for each set of image arrays for a given mouse.

### Statistical analyses

Statistical analyses were performed in GraphPad Prism 9.5.1 with an alpha threshold of 0.05. Groupwise comparisons were tested using Kruskal-Wallis (one-way ANOVA on ranks) test with Dunn’s post-hoc correction for multiple comparison. Differences in survival were compared by Mantel-Cox Log-Rank test with Bonferroni correction for multiple comparisons. Pairwise comparisons between normal and malignant tumors was conducted using two-tailed paired t test. The correlation between mTOR and YAP signatures among TCGA tumors was determined initially by performing a simple linear regression and tested for significance using a two-tailed Spearman’s test. Additional statistical analyses on bulk RNAseq, scRNAseq, CUT&Tag, ATACseq data, and for multiomics analyses were performed in R (v4.1.2, 2021-11-01, “Bird Hippie”); see respective sections for details.

### Bulk RNAseq analyses

Paired-end reads were aligned using STAR v2.7.9 using default settings. STAR index was created using the GRCm39 primary genome FASTA and annotation files. The resulting BAM files were sorted by name using samtools v1.7 then gene counts were quantified using HTSeq-count v0.13.5. Pairwise differential expression was calculated and principal component analysis plots were created using DESeq2 v1.34.0. DEGs were defined at thresholds of p_adj_<0.01 and log_2_FC>1.

### GO and GSEA analyses

Gene ontology (GO) analyses were performed using GeneOntology.org (Panther 17.0) using significantly differentially expressed genes at (|log2FC| > 1, p-value < 0.01). Gene set enrichment analysis (GSEA^88^) was conducted using the Julia packages Match.jl and BioLab.jl, which contain bioinformatics and computational biology functions under active development. Prior to GSEA analyses, raw bulk RNAseq reads were aligned to the human reference transcriptome with the pseudo-aligner Kallisto^89^ using the “quant” command. Transcript expression values were normalized to transcript per million. Transcript expression was converted to gene expression using the maximum individual transcript expression. Single sample GSEA^57^ was performed with rank normalization against MSigDB^31,32^ gene set collections c2, c3, c5, and c6, with 10,000 permutations. Enriched and depleted gene sets were prioritized based on respective information coefficients^90–92^ and Bonferroni-corrected chi-square p-values.

### Global clustering

Single cell gene expression data was processed from the Illumina sequencer files using Cell Ranger (v5.0.0) and its prebuilt mouse reference genome. Individual sample data was then processed and merged using the Seurat (v4.3.0) SCTransform pipeline. Low quality cells (mitochondrial percentage >7, features <1000 and >5500, transcripts per cell >30,000) were filtered prior to data scaling and normalization. After filtering, data was transformed using SCTransform with default parameters, regressing on percent mitochondrial content. Principal component analysis was performed with RunPCA, using the top 50 PCs. Dimensionality reduction was performed with RunUMAP, using the top 30 dimensions. Nearest-neighbor analysis was performed using FindNeighbors using the top 30 dimensions and with k.param set to 50. Clustering was performed with FindClusters with resolution 0.3. Marker genes were calculated using FindAllMarkers with default parameters. Cluster identities were assigned by analysis of differential gene expression.

### Epithelial and immune cell sub-clustering

Epithelial and immune cell subset Seurat objects were generated and analyzed individually using Seurat version 4.3.0. Following assignment of cells to epithelial or immune cell subsets, the analysis pipeline used for the combined analysis was run again for each individual subset with identical parameters. Following initial subset clustering, contaminating residual immune or epithelial cells were removed from the subset Seurat objects, and the subset data reanalyzed in Seurat.

### Transgene alignment

For quantification of the transgene expression in single cells, STARsolo algorithm implemented to STAR aligner version 2.7.9a was applied. Briefly, the custom FASTA file was generated by merging the mm10 mouse genome and the transgene sequences. The index file for this custom genome was generated by STAR using the custom GTF file including the annotations for transgenes with the following parameters; *--sjdbOverhang 100*, *--genomeSAsparseD 3*. Subsequently, the FASTQ files including cDNA reads and cell barcodes of each sample were aligned to the custom genome by STARsolo with the following parameters; *--soloType CB_UMI_Simple*, *--clipAdapterType CellRanger4*, *--outFilterScoreMin 30*, *--soloCBmatchWLtype 1MM_multi_Nbase_pseudocounts*, *--soloUMIfiltering MultiGeneUMI_CR*, *-soloUMIdedup 1MM_CR*, *--soloCellFilter EmptyDrops_CR*. Finally, the cell-gene count arrays of the transgenes for each sample were obtained as the output of STARsolo. The data was imported to R and normalized with Seurat R package with “NormalizeData” function using the following options; *normalization.method = “LogNormalize”* and *scale.factor = 10000*. The normalized transgene expression arrays were merged with the Seurat object of the epithelial cell cluster by cell barcodes for downstream analysis.

### Weighted Gene Co-expression Network Analysis (WGCNA) of scRNAseq data

R package hdWGCNA version 0.2.03 (https://smorabit.github.io/hdWGCNA/) was used for WGCNA analysis in the scRNAseq dataset. Normalization of the integrated Seurat object containing cell-gene expression arrays of EY-genotype epithelial cells was performed using NormalizeMetacells using parameters k=10, max_shared=10, min_cells=20. A soft thresholding power was determined as 8 using the function TestSoftPowers and applied for estimation of co-expressing network in the EY-genotype scRNAseq dataset. Significantly co-expressed module genes and highly connected genes within each module (hub genes) were identified by computing eigengene-based connectivity (kME). The heatmap representing topology overlap matrix (TOM) of module genes was generated using R package ComplexHeatmap version 2.14.0. Genes with signed module eigengene-based connectivity measure (kME) greater 0.3 were considered as moderate to high confidence module genes. Modules were assigned functional annotations based on enrichment of member genes for biological processes using Enrichr^93,94^ and MetaScape.^95^

### Sequence: HPV16^E6-E7^

ATGCACCAAAAGAGAACTGCAATGTTTCAGGACCCACAGGAGCGACCCAGAAAGTTACCACAGTTATGCACAGAGCTGCAAACAACTATACATGATATAATATTAGAATGTGTGTACTGCAAGCAACAGTTACTGCGACGTGAGGTATATGACTTTGCTTTTCGGGATTTATGCATAGTATATAGAGATGGGAATCCATATGCTGTATGTGATAAATGTTTAAAGTTTTATTCTAAAATTAGTGAGTATAGACATTATTGTTATAGTTTGTATGGAACAACATTAGAACAGCAATACAACAAACCGTTGTGTGATTTGTTAATTAGGTGTATTAACTGTCAAAAGCCACTGTGTCCTGAAGAAAAGCAAAG ACATCTGGACAAAAAGCAAAGATTCCATAATATAAGGGGTCGGTGGACCGGTCGATGTATGTCTTGTTGCAGATCATCAAGAACACGTAGAGAAACCCAGCTGTAATCATGCATGGAGATACACCTACATTGCATGAATATATGTTAGATTTGCAACCAGAGACAACTGATCTCTACTGTTATGAGCAATTAAATGACAGCTCAGAGGAGGAGGATGAAATAGATGGTCCAGCTGGACAAGCAGAACCGGACAGAGCCCATTACAATATTGTAACCTTTTGTTGCAAGTGTGACTCTACGCTTCGGTTGTGCGTACAAAGCACACACGTAGACATTCGTACTTTGGAAGACCTGTTAATGGGCACACTAGGAATTGTGTGCCCCATCTGTTCTCAGAAACCATAA

### Sequence: YAP1^S127A^ (S127A codon underlined and in bold)

ATGGATCCCGGGCAGCAGCCGCCGCCTCAACCGGCCCCCCAGGGCCAAGGGCAGCCGCCTTCGCAGCCCCCGCAGGGGCAGGGCCCGCCGTCCGGACCCGGGCAACCGGCACCCGCGGCGACCCAGGCGGCGCCGCAGGCACCCCCCGCCGGGCATCAGATCGTGCACGTCCGCGGGGACTCGGAGACCGACCTGGAGGCGCTCTTCAACGCCGTCATGAACCCCAAGACGGCCAACGTGCCCCAGACCGTGCCCATGAGGCTCCGGAAGCTGCCCGACTCCTTCTTCAAGCCGCCGGAGCCCAAATCCCACTCCCGACAGGCCAGTACTGATGCAGGCACTGCAGGAGCCC TGACTCCACAGCATGTTCGAGCTCAT**<u>gCC</u>**TCTCCAGCTTCTCTGCAGTTGGGAGCTGTTTC TCCTGGGACACTGACCCCCACTGGAGTAGTCTCTGGCCCAGCAGCTACACCCACAGCTCAGCATCTTCGACAGTCTTCTTTTGAGATACCTGATGATGTACCTCTGCCAGCAGGTTGGGAGATGGCAAAGACATCTTCTGGTCAGAGATACTTCTTAAATCACATCGATCAGACAACAACATGGCAGGACCCCAGGAAGGCCATGCTGTCCCAGATGAACGTCACAGCCCCCACCAGTCCACCAGTGCAGCAGAATATGATGAACTCGGCTTCAGGTCCTCTTCCTGATGGATGGGAACAAGCCATGACTCAGGATGGAGAAATTTACTATATAAACCATAAGAACAAGACCACCTCTTGGCTAGACCCAAGGCTTGACCCTCGTTTTGCCATGAACCAGAGAATCAGTCAGAGTGCTCCAGTGAAACAGCCACCACCCCTGGCTCCCCAGAGCCCACAGGGAGGCGTCATGGGTGGCAGCAACTCCAACCAGCAGCAACAGATGCGACTGCAGCAACTGCAGATGGAGAAGGAGAGGCTGCGGCTGAAACAGCAAGAACTGCTTCGGCAGGCAATGCGGAATATCAATCCCAGCACAGCAAATTCTCCAAAATGTCAGGAGTTAGCCCTGCGTAGCCAGTTACCAACACTGGAGCAGGATGGTGGGACTCAAAATCCAGTGTCTTCTCCCGGGATGTCTCAGGAATTGAGAACAATGACGACCAATAGCTCAGATCCTTTCCTTAACAGTGGCACCTATCACTCTCGAGATGAGAGTACAGACAGTGGACTAAGCATGAGCAGCTACAGTGTCCCTCGAACCCCAGATGACTTCCTGAACAGTGTGGATGAGATGGATACAGGTGATACTATCAACCAAAGCACCCTGCCCTCACAGCAGAACCGTTTCCCAGACTACCTTGAAGCCATTCCTGGGACAAATGTGGACCTTGGAACACTGGAAGGAGATGGAATGAACATAGAAGGAGAGGAGCTGATGCCAAGTCTGCAGGAAGCTTTGAGTTCTGACATCCTTAATGACATGGAGTCTGTTTTGGCTGCCACCAAGCTAGATAAAGAAAGCTTTCTTACATGGTTA

## RESOURCE AVAILABILITY

### Materials availability

Transgenic mice and cell lines generated in this study are available from the Lead Contact upon reasonable request and completion of Material Transfer Agreements (MTA). There may restrictions to the availability of these reagents due to cost or limited quantities.

### Data Availability

The authors confirm that the source data underlying the findings are fully available. Bulk and single cell gene expression, ATAC sequencing, and CUT&Tag sequencing data are available in the NCBI Gene Expression Omnibus database (https://www.ncbi.nlm.nih.gov/geo) under the GEO series records: GSE276778 (ATACseq), GSE276779 (CUT&Tag for YAP, H3K27ac, and H3K27me3), GSE276781 (RNAseq, primary cells); GSE276782 (RNAseq, tissue), GSE276783 (scRNAseq).

## Notes

### Competing Interest Statement

J.S.G. has received other commercial research support from Kura Oncology, Mavupharma, Dracen, SpringWorks Therapeutics, is a consultant/advisory board member for Oncoceutics Inc., Vividion Therapeutics, Domain Therapeutics, and Pangea Therapeutics, and io9, and founder of Kadima Pharmaceuticals. The remaining authors declare no competing interests.

### Summary of Updates

1. We have isolated primary epithelial cells from our conditional YAPS127A/E6-E7 transgenic mice (EY cells; New Ext Data Fig 2f-h) and demonstrate their tumor initiating capacity. These cells were used to further dissect mechanisms underlying YAP-mediated tumor initiation through new epigenetic and integrative multiomic analyses (New Fig 3 and New Ext Data Fig 5), experiments evaluating the effects of mTOR inhibition (New Ext Data Fig 9j-k), and unbiased analyses of immune specific transcripts (New Ext Data Fig 7b-c & 8a-d). 2. In a first of its kind analysis, we have mapped YAP genome-wide localization to native chromatin using CUT&Tag, revealing ~38,000 YAP binding sites; ~11,000 gained with EY activation (New Fig 3a). 3. We have performed integrated multiomic analysis of RNAseq, YAP CUT&Tag, H3K27ac CUT&Tag, and ATACseq to identify key YAP-driven pathways in tumor initiating cells (New Fig 3). 4. We now show that YAP directly regulates expression of chemokines and cytokines mediating the recruitment of myeloid immune cells (G-MDSC) to the invasive front of EY-induced tumors, as part of the cancer-initiation process (New Ext Data Fig 8a). 5. We have depleted G-MDSCs using antibody and blocked key chemokines promoting G-MDSC recruitment (using small molecule inhibition of CXCR1/CXCR2) in vivo, obtaining direct evidence of a functional role for G-MDSC in tumor initiation (New Fig 5j and New Ext Data Fig 8g). 6. We have dissected the mechanistic link between YAP and mTOR in detail, unveiling a novel autocrine mechanism where YAP-dependent expression of NRG1 and consequent HER3/EGFR activation, converging with YAP-driven AXL expression to activate mTOR in tumor initiating cells (New Fig 6). 7. Importantly, we now show that mTOR inhibitor treatment diminishes tumor initiation in vivo (New Fig 6m-p), supporting the direct relevance of novel YAP-mTOR crosstalk in cancer initiation. 8. We found that YAP and mTOR pathway activation is enriched in malignant tumors compared to matched normal tissue, and YAP and mTOR activation are strongly correlated in human HNSC tumors and associated with poor survival (New Fig 7a-f).

