## Supplementary Figures 1-10 for "YAP-Driven Oral Epithelial Stem Cell Malignant Reprogramming at Single Cell Resolution"

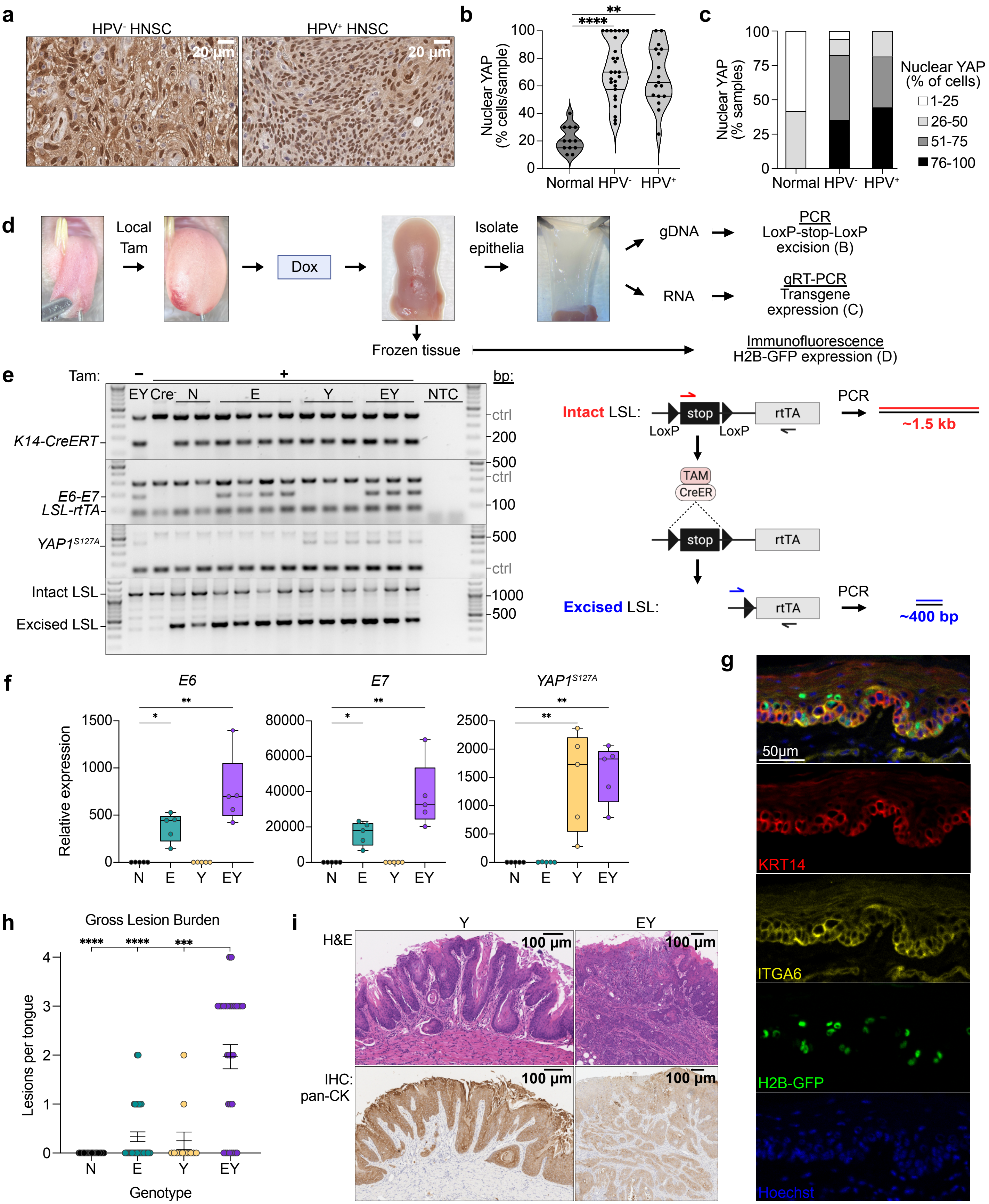
a
HPV- HNSC
HPV+ HNSC
20 µm
b
Nuclear YAP (% cells/sample)
Normal
HPV-
HPV+
c
Nuclear YAP (% samples)
Nuclear YAP (% of cells)
1-25
26-50
51-75
76-100
d
Local Tam
Dox
Isolate epithelia
gDNA
RNA
PCR
LoxP-stop-LoxP excision (B)
qRT-PCR
Transgene expression (C)
Frozen tissue
Immunofluorescence
H2B-GFP expression (D)
e
Tam:
EY
Cre-
N
E
Y
EY
NTC
bp:
K14-CreERT
E6-E7
LSL-rtTA
YAP1S127A
Intact LSL
Excised LSL
ctrl
Intact LSL:
stop
LoxP
rtTA
PCR
~1.5 kb
TAM
CreER
Excised LSL:
~400 bp
f
E6
E7
YAP1S127A
Relative expression
g
50µm
KRT14
ITGA6
H2B-GFP
Hoechst
h
Gross Lesion Burden
Lesions per tongue
Genotype
i
Y
EY
H&E
IHC: pan-CK
100 µm

**Extended Data Figure 1. Spatiotemporally controlled YAP and E6-E7 activation in OEPCs**
(a) Representative images of nuclear YAP protein in tissues by IHC in HPV negative (left) and HPV positive (right) malignant human HNSC tissue.
(b) Percent of cells with nuclear YAP in normal human tissue (n=12) and malignant HNSC (n=44) from tissue microarrays stratified by HPV status. Two-tailed Mann-Whitney test: **p<0.01, ****p<0.0001.
(c) Percent of samples with positive nuclear YAP in normal human tissues (n=12) and malignant HNSC (n=44) stratified by HPV status.
(d) Overview of local transgene induction in lingual epithelia and strategies for validation of transgene activation at genomic and transcript levels, and *in situ*.
(e) Left: Genomic DNA extracted from tongue epithelia 10 days after transgene induction with tamoxifen and doxycycline treatment. Top three gel images show presence of transgene and internal control (ctrl) PCR products. Bottom gel image confirms $Lox_p$-Stop-$Lox_p$ (LSL) recombination for *LSL-rtTA*. Lane 1 mouse tongue epithelial gDNA from a tamoxifen and doxycycline untreated mouse bearing all transgenes. NTC, no template control. Right: Schematic of oligonucleotide primer design to assay LSL excision.
(f) qRT-PCR quantification of $HPV^{E6}$, $HPV^{E7}$, and $YAP^{S127A}$ expression in tongue epithelia 10 days after transgene induction. Boxplots show median, interquartile range (IQR), and range.
(g) Immunofluorescence image of tongue epithelia from a Krt14-CreER$^+$LSL-rtTA$^+$tetON_H2B-GFP$^+$ mouse 6 days after transgene induction showing basal localization of KRT14 and ITGA6, and nuclear expression of H2B-GFP. Nuclei are counterstained with Hoescht.
(h) Total number of lesions per tongue 20 days after transgene induction. Means with SEM are shown.
(i) Hematoxylin and eosin (top) and pan-cytokeratin (pan-CK) IHC (bottom) stained tongue epithelia demonstrating infiltrative carcinoma in Y (left panels) and EY (right panels) mice.
**Related to Fig. 1.**

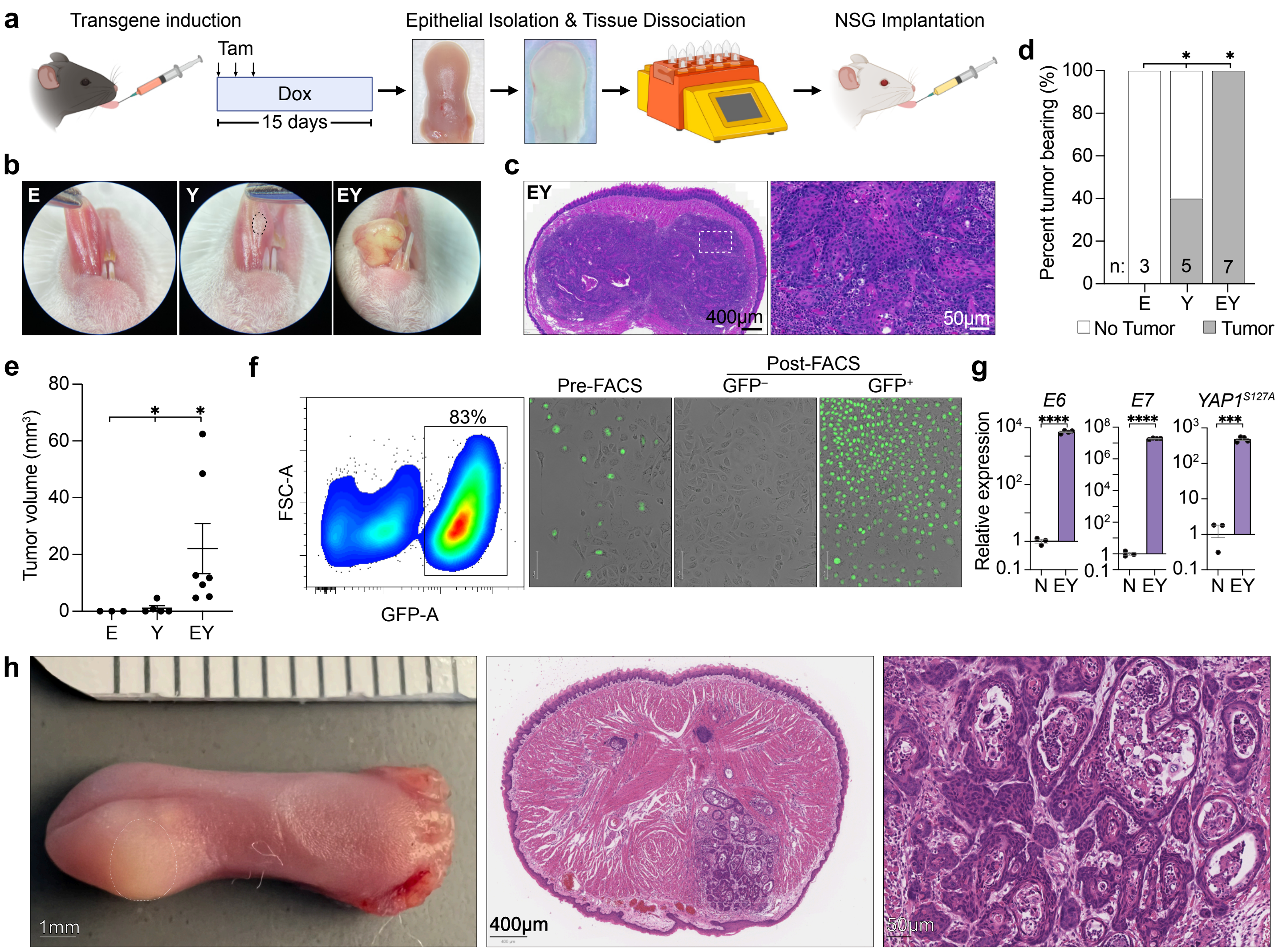

a
Transgene induction
Tam
Dox
15 days
Epithelial Isolation & Tissue Dissociation
NSG Implantation
b
E
Y
EY
c
EY
400µm
50µm
d
Percent tumor bearing (%)
100
80
60
40
20
0
n: 3
5
7
E
Y
EY
No Tumor
Tumor
e
Tumor volume (mm³)
80
60
40
20
0
E
Y
EY
f
83%
FSC-A
GFP-A
Pre-FACS
Post-FACS
GFP⁻
GFP⁺
g
E6
E7
YAP1 S127A
Relative expression
****
***
N EY
h
1mm
400µm
50µm

**Extended Data Figure 2. YAP and E6-E7 activation in OEPCs induces tumor initiating cells**
(a) Experimental approach for the generation and orthotopic implantation of transgene-induced epithelial cell suspensions.
(b) Representative photographs of NSG mouse tongues after orthotopic implantation transgene-induced epithelial cell suspensions.
(c) Representative H&E stained section of an NSG mouse tongue after orthotopic implantation with transgene-induced EY epithelial cell suspension.
(d) Implanted tumor outgrowth frequency; n, number of orthotopically implanted mouse tongues. Between group comparisons were made using Fisher's exact test with Bonferroni correction.
(e) Tumor volumes 10 days after implantation. Mean with SEM is shown. ANOVA with Tukey correction for multiple comparisons used for between group comparisons.
(f) Left: Gating strategy for FACS enrichment of H2B-GFP transgene positive cells derived from EY tongue epithelium based on GFP expression. Right: Representative fluorescence microscopy of primary EY cell cultures pre- and post-FACS enrichment.
(g) qRT-PCR quantification of $HPV^{E6}$, $HPV^{E7}$, and $YAP^{S127A}$ expression in FACS-purified N and EY cell cultures. Barplots show mean and SEM.
(h) Representative gross (left) and H&E stained sections (right) after implantation of 5,000 FACS-enriched EY cells in NSG mouse tongues.
For all panels with asterisks denoting significance: *$p<0.05$, **$p<0.01$, ***$p<0.001$, ****$p<0.0001$.
**Related to Fig. 1.**

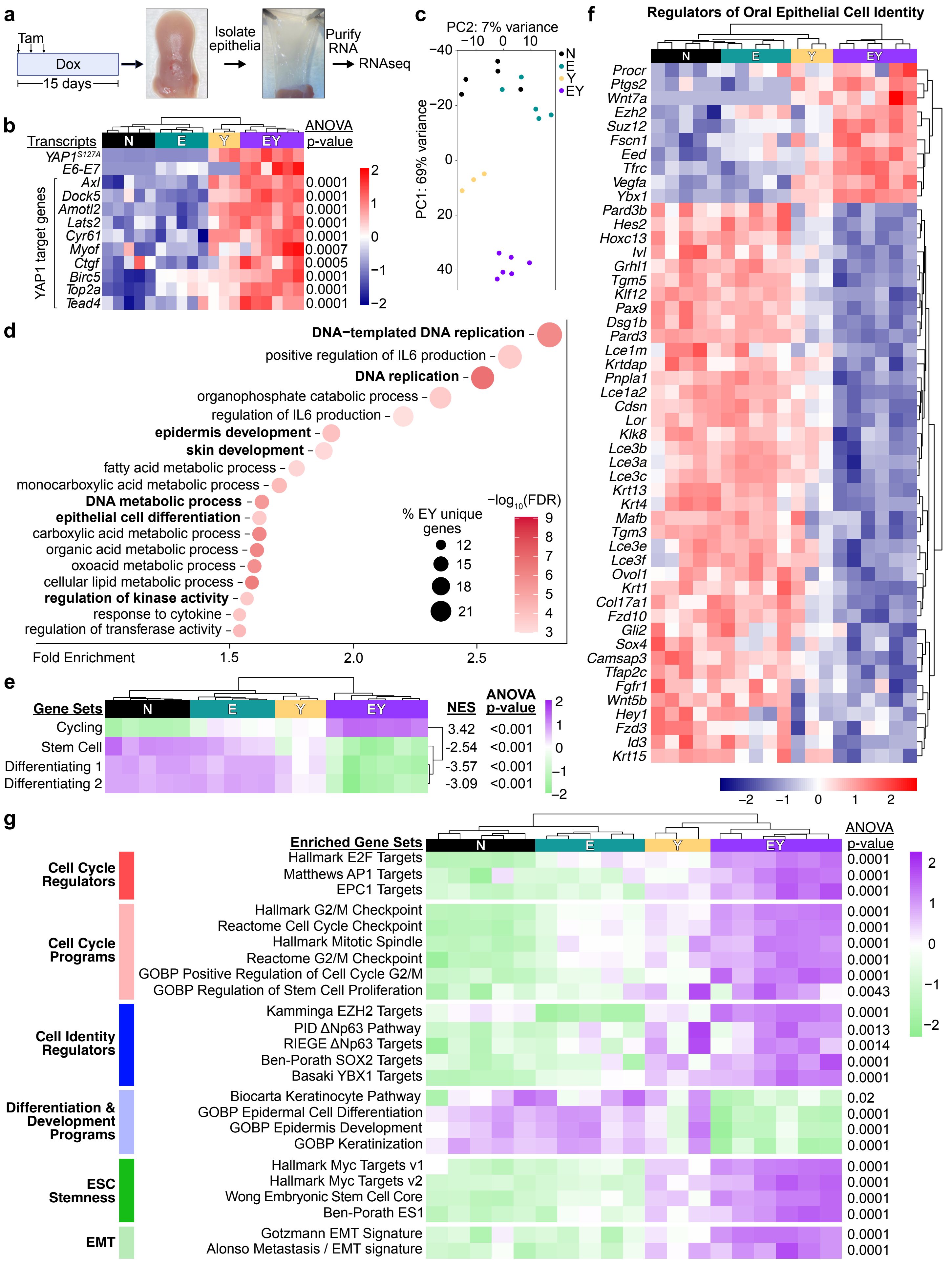

a
Tam
Dox
15 days
Isolate epithelia
Purify RNA
RNAseq
b
Transcripts
N
E
Y
EY
ANOVA p-value
YAP1S127A
E6-E7
YAP1 target genes
Axl 0.0001
Dock5 0.0001
Amotl2 0.0001
Lats2 0.0001
Cyr61 0.0001
Myof 0.0007
Ctgf 0.0005
Birc5 0.0001
Top2a 0.0001
Tead4 0.0001
c
PC2: 7% variance
PC1: 69% variance
d
DNA−templated DNA replication
positive regulation of IL6 production
DNA replication
organophosphate catabolic process
regulation of IL6 production
epidermis development
skin development
fatty acid metabolic process
monocarboxylic acid metabolic process
DNA metabolic process
epithelial cell differentiation
carboxylic acid metabolic process
organic acid metabolic process
oxoacid metabolic process
cellular lipid metabolic process
regulation of kinase activity
response to cytokine
regulation of transferase activity
Fold Enrichment
% EY unique genes
−log10(FDR)
e
Gene Sets
NES
ANOVA p-value
Cycling 3.42 <0.001
Stem Cell -2.54 <0.001
Differentiating 1 -3.57 <0.001
Differentiating 2 -3.09 <0.001
f
Regulators of Oral Epithelial Cell Identity
Procr
Ptgs2
Wnt7a
Ezh2
Suz12
Fscn1
Eed
Tfrc
Vegfa
Ybx1
Pard3b
Hes2
Hoxc13
Ivl
Grhl1
Tgm5
Klf12
Pax9
Dsg1b
Pard3
Lce1m
Krtdap
Pnpla1
Lce1a2
Cdsn
Lor
Klk8
Lce3b
Lce3a
Lce3c
Krt13
Krt4
Mafb
Tgm3
Lce3e
Lce3f
Ovol1
Krt1
Col17a1
Fzd10
Gli2
Sox4
Camsap3
Tfap2c
Fgfr1
Wnt5b
Hey1
Fzd3
Id3
Krt15
g
Enriched Gene Sets
ANOVA p-value
Cell Cycle Regulators
Hallmark E2F Targets 0.0001
Matthews AP1 Targets 0.0001
EPC1 Targets 0.0001
Cell Cycle Programs
Hallmark G2/M Checkpoint 0.0001
Reactome Cell Cycle Checkpoint 0.0001
Hallmark Mitotic Spindle 0.0001
Reactome G2/M Checkpoint 0.0001
GOBP Positive Regulation of Cell Cycle G2/M 0.0001
GOBP Regulation of Stem Cell Proliferation 0.0043
Cell Identity Regulators
Kamminga EZH2 Targets 0.0001
PID ΔNp63 Pathway 0.0013
RIEGE ΔNp63 Targets 0.0014
Ben-Porath SOX2 Targets 0.0001
Basaki YBX1 Targets 0.0001
Differentiation & Development Programs
Biocarta Keratinocyte Pathway 0.02
GOBP Epidermal Cell Differentiation 0.0001
GOBP Epidermis Development 0.0001
GOBP Keratinization 0.0001
ESC Stemness
Hallmark Myc Targets v1 0.0001
Hallmark Myc Targets v2 0.0001
Wong Embryonic Stem Cell Core 0.0001
Ben-Porath ES1 0.0001
EMT
Gotzmann EMT Signature 0.0001
Alonso Metastasis / EMT signature 0.0001

**Extended Data Figure 3. Oncogenic transcriptional reprogramming defines YAP and E6-E7 activated epithelia**
(a) Schematic demonstrating strategy for bulk RNAseq of microdissected tongue epithelia at 15 days post-transgene induction.
(b) *HPV*$^{E6E7}$, *YAP*$^{S127A}$, and YAP target gene expression in transgenic epithelia transcriptomes.
(c) Two-dimensional principal component analysis of individual N, E, Y, and EY transcriptome profiles.
(d) Enriched cellular processes among EY unique DEGs by Gene Ontology.
(e) Single sample GSEA showing enrichment or depletion of oral epithelial stem cell states including Cell Cycle Progression (G1/S, G2/M); Stem Cell; and Differentiation (Diff 1 and Diff 2). These gene signatures were derived from cluster defining genes from Jones et al. For gene set details, please see Supplementary Table 3.
(f) Relative expression of EY-unique DEGs regulating oral epithelial cell identity.
(g) Single sample GSEA for all transgenic conditions for gene signatures related to cell cycle, differentiation, stemness, and EMT. For gene set details, please see Supplementary Table 3.
Each row in panels b, e, f, and g was analyzed by ANOVA with Tukey correction for multiple comparisons: *$p<0.05$, **$p<0.01$, ***$p<0.001$, ****$p<0.0001$.
**Related to Fig. 2.**

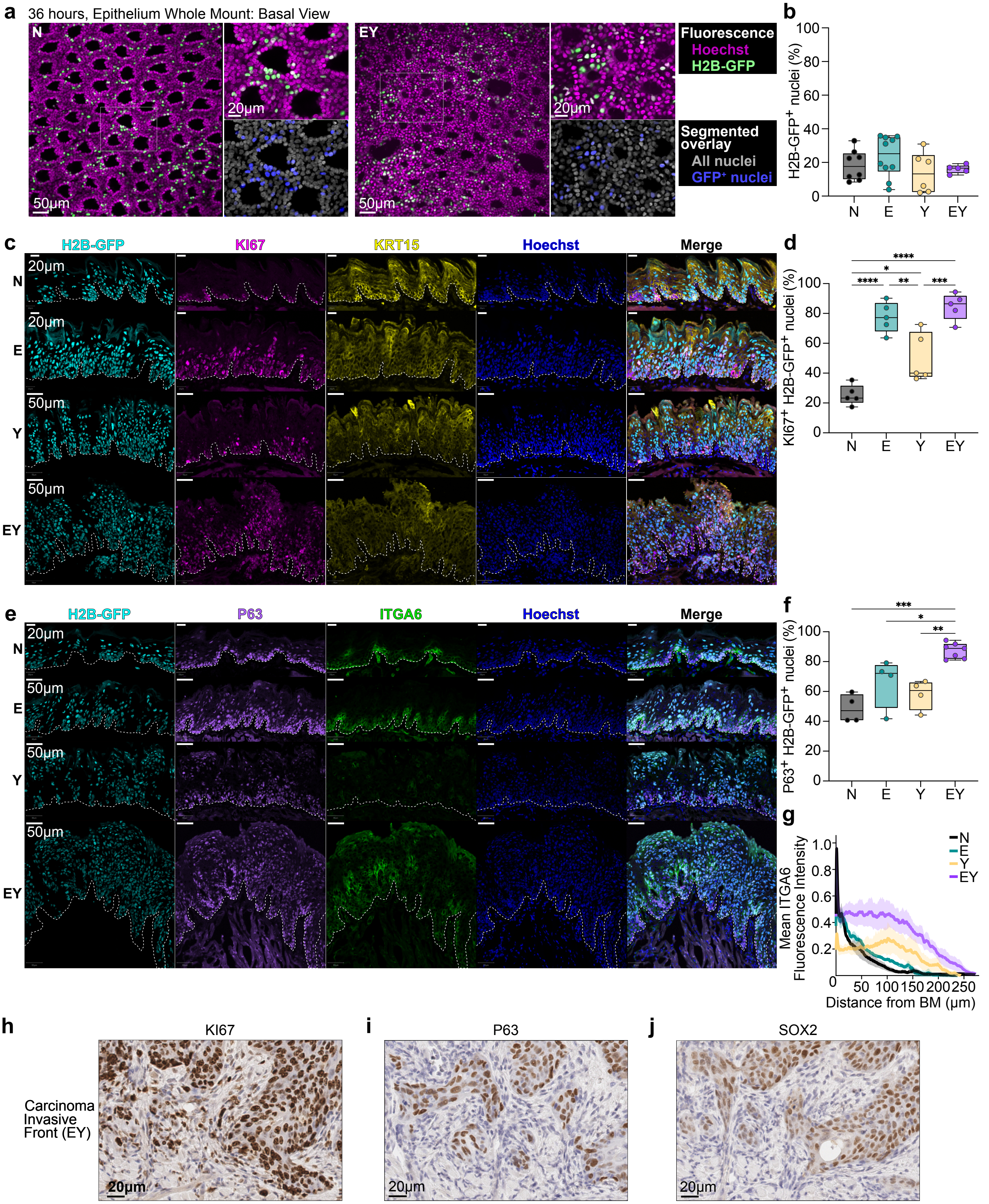
a
36 hours, Epithelium Whole Mount: Basal View
N
EY
50μm
20μm
Fluorescence
Hoechst
H2B-GFP
Segmented overlay
All nuclei
GFP+ nuclei
b
H2B-GFP+ nuclei (%)
N
E
Y
EY
c
H2B-GFP
KI67
KRT15
Hoechst
Merge
d
KI67+ H2B-GFP+ nuclei (%)
e
P63
ITGA6
f
P63+ H2B-GFP+ nuclei (%)
g
Mean ITGA6 Fluorescence Intensity
Distance from BM (μm)
h
KI67
Carcinoma Invasive Front (EY)
i
P63
j
SOX2

**Extended Data Fig. 4. Invasive carcinoma is preceded by the expansion of a stem-like cell population**

(a-b) Lineage tracing by fluorescent microscopy using the *H2B-GFP* reporter transgene to track and quantify $GFP^+$ nuclei. (a) Representative fluorescence images of and (b) percent basal H2B-$GFP^+$ nuclei in tongue epithelial basal layer whole mounts 36 hours after transgene induction.

(c) Representative immunofluorescence images of H2B-GFP, KI67, and KRT15 expression in tongue epithelia 15 days after transgene induction. White dashed line represents the basement membrane.

(d) Related to c. Percent $KI67^+GFP^+$ nuclei in tongue epithelia.

(e) Representative immunofluorescence images of H2B-GFP, P63, and ITGA6 expression in tongue epithelia 15 days after transgene induction. White dashed line represents the basement membrane.

(f) Related to e. Percent $P63^+GFP^+$ nuclei in tongue epithelia.

(g) Related to e. Quantitative spatial analysis showing normalized fluorescent intensity of ITGA6 signal from basement membrane to epidermal surface. Probability cloud shows 95% confidence interval.

(h-j) Magnified images of cells at the invasive front from Fig. 2m demonstrating (h) $KI67^+$ nuclei in Fig. 2i, (i) $P63^+$ nuclei in Fig. 2k, (j) $SOX2^+$ nuclei in Fig. 2m.

Panels b, d, and f were analyzed by ANOVA with Tukey correction for multiple comparisons: *$p<0.05$, **$p<0.01$, ***$p<0.001$, ****$p<0.0001$. Boxplots show median, interquartile range (IQR), and range.

**Related to Fig. 2.**

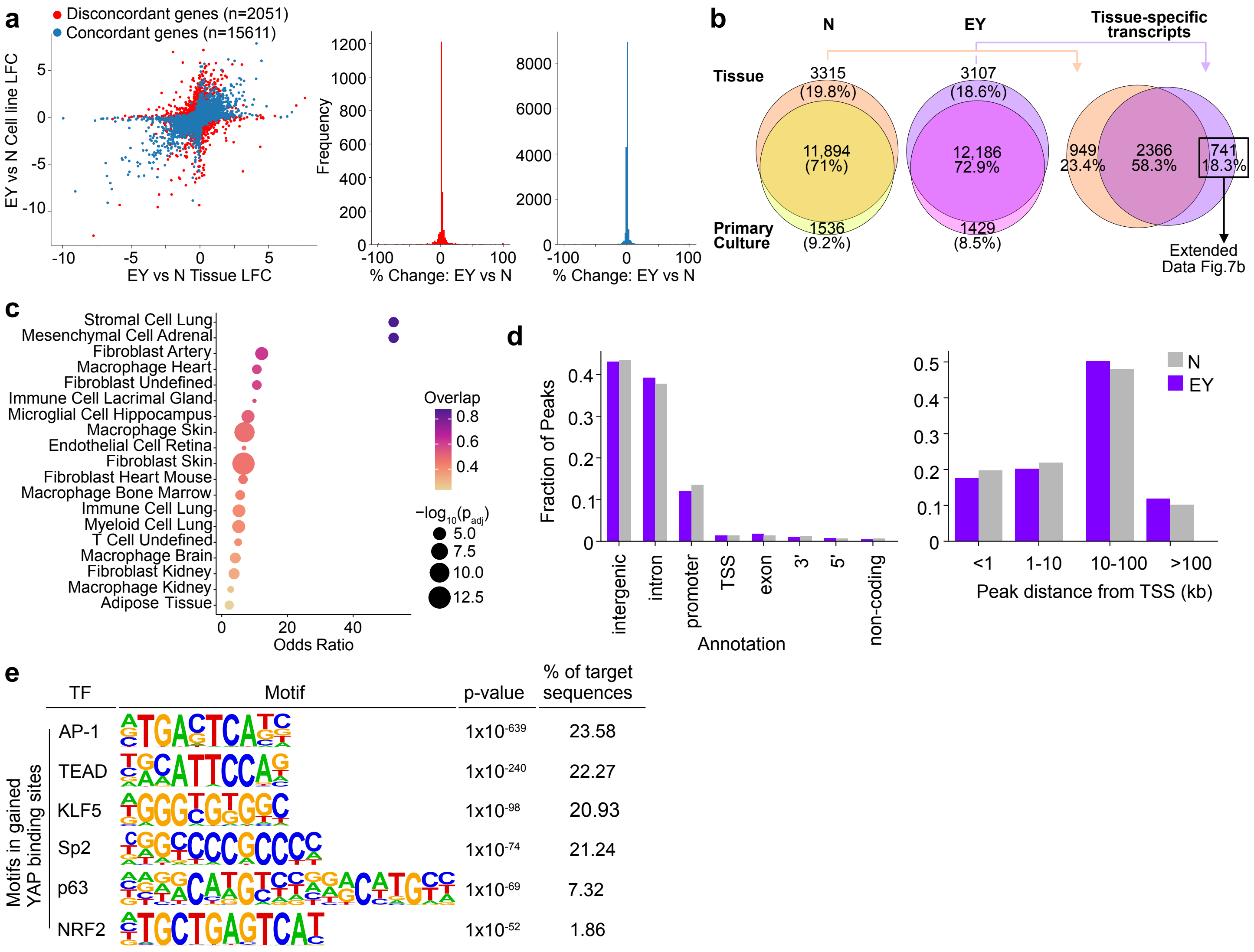
a
Disconcordant genes (n=2051)
Concordant genes (n=15611)
EY vs N Cell line LFC
EY vs N Tissue LFC
Frequency
% Change: EY vs N
% Change: EY vs N
b
N
EY
Tissue-specific transcripts
Tissue
3315 (19.8%)
11,894 (71%)
Primary Culture
1536 (9.2%)
3107 (18.6%)
12,186 72.9%
1429 (8.5%)
949 23.4%
2366 58.3%
741 18.3%
Extended Data Fig.7b
c
Stromal Cell Lung
Mesenchymal Cell Adrenal
Fibroblast Artery
Macrophage Heart
Fibroblast Undefined
Immune Cell Lacrimal Gland
Microglial Cell Hippocampus
Macrophage Skin
Endothelial Cell Retina
Fibroblast Skin
Fibroblast Heart Mouse
Macrophage Bone Marrow
Immune Cell Lung
Myeloid Cell Lung
T Cell Undefined
Macrophage Brain
Fibroblast Kidney
Macrophage Kidney
Adipose Tissue
Odds Ratio
Overlap
−log10(padj)
d
Fraction of Peaks
intergenic
intron
promoter
TSS
exon
3'
5'
non-coding
Annotation
N
EY
<1
1-10
10-100
>100
Peak distance from TSS (kb)
e
Motifs in gained YAP binding sites
TF
Motif
p-value
% of target sequences
AP-1 1x10-639 23.58
TEAD 1x10-240 22.27
KLF5 1x10-98 20.93
Sp2 1x10-74 21.24
p63 1x10-69 7.32
NRF2 1x10-52 1.86

**Extended Data Figure 5. Epigenetic reprogramming in YAP and E6-E7 activated epithelia and EY cell lines**

(a) Left: DESeq interaction model comparing genes enriched in EY compared to N tissue versus in EY compared to N primary culture cells. Discordant genes between epithelial tissue and cell lines are shown in red, and concordant genes (88%) in blue. Concordance threshold based on log fold difference, $p > 0.05$. Right: Frequency of genes with an observed difference (%) in EY versus N across EY and N tissues and cell lines.

(b) Left two Venn diagrams: overlap of transcripts detected in N (left) or EY (middle) epithelial tissue (top circles) versus primary cell culture (bottom circles). Right Venn diagram: overlap of tissue-specific N or EY transcripts identify 741 transcripts identified only in EY epithelial tissue.

(c) Enrichr analysis of the 2366 tissue-specific transcripts detected in both N and EY tissue found in the Cell Marker 2024 library.

(d) Fraction of YAP CUT&Tag peaks that localize to genomic region annotations defined using HOMER.

(e) Absolute distance of YAP CUT&Tag peaks to the nearest transcription start site (TSS).

(f) Top transcription factor binding motifs enriched in the EYgaine YAP CUT&Tag binding sites. P values were calculated using HOMER.

**Related to Fig. 3.**

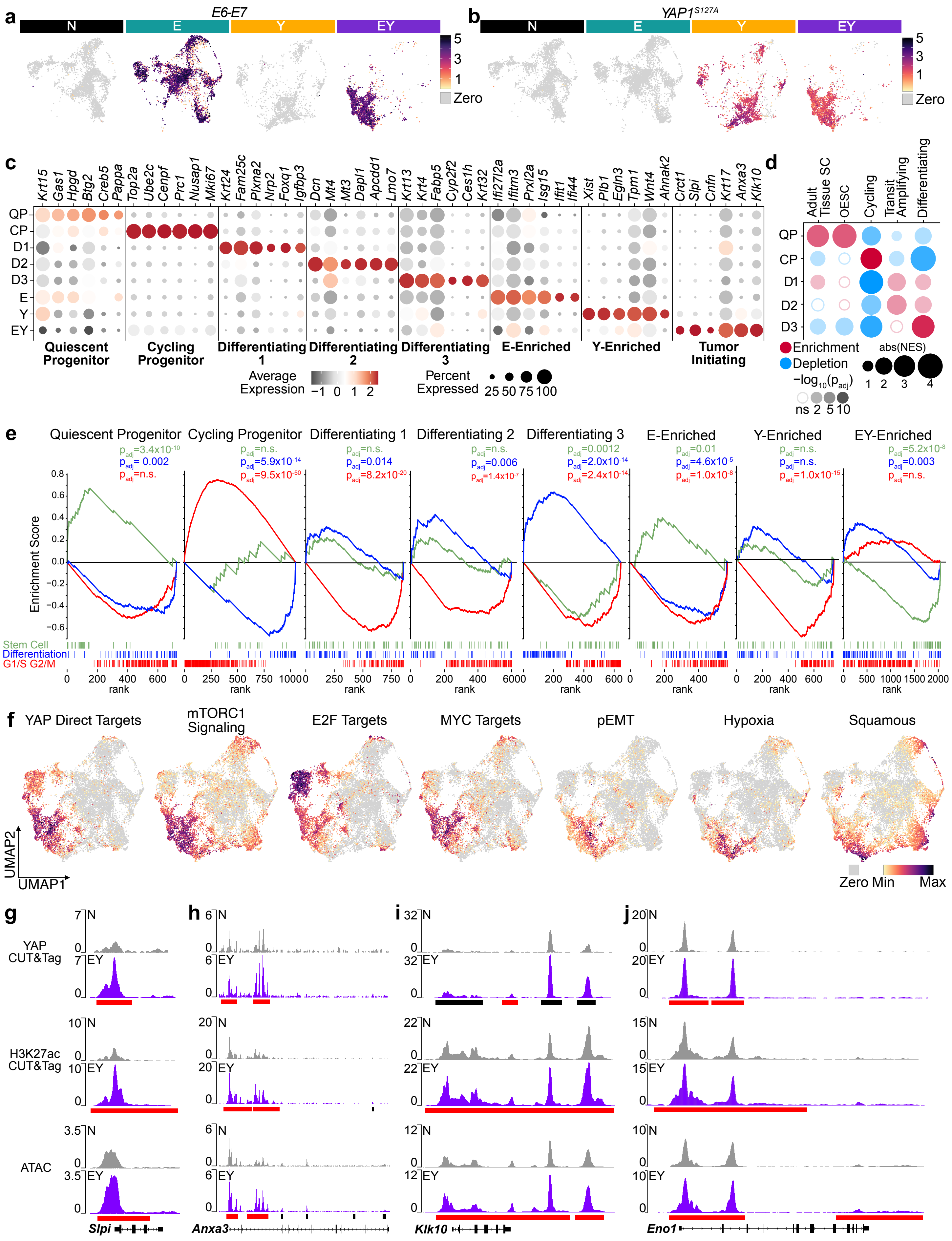

a
E6-E7
N
E
Y
EY
5
3
1
Zero
b
YAP1S127A
N
E
Y
EY
5
3
1
Zero
c
Krt15 Gas1 Hpgd Btg2 Creb5 Pappa Top2a Ube2c Cenpf Prc1 Nusap1 Mki67 Krt24 Fam25c Plxna2 Nrp2 Foxq1 Igfbp3 Dcn Mt4 Mt3 Dapl1 Apcdd1 Lmo7 Krt13 Krt4 Fabp5 Cyp2f2 Ces1h Krt32 Ifi27l2a Ifitm3 Prxl2a Isg15 Ifit1 Ifi44 Xist Plb1 Egln3 Tpm1 Wnt4 Ahnak2 Crct1 Slpi Cnfn Krt17 Anxa3 Klk10
QP
CP
D1
D2
D3
E
Y
EY
Quiescent Progenitor
Cycling Progenitor
Differentiating 1
Differentiating 2
Differentiating 3
E-Enriched
Y-Enriched
Tumor Initiating
Average Expression
−1 0 1 2
Percent Expressed
25 50 75 100
d
Adult Tissue SC
OESC
Cycling
Transit Amplifying
Differentiating
QP
CP
D1
D2
D3
Enrichment
Depletion
−log10(padj)
ns 2 5 10
abs(NES)
1 2 3 4
e
Quiescent Progenitor
Cycling Progenitor
Differentiating 1
Differentiating 2
Differentiating 3
E-Enriched
Y-Enriched
EY-Enriched
padj=3.4x10-10
padj= 0.002
padj=n.s.
padj=n.s.
padj=5.9x10-14
padj=9.5x10-50
padj=n.s.
padj=0.014
padj=8.2x10-20
padj=n.s.
padj=0.006
padj=1.4x10-7
padj=0.0012
padj=2.0x10-14
padj=2.4x10-14
padj=0.01
padj=4.6x10-5
padj=1.0x10-8
padj=n.s.
padj=n.s.
padj=1.0x10-15
padj=5.2x10-8
padj=0.003
padj=n.s.
Enrichment Score
0.8
0.6
0.4
0.2
0.0
−0.2
−0.4
−0.6
Stem Cell
Differentiation
G1/S G2/M
rank
f
YAP Direct Targets
mTORC1 Signaling
E2F Targets
MYC Targets
pEMT
Hypoxia
Squamous
UMAP2
UMAP1
Zero Min
Max
g
h
i
j
YAP CUT&Tag
H3K27ac CUT&Tag
ATAC
N
EY
Slpi
Anxa3
Klk10
Eno1

**Extended Data Figure 6. Single cell analysis of E6-E7 and YAP activated epithelial cells**
(a) *E6-E7* transgene expression stratified by genotype.
(b) *YAP1*$^{S127A}$ transgene expression stratified by genotype.
(c) Expression of the top 6 cluster defining genes among epithelial cell clusters. Dot size denotes the percentage of cells within a cluster expressing each transcript and the color indicates average gene expression across all cells in each cluster.
(d) GSEA of physiologic cell states across epithelial cell clusters. Circle color indicates enrichment (red) or depletion (blue). Circle size encodes the absolute value (abs) of the normalized enrichment score (NES). Circle opacity represents $-\log_{10}$ of the adjusted p-value ($p_{adj}$); circles are hollow if $p_{adj}>0.05$. For gene set details, please see Supplementary Table 3.
(e) GSEA enrichment plots for the Jones Stem Cell, Jones Differentiation, and Jones G1/S G2/M physiologic OEPC gene sets across epithelial cell clusters.
(f) Feature plots showing expression of the YAP Direct Target, mTORC1 Signaling, E2F Targets, MYC Targets, and the Barkley *et al.* recurring cancer cell state gene sets: partial epithelial to mesenchymal transition (pEMT), hypoxia, and squamous differentiation in single epithelial cells.
(g) IGV tracks of YAP CUT&Tag, H3K27ac CUT&Tag, and ATACseq peaks at the *Slpi* gene locus.
(h) IGV tracks of YAP CUT&Tag, H3K27ac CUT&Tag, and ATACseq peaks at the *Anxa3* gene locus.
(i) IGV tracks of YAP CUT&Tag, H3K27ac CUT&Tag, and ATACseq peaks at the *Klk10* gene locus.
(j) IGV tracks of YAP CUT&Tag, H3K27ac CUT&Tag, and ATACseq peaks at the *Eno1* gene locus.
For panels g-j: black bars indicate significant peaks. Red bars indicate EY-gained peaks.
**Related to Fig. 4.**

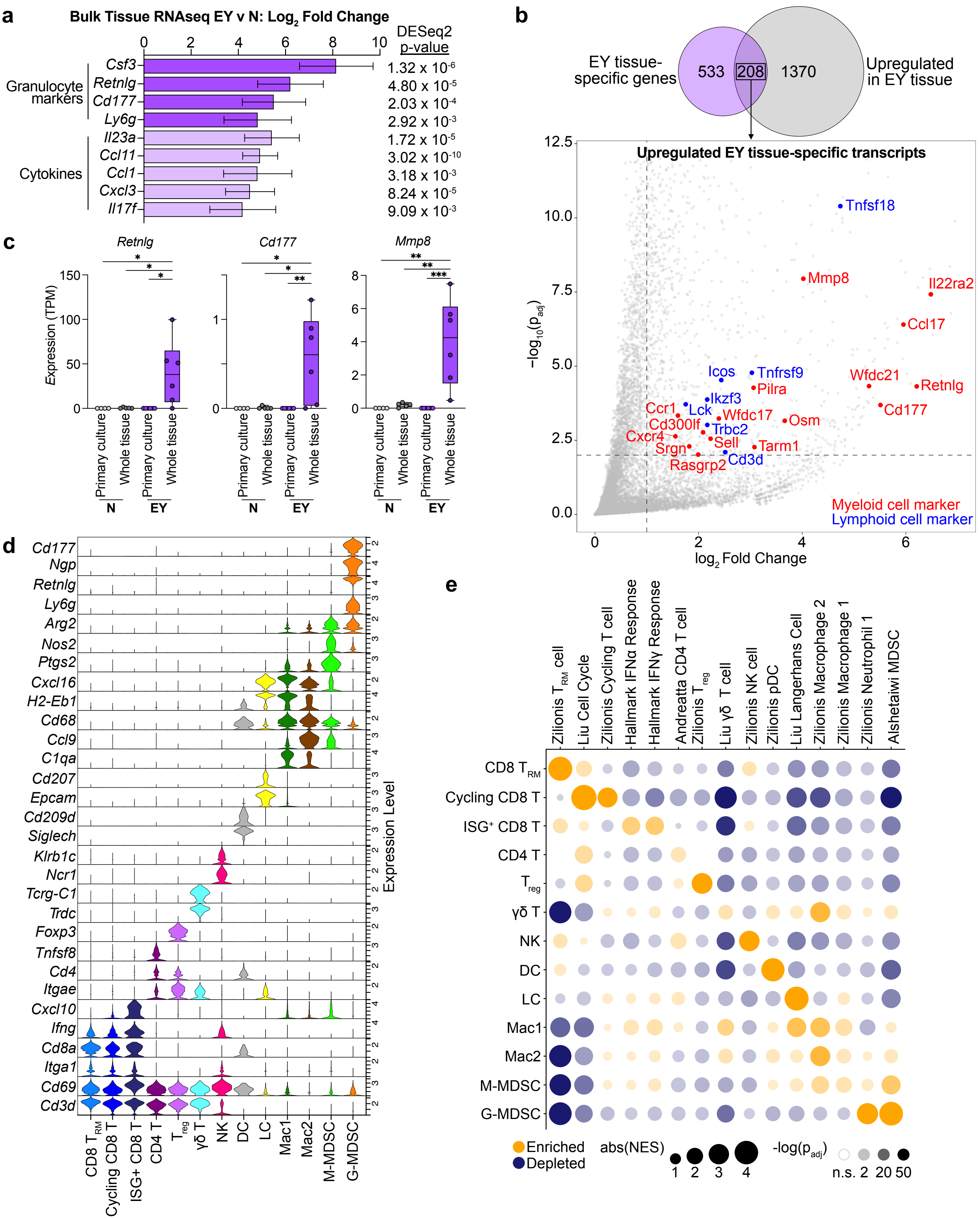
a
Bulk Tissue RNAseq EY v N: Log2 Fold Change
0 2 4 6 8 10
DESeq2 p-value
Granulocyte markers
Csf3
Retnlg
Cd177
Ly6g
Cytokines
Il23a
Ccl11
Ccl1
Cxcl3
Il17f
1.32 x 10-6
4.80 x 10-5
2.03 x 10-4
2.92 x 10-3
1.72 x 10-5
3.02 x 10-10
3.18 x 10-3
8.24 x 10-5
9.09 x 10-3
b
EY tissue-specific genes
533
208
1370
Upregulated in EY tissue
Upregulated EY tissue-specific transcripts
12.5
10.0
7.5
5.0
2.5
0.0
−log10(padj)
0 2 4 6
log2 Fold Change
Tnfsf18
Mmp8
Il22ra2
Ccl17
Wfdc21
Retnlg
Icos
Tnfrsf9
Pilra
Ikzf3
Cd177
Ccr1
Lck
Wfdc17
Osm
Cd300lf
Trbc2
Cxcr4
Sell
Tarm1
Srgn
Cd3d
Rasgrp2
Myeloid cell marker
Lymphoid cell marker
c
Retnlg
Cd177
Mmp8
Expression (TPM)
150
100
50
0
1
0
8
6
4
2
0
Primary culture
Whole tissue
Primary culture
Whole tissue
N
EY
d
Cd177
Ngp
Retnlg
Ly6g
Arg2
Nos2
Ptgs2
Cxcl16
H2-Eb1
Cd68
Ccl9
C1qa
Cd207
Epcam
Cd209d
Siglech
Klrb1c
Ncr1
Tcrg-C1
Trdc
Foxp3
Tnfsf8
Cd4
Itgae
Cxcl10
Ifng
Cd8a
Itga1
Cd69
Cd3d
Expression Level
CD8 TRM
Cycling CD8 T
ISG+ CD8 T
CD4 T
Treg
γδ T
NK
DC
LC
Mac1
Mac2
M-MDSC
G-MDSC
e
Zilionis TRM cell
Liu Cell Cycle
Zilionis Cycling T cell
Hallmark IFNα Response
Hallmark IFNγ Response
Andreatta CD4 T cell
Zilionis Treg
Liu γδ T cell
Zilionis NK cell
Zilionis pDC
Liu Langerhans Cell
Zilionis Macrophage 2
Zilionis Macrophage 1
Zilionis Neutrophil 1
Alshetaiwi MDSC
CD8 TRM
Cycling CD8 T
ISG+ CD8 T
CD4 T
Treg
γδ T
NK
DC
LC
Mac1
Mac2
M-MDSC
G-MDSC
Enriched
Depleted
abs(NES)
1 2 3 4
-log(padj)
n.s. 2 20 50

**Extended Data Figure 7. Single cell analysis of E6-E7 and YAP polarized epithelial immune infiltrate**

(a) RNAseq data from bulk epithelia for granulocyte markers and cytokines among EY DEGs. Means with SEM are shown.

(b) Top: Venn diagram showing overlap of EY tissue-specific and EY upregulated genes. Bottom: Volcano plot of epithelial bulk RNAseq data comparing EY vs N highlighting EY upregulated tissue-specific genes. Myeloid cell associated genes enriched in EY epithelial tissue are shown in red and lymphoid cell associated genes enriched in EY epithelial tissue are shown in blue.

(c) Bulk RNAseq expression in TPM in transcripts per million (TPM) of the EY upregulated tissue-specific myeloid cell markers, *Retnlg*, *Cd177*, and *Mmp8*, across N and EY samples, and epithelial tissue and primary culture conditions.

(d) Expression of select cell type defining genes among immune cells by scRNAseq.

(e) Dot Plot demonstrating GSEA of published cell-type defining gene lists across immune cell types. Dot color indicates enrichment (Orange) or depletion (Blue). Dot size represents the absolute value of the normalized enrichment score (NES). Dot opacity represents -$\log_{10}$ of the adjusted p-value. For gene set details, please see Supplementary Table 3.

**Related to Fig. 5.**

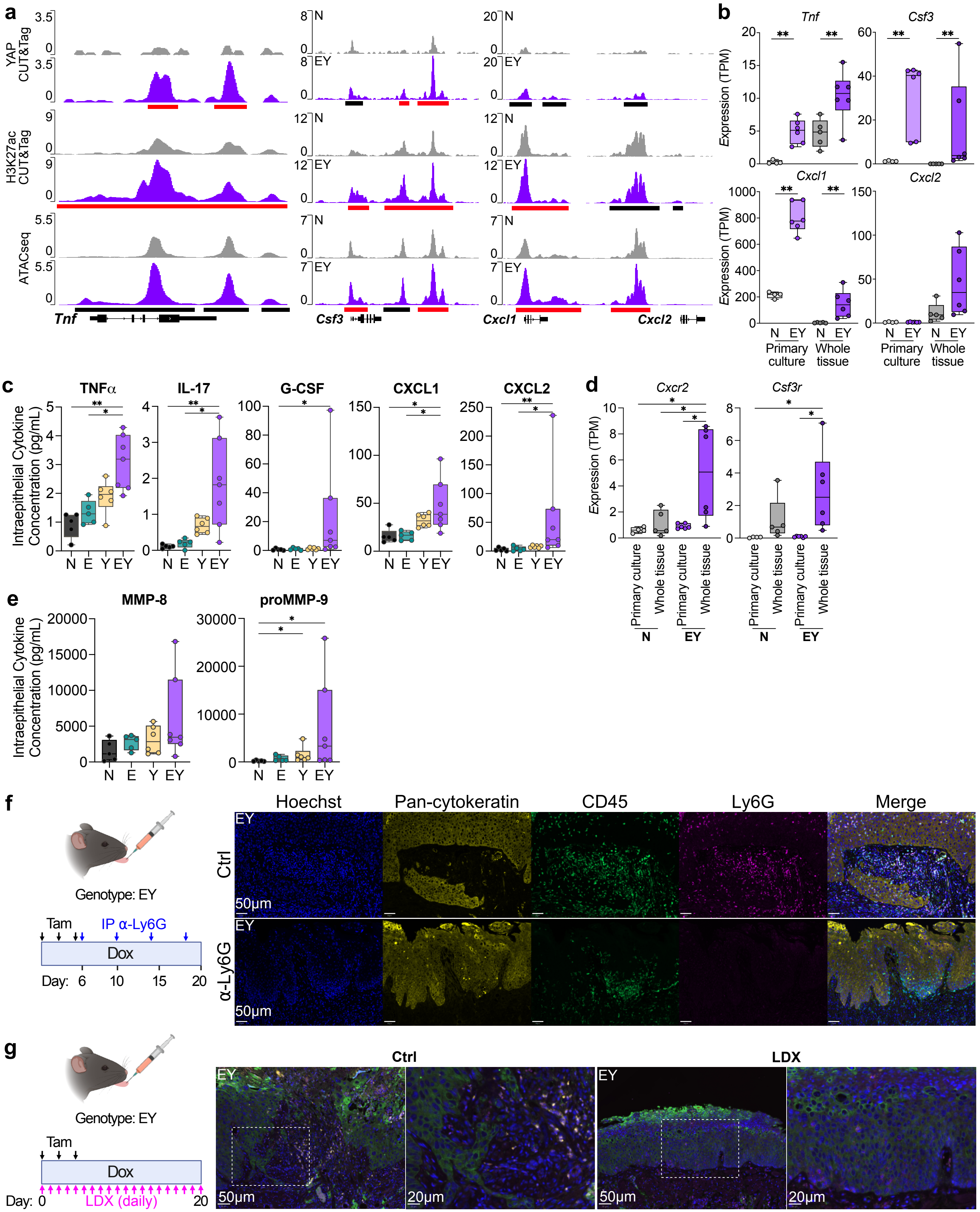

a
YAP CUT&Tag
H3K27ac CUT&Tag
ATACseq
N
EY
Tnf
Csf3
Cxcl1
Cxcl2
b
Tnf
Csf3
Cxcl1
Cxcl2
Expression (TPM)
N EY N EY
Primary culture
Whole tissue
c
TNFα
IL-17
G-CSF
CXCL1
CXCL2
Intraepithelial Cytokine Concentration (pg/mL)
N E Y EY
d
Cxcr2
Csf3r
Expression (TPM)
Primary culture
Whole tissue
N
EY
e
MMP-8
proMMP-9
Intraepithelial Cytokine Concentration (pg/mL)
N E Y EY
f
Genotype: EY
Tam
IP α-Ly6G
Dox
Day: 6 10 15 20
Hoechst
Pan-cytokeratin
CD45
Ly6G
Merge
Ctrl
α-Ly6G
EY
50μm
g
Genotype: EY
Tam
Dox
Day: 0 20
LDX (daily)
Ctrl
LDX
EY
50μm
20μm

**Extended Data Figure 8.**
(a) IGV tracks of YAP CUT&Tag, H3K27ac CUT&Tag, and ATACseq peaks at the *Tnf, Csf3, Cxcl1, Cxcl2* gene loci. Black bars indicate significant peaks. Red bars indicate EY-gained peaks.
(b) Bulk RNAseq expression in TPM of *Tnf, Csf3, Cxcl1, Cxcl2* across N and EY samples, and epithelial tissue and primary culture conditions.
(c) G-CSF, IL-17, TNFα, CXCL1, and CXCL2 protein abundance in epithelial lysates from N, E, Y, and EY mice 15 days after transgene induction.
(d) Bulk RNAseq expression in TPM of *Cxcr2, Csf3r* across N and EY samples, and epithelial tissue and primary culture conditions.
(e) MMP8 and pro-MMP9 protein abundance in epithelial lysates from N, E, Y, and EY mice 15 days after transgene induction.
(f) Left: Experimental approach for depletion of $LY6G^+$ G-MDSCs in transgene induced EY mice. Right: Representative images of $CD45^+$ and $Ly6G^+$ immune infiltrates in EY mouse tongue epithelia 20 days after transgene induction after treatment with vehicle (top) or anti-Ly6G depleting antibody (bottom).
(g) Left: Experimental approach for treatment with CXCR1/2 dual inhibitor ladarixin in transgene induced EY mice. Right: Representative images of $CD45^+$ and $Ly6G^+$ immune infiltrates in EY mouse tongue epithelia 20 days after transgene induction after treatment with vehicle (Left) or ladarixin (Right).
Panels b-e were analyzed by ANOVA with Tukey correction for multiple comparisons. Boxplots show median, interquartile range (IQR), and range. *p<0.05, **p<0.01
**Related to Fig. 5.**

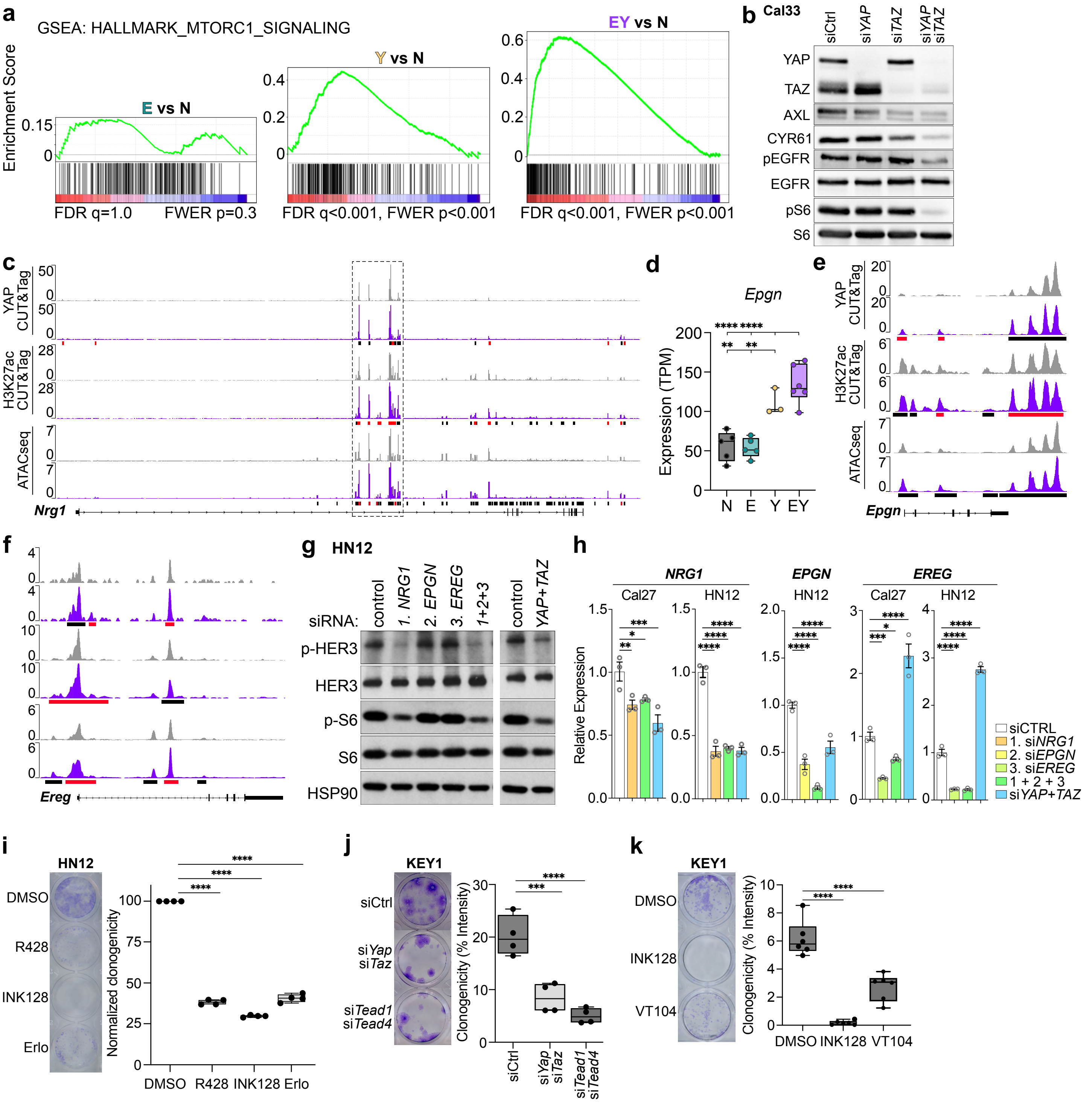
a
GSEA: HALLMARK_MTORC1_SIGNALING
Enrichment Score
E vs N
FDR q=1.0 FWER p=0.3
Y vs N
FDR q<0.001, FWER p<0.001
EY vs N
FDR q<0.001, FWER p<0.001
b
Cal33
siCtrl
si YAP
si TAZ
si YAP si TAZ
YAP
TAZ
AXL
CYR61
pEGFR
EGFR
pS6
S6
c
YAP CUT&Tag
H3K27ac CUT&Tag
ATACseq
Nrg1
d
Epgn
Expression (TPM)
N
E
Y
EY
e
YAP CUT&Tag
H3K27ac CUT&Tag
ATACseq
Epgn
f
Ereg
g
HN12
siRNA:
control
1. NRG1
2. EPGN
3. EREG
1+2+3
control
YAP+TAZ
p-HER3
HER3
p-S6
S6
HSP90
h
NRG1
Cal27
HN12
EPGN
HN12
EREG
Cal27
HN12
Relative Expression
siCTRL
1. siNRG1
2. siEPGN
3. siEREG
1 + 2 + 3
siYAP+TAZ
i
HN12
DMSO
R428
INK128
Erlo
Normalized clonogenicity
DMSO
R428
INK128
Erlo
j
KEY1
siCtrl
siYap siTaz
siTead1 siTead4
Clonogenicity (% Intensity)
siCtrl
siYap siTaz
siTead1 siTead4
k
KEY1
DMSO
INK128
VT104
Clonogenicity (% Intensity)
DMSO
INK128
VT104

**Extended Data Figure 9. YAP-mediated transcriptional activation of mTOR signaling**
(a) GSEA enrichment plots for the HALLMARK_MTORC1_SIGNALING gene set for E-, Y-, and EY-specific differentially expressed genes. Right: GSEA normalized enrichment scores and statistics for E-, Y-, and EY-specific differentially expressed genes.
(b) Western blot showing effect of siRNA-mediated knockdown of YAP, TAZ, and AXL on YAP target gene abundance and the abundance and phosphorylation of EGFR and S6 ribosomal protein in Cal33 cells.
(c) IGV tracks of YAP CUT&Tag, H3K27ac CUT&Tag, and ATACseq peaks at the *Nrg1* gene locus. Dotted line box indicates segment displayed in Fig. 6f.
(d) *Epgn* expression in transgenic tongue epithelia by RNAseq.
(e) IGV tracks of YAP CUT&Tag, H3K27ac CUT&Tag, and ATACseq peaks at the *Epgn* gene locus.
(f) IGV tracks of YAP CUT&Tag, H3K27ac CUT&Tag, and ATACseq peaks at the *Ereg* gene locus.
(g) The effects of siRNA-mediated knockdown of *NRG1*, *EREG*, *EPGN* on phospho- and total HER3, and S6 expression in HN12 whole cell lysate.
(h) The effects of siRNA-mediated knockdown of *NRG1*, *EREG*, *EPGN* on *NRG1*, *EREG*, *EPGN* transcript expression in Cal27 and HN12 cells. *EPGN* expression was undetectable in Cal27 cells (not shown).
(i) Representative wells (left) and quantification (right) of clonogenic assays in HN12 cells treated with DMSO, R428, INK128, or Erlotinib (Erlo).
(j) Representative wells (left) and quantification (right) of clonogenic assays in EY primary cultured cells cells treated with siRNA targeting, *YAP+TAZ* or *TEAD1+TEAD4*
(k) Representative wells (left) and quantification (right) of clonogenic assays in EY primary cultured cells cells treated with the mTOR inhibitor rapamycin or the YAP-TEAD inhibitor VT104.
For panels c, e , f black bars indicate significant peaks. Red bars indicate EY-gained peaks.
For panels d, i, j, k boxplots show median, interquartile range (IQR), and range; ANOVA with Tukey correction for multiple comparisons. *$p<0.05$, **$p<0.01$, ***$p<0.001$, ****$p<0.0001$.
**Related to Fig. 6.**

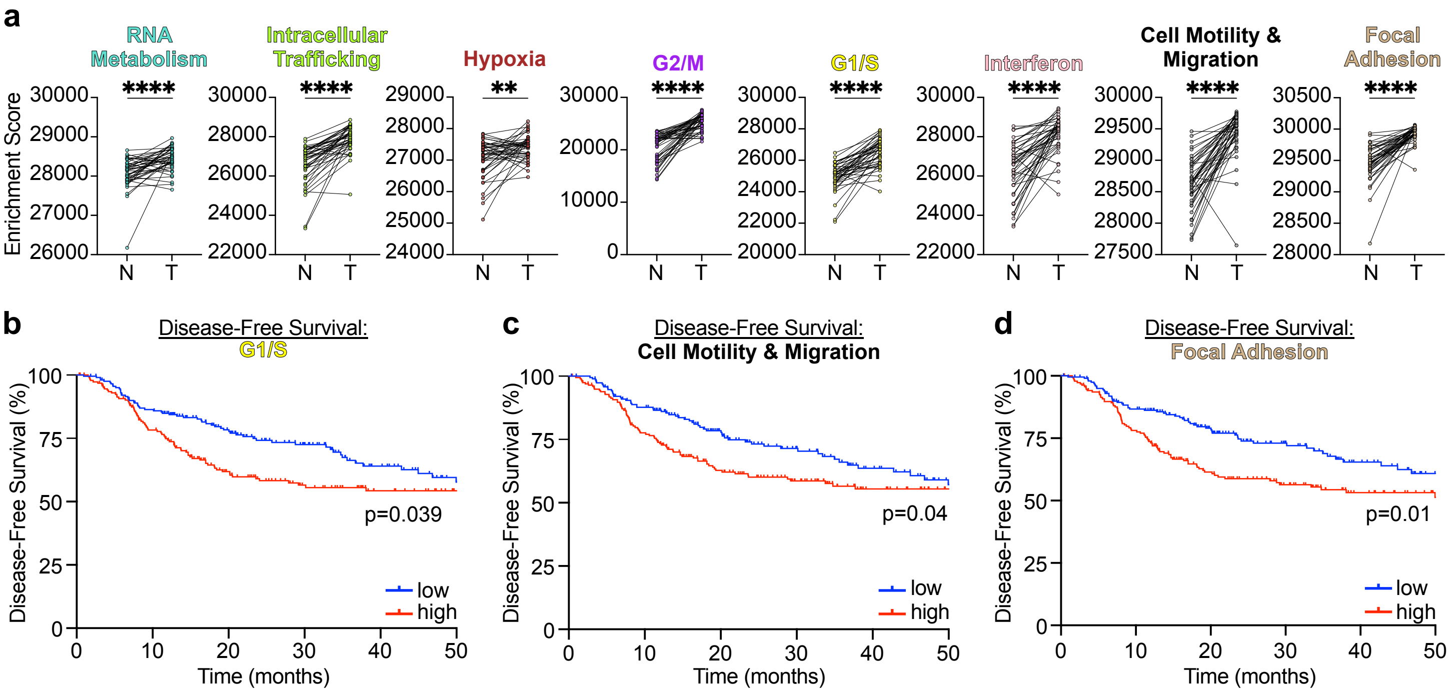
a
RNA Metabolism
Intracellular Trafficking
Hypoxia
G2/M
G1/S
Interferon
Cell Motility & Migration
Focal Adhesion
****
**
Enrichment Score
N
T
b
Disease-Free Survival: G1/S
p=0.039
c
Disease-Free Survival: Cell Motility & Migration
p=0.04
d
Disease-Free Survival: Focal Adhesion
p=0.01
Disease-Free Survival (%)
Time (months)
low
high

**Extended Data Figure 10. TIC programs are enriched in HNSC and associated with disease-free survival**

(a) Related to Figure 7i. EY-module enrichment in malignant tumors (T) compared to matched normal solid tissues (N) by single sample GSEA among subjects in The Cancer Genome Atlas (TCGA) Head and Neck Squamous Carcinoma (HNSC) cohort (n=43 subjects with matched T and N samples). Two-tailed paired T-test: **p<0.01, ****p<0.0001.

(b-d) Kaplan-Meier plots demonstrating worse disease-free survival (n=393) among TCGA-HNSC subjects with greater median enrichment for the (b) G1/S, (c) pEMT, and (d) Focal Adhesion EY-modules.

**Related to Fig. 7.**
